# Kineochelins - a novel group of siderophores from an Antarctic bacterium

**DOI:** 10.64898/2026.02.23.707395

**Authors:** Stanislava Kralova, Peter Spacek, Johannes Gafriller, Matej Bezdicek, Viktoria Medvedcova, Joana Séneca Cardoso Da Silva, Jay Osvatic, Ulrike Grienke, Thomas Rattei, Olga N. Sekurova, Sergey B. Zotchev, Martin Zehl, Alexander Loy

## Abstract

**Background:** The global rise of antimicrobial resistance has intensified the search for new microbial metabolites from underexplored environments and taxonomic groups. Extreme and geographically isolated habitats, such as Antarctic terrestrial ecosystems, represent promising reservoirs of novel biosynthetic diversity, particularly among rare and difficult-to-cultivate actinomycetes, where chemical mediators are thought to play key roles in microbial persistence and interaction under resource-limited conditions.

**Results:** Here, we report the characterization of kineochelins, a previously undescribed group of siderophores produced by an Antarctic isolate, *Actinokineospora* sp. UV203 representing difficult to cultivate actinomycetes. Structural elucidation revealed a set of closely related congener molecules with a mixed-ligand architecture consistent with metal-chelating activity. Genome mining combined with transcriptomic analysis identified the involvement of a dedicated nonribosomal peptide synthetase-encoding biosynthetic gene cluster responsible for kineochelin production. Comparative genomic analyses showed that, although kineochelin biosynthetic genes share limited homology with those of known mixed-ligand siderophores, their biosynthetic pathways differ substantially in gene content and organization, indicating a distinct evolutionary lineage. Functional characterization of kineochelins demonstrated strong and selective iron chelation, with pronounced affinity for ferric and ferrous iron. Crude culture extracts inhibited the growth of bacterial strains isolated from the same Antarctic environment, suggesting that kineochelin-associated chemistry contributes to iron-mediated competitive interactions within native microbial communities. In addition, kineochelin-enriched fractions exhibited selective inhibitory activity against the opportunistic yeast pathogen *Nakaseomyces glabratus* and a clinical isolate of *Saccharomyces cerevisiae* associated with invasive infection.

**Conclusions:** Together, these findings expand the known chemical and biosynthetic diversity of the genus Actinokineospora and demonstrate that Antarctic rare actinomycetes are a valuable source of novel natural products with potential relevance for microbial ecology and biotechnology. The ecological activities of kineochelins highlight the role of iron acquisition in shaping microbial interactions in extreme environments and underscore the biotechnological potential of metabolites derived from underexplored polar microorganisms.

## Introduction

Antimicrobial resistance is an escalating global crisis that threatens to erode decades of medical progress, already causing millions of deaths annually[1]. The alarming spread of multidrug-resistant microorganisms, “superbugs”, has created an urgent need for new classes of antimicrobials with novel modes of action[2,3]. Yet, the antibiotic discovery pipeline has been in steady decline, with most approved drugs representing derivatives of existing scaffolds rather than structurally innovative compounds[4]. Strikingly, of the 13 new antimicrobial drugs approved between 2017 and 2023, only two met the World Health Organization’s innovation criteria[2,3]. To overcome this stagnation, efforts have also explored non–natural poduct strategies such as bacteriophages, nanoparticles, and monoclonal antibodies[5,6]. Nevertheless, the search for structurally novel natural products (NPs) remains central, with increasing focus on underexplored habitats that may harbour organisms with distinctive biosynthetic repertoires and provide access to untapped chemical space[7–9].

Extreme and geographically isolated environments have emerged as important reservoirs of microbial diversity with a high potential for the discovery of novel NPs[10]. In such habitats, strong selective pressures, including temperature extremes, nutrient limitation, desiccation, and radiation, favour microorganisms with specialized metabolic and biosynthetic capabilities, often resulting in chemically distinctive specialised (secondary) metabolites, commonly referred to as NPs[11]. Antarctic terrestrial ecosystems represent one of the most extreme examples of these conditions, combining chronic low temperatures with recurrent freeze–thaw events, seasonal fluctuations in light and nutrients, and long-term ecological isolation[12–14]. These constraints have shaped highly adapted microbial communities in which SMs play central roles in resource acquisition, stress tolerance, and interspecies interactions[12,15–17]. Antarctic microbiota have attracted increasing attention as a source of structurally and functionally diverse NPs with biomedical and biotechnological relevance[18–21]. However, despite extensive cultivation-independent surveys revealing a rich diversity of biosynthetic gene clusters (BGCs) in Antarctic microbes, relatively few of these predicted pathways have been linked to purified metabolites or experimentally validated biosynthetic systems[12]. This disconnection between genomic potential and chemical characterization highlights the need for integrated approaches that couple cultivation, metabolite isolation, and genomic analysis to link biosynthetic capacity with expressed chemistry and microbial interactions in Antarctic soil communities.

In this study, we established a bacterial culture collection from Antarctic soils sampled at the James Ross Island (2007-2013) that consist of strains of diverse phylogenetic origin and with broad genomic potential for SM production. Using a selected strain, that represents a potentially novel *Actinokineospora* species, we show how this resource can be exploited for the discovery of previously unknown secondary metabolites. The BGC repertoire and liquid chromatography–mass spectrometry (LC–MS)-based SM analysis of *Actinokineospora* sp. UV203 guided the isolation and structure elucidation of a new group of mixed-ligand siderophores with antimicrobial properties, which we named kineochelins. Siderophores belong to the broader family of metallophores, molecules that mediate microbial competition through metal acquisition[22]. In the context of antimicrobial resistance, metallophores may be used to deprive pathogens of essential metals or exploiting their metal transport systems for targeted delivery of antimicrobials[23–26]. Beyond clinical applications, microbial metallophores are increasingly recognized as versatile tools in biotechnology, with emerging roles in bioremediation, metal recovery, biosensing, and the modulation of microbial communities[27–29].

## Results

### Isolation, taxonomic placement, and biosynthetic gene clusters of Actinokineospora sp. UV203

As part of a long-term biodiversity campaign on James Ross Island, Antarctica (since 2007), we established a collection of 972 morphologically and phylogenetically diverse bacterial strains that were isolated from the active layer of soils above permafrost. Preliminary classification based on 16S rRNA gene sequencing and phylogenetic analysis assigned most of these isolates to four phyla: Actinomycetota, Bacillota, Bacteroidota, and Pseudomonadota (Figure 1A). A subset of phylogenetically divergent strains (n=145), prioritized based on 16S rRNA gene sequence dissimilarity and unique phylogenetic branching patterns, was selected for whole-genome sequencing. Most genome-sequenced strains, including strain UV203, represent previously undescribed species or higher taxa. Strain UV203 was isolated as a colony on starch-casein agar from a pretreated active layer soil sample collected during the austral summer of 2022 at Abernethy Flats. Phylogenomic analysis suggested that strain UV203 may represent a new species within the *Actinokineospora* genus. This classification was supported by genome distance metrics, including average nucleotide identity (ANI: 93.6%) and digital DNA–DNA hybridization (dDDH: 51.8%) values to the closest related species *Actinokineospora alba* DSM 45114^T^, both falling below the accepted thresholds for species delineation[30]. Core genes within the up-to-date bacterial core gene (UBCG) pipeline[31] further supported the placement of UV203 as a distinct species-level lineage (Figure 1B). Although genome sequences are not yet available for three validly described *Actinokineospora* species (*Actinokineospora riparia*, *Actinokineospora cibodasensis*, and *Actinokineospora acnipugnans*), 16S rRNA gene phylogeny indicated that strain UV203 does not cluster with these taxa either (Supplementary Fig. S1).

**Figure 1.**
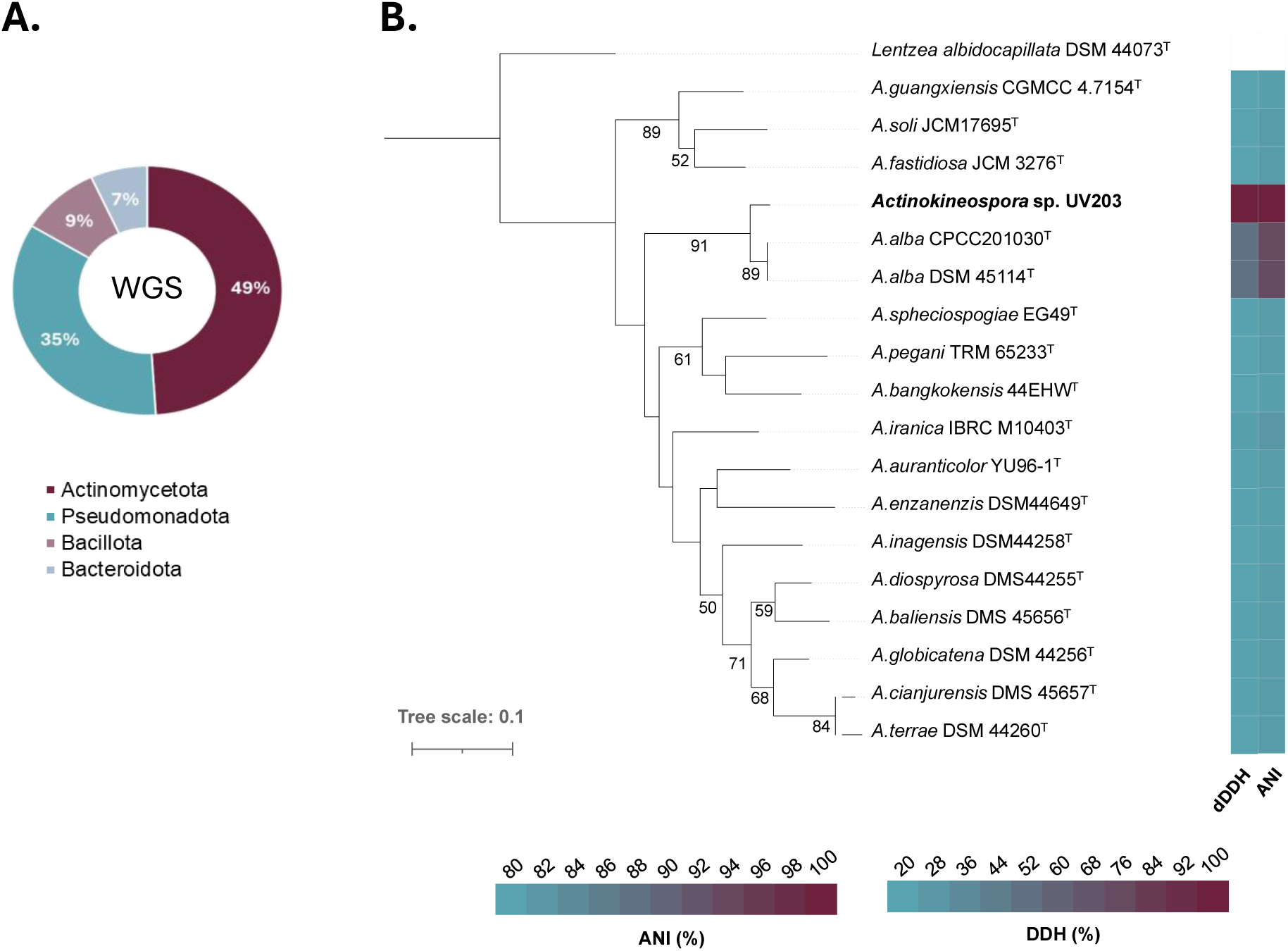
Phylogeny of strain UV203 suggests it represents a yet undescribed *Actinokineospora* species in the Actinomycetota. (A) Phylum-level composition of the isolates selected for whole-genome sequencing (n = 145), based on genome-derived taxonomic assignments. (B) Genome-scale up-to-date bacterial core gene (UBCG) phylogeny placed strain UV203 as a distinct lineage within *Actinokineospora*; scale bar indicates 0.1 substitutions per site. The numbers at the nodes indicate the gene support index (maximum value, 91)[31]. Pairwise genomic relatedness between UV203 and type strains of validly published *Actinokineospora* species with available genomes is shown as heatmaps of Average Nucleotide Identity (ANI, %; FastANI) and digital DNA–DNA hybridization (dDDH, %; GGDC)[32,33]. Colour scales shown below.

To further explore the biosynthetic potential of *Actinokineospora* sp. UV203, we combined genome-based prediction of BGCs and metabolomic analysis using an untargeted LC-MS workflow. antiSMASH predicted 21 BGCs, many without close homologs in public databases (Table 1). Manual inspection of the antiSMASH output indicated that this number is likely an underestimate, as closely spaced or partially overlapping BGCs are not always fully resolved by automated prediction algorithms, a known limitation of automated BGC boundary prediction[34]. The production of >28 SMs was detected under various tested culturing conditions, whereby many of them could be grouped together as congeners either by tentative identification or because the MS/MS spectra strongly indicated shared substructures (Supplementary Table S1). In NP research, the term congeners usually refers to a group of compounds produced by the same BGC, thus sharing the same core scaffold but differing in size, specific substituents, or functional groups[35]. Next to several SMs that were tentatively identified, such as gaburedins A-D and F[36], a large number of them yielded MS and MS/MS spectra that could not be matched to any known NP present in GNPS[37], NPAtlas[38], or the CAS database. We thus attempted the isolation and characterization of some of these potentially novel NPs.

We initially focused on a pair of isomeric compounds detected as [M+H]^+^ ions at *m/z* 575.0587±0.0029 and with the proposed sum formula of C_22_H_24_Cl_2_N_4_O_6_S_2_. They stood out out by the presence of two Cl-atoms as inferred from the isotopic pattern and two sulphur atoms (Supplementary Table S1). BGC 2.18 is the only cluster in the UV203 genome encoding a halogenase, specifically a tryptophane-halogenase. In addition, it contains at least two non-ribosomal peptide synthetase (NRPS)-modules both predicted to incorporate a cysteine, the sulphur-containing amino acid. Despite partial similarity to the coelibactin BGC detected by antiSMASH, the detected chlorinated metabolites are inconsistent with coelibactin[39], and no other known product could be confidently assigned to BGC 2.18.

**Table 1.**
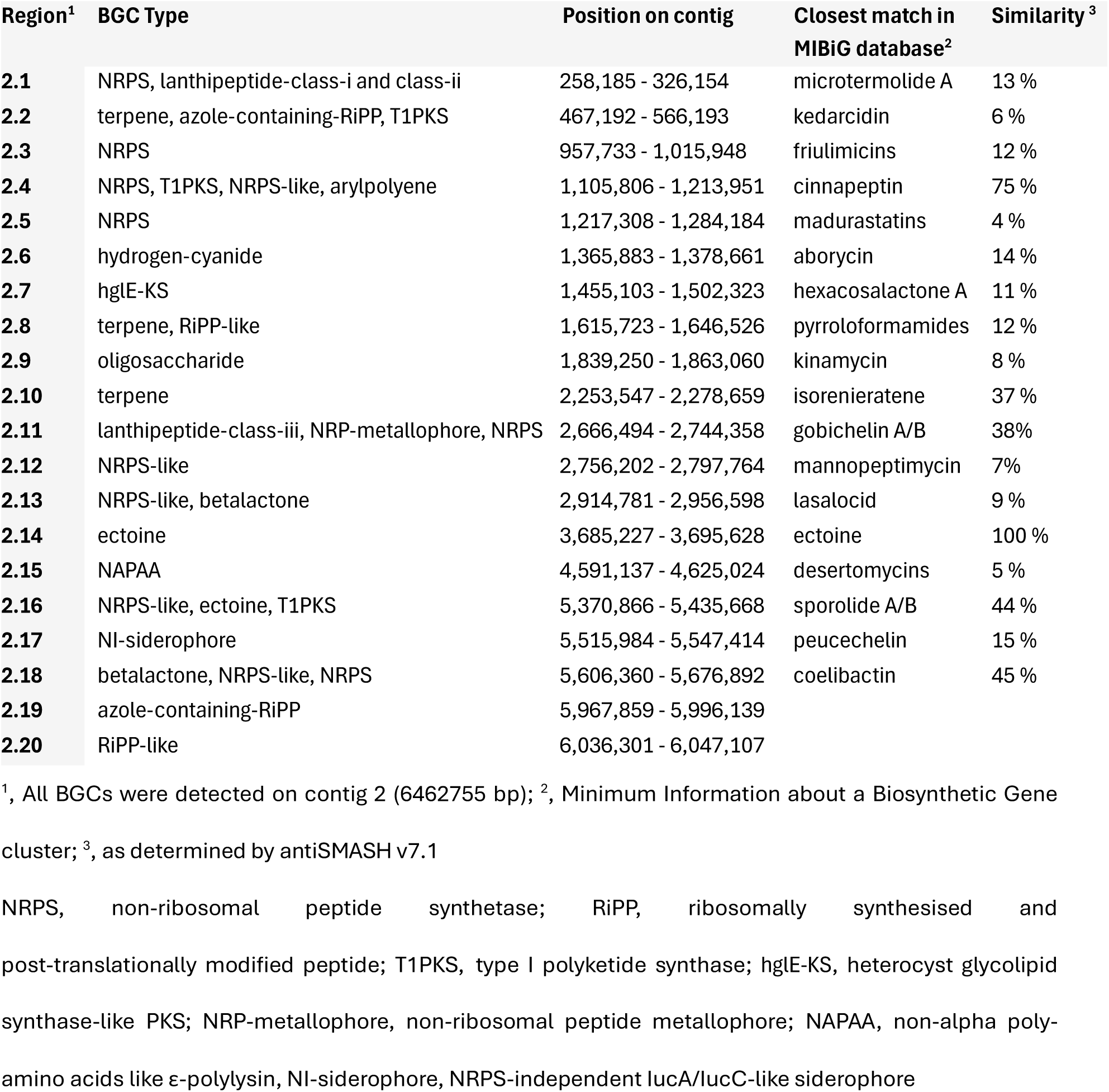
Tabulated view of all detected BGCs in the UV203 genome with antiSMASH v7.1.

Since the wild-type strain only produced very low amounts of these two isomers under various cultivation conditions, we aimed for targeted overexpression. To this end, we first used a pSET152 integrative vector carrying the constitutive *ermE*\*p promoter[40] together with two pathway-specific regulators from BGC 2.18 (LuxR and HxlR-like). While particularly the overexpression of LuxR seemed to have a positive effect, the production was still far too low to attempt isolation, likely due to host-specific promoter inefficiency[41,42]. As a second strategy, we replaced the *ermE*\*p promoter in the vectors with the promoter from the UV203 *rrn* operon, which is considered highly active based on its strong transcriptional output in bacteria[43,44]. However, while again showing positive effects compared to the empty vector control, this modification also failed to sufficiently raise the production of the target metabolites for purification. This indicates that additional regulatory or physiological constraints are in place in the native host.

In an unanticipated outcome, one of the resulting mutants exhibited a complete loss of antimicrobial activity (Supplementary Fig. S2). Comparative metabolomic analysis revealed the absence of a large group of structurally related congeners in the extract from the non-active mutant that covered a wide range of monoisotopic masses from 219 to 722 Da (Supplementary Fig. S2 and Supplementary Table S1). The three smallest congeners produced by the wild type were tentatively identified as asteroidic acid[45], pseudomobactin A[46], and vulnibactin 2[47]. These compounds are small siderophores biosynthesized from salicylic acid and threonine. The congeners with higher masses shared with them either the 2-(2-hydroxyphenyl)-5-methyl-4,5-dihydrooxazole-4-carbonyl or 2-(2-hydroxyphenyl)-5-methyl-4,5-oxazole-4-carbonyl subunit, but differed in mass by increments matching to amino acids such as glycine, serine, and *N*-hydroxy-ornithine derivatives, the latter typically found in hydroxamate siderophores (Figure 2). The tentative structures derived from the LC–MS data did not match any known compounds in available databases. We provisionally designated this metabolite group as kineochelins and named the individual members based on the tentative structures derived from the MS/MS spectra, with the letters A-E referring to the number of amino acids attached to the vulnibactin 2 core from 5 to 1, respectively, and the subscript index 1 or 2 referring to the oxidation sate of the oxazole-ring (Figure 2).

In conclusion, while the engineered overproduction of the intended metabolite from BGC 2.18 was unsuccessful, this functional-genomics strategy unexpectedly led to the discovery of a structurally unique and likely biologically active group of NPs.

**Figure 2.**
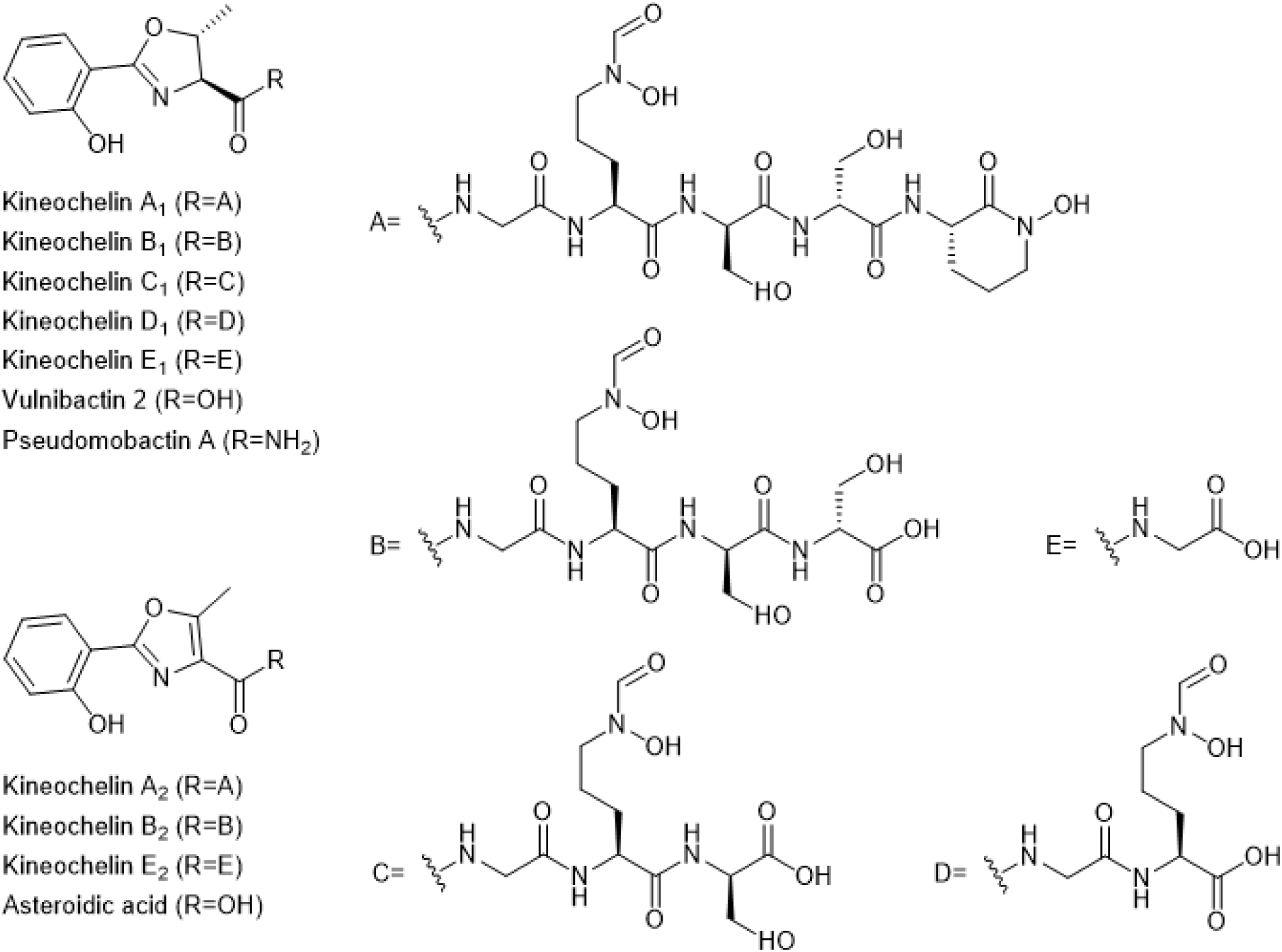
Proposed naming and structures of the kineochelin group and the known shunt products. Shown are kineochelins and known shunt products vulnibactin 2, pseudomobactin A, and asteroidic acid. Only the structures of kineochelin A1 and E1 were elucidated by comprehensive NMR and Marfey’s analysis in this study, while for all other congeners they remain tentative.

### Structural analysis of kineochelins

The kineochelins were enriched from the SM17 medium supernatant by adsorption to HP-20 resin, from which they were extracted with *n*-butanol. The dried extract was then fractionated by flash chromatography and pooled into 21 fractions according to results from analysis with ultra-high-performance liquid chromatography coupled to photodiode-array and evaporative light-scattering detection (UHPLC-PDA-ELSD).

Analysis by UHPLC-ELSD as well as by UHPLC-MS demonstrated fraction 17 (2.0 mg) to contain kineochelin E_1_ (**1**) of high purity (Supplementary Fig. 3). This compound, which was assigned the molecular formula C_13_H_14_N_2_O_5_ (Supplementary Table S1), was subsequently analyzed by 1D and 2D NMR spectroscopy. Signals of 13 carbon and 11 non-exchanging hydrogen atoms were detected, while three fast-exchanging hydrogens could not be observed in MeOH-d_4_ (Table 2). The ^1^H signals belonged to three spin systems, one made up by four aromatic methines (H-2 to H-5), the second one consisting of two aliphatic methines (H-9 and H-10) and a methyl group (H-8), and finally an isolated methylene group (H-12a and H-12b). A thorough interpretation of the 1D and 2D NMR spectra (Figure 3 and Supplementary Fig. S4-8) confirmed that the former two spin systems are indeed characteristic for the 2-(2-hydroxyphenyl)-5-methyl-4,5-dihydrooxazole-4-carbonyl substructure that was expected from the LC-MS data. The observed chemical shifts of C-1 to C-11 and H-2 to H-10 as well as the measured coupling constants of the latter match very well to the previously reported data of pseudomobactin A[46] (Table 2), amychelin C[48], and the catenulobactins A and B[49]. The heteronuclear multiple-bond correlations (HMBC) of the methylene hydrogens (H-12a and H-12b) to the carbonyl C-11 proof that a glycine is attached to the C-terminus of the oxazoline moiety via an amide bond.

Although the excellent agreement of the NMR data with those of pseudomobactin A (Table 2) suggests that kineochelin E_1_ is also derived from L-Thr, the configuration around the two chiral centers C-9 and C-10 could not be established with certainty from the NMR data. Furthermore, a second set of signals was observed in the NMR data in a ratio of approximately 3:5 to the one described above (Supplementary Table S2). This set of signals was hypothesized to belong to a stereoisomer of **1** obtained by configurational inversion at one of the chiral centers. The absolute configuration was determined by Marfey’s analysis after hydrolysis of 410 μg of kineochelin E_1_[50]. Comparison with all four possible stereoisomers L-Thr, D-Thr, D-allo-Thr, and L-allo-Thr by LC-MS proofed that kineochelin E_1_ is indeed derived from L-Thr and that there is no other stereoisomer present in the sample (Supplementary Fig. S9). Kineochelin E_1_ is thus characterized as the new compound {[(4*S*,5*R*)-2-(2-hydroxyphenyl)-5-methyl-4,5-dihydro-1,3-oxazole-4-carbonyl]amino}acetic acid (Figure 3).

**Table 2.** ^1^H (600 MHz) and ^13^C NMR data (151 MHz) of kineochelin E1 (**1**) and kineochelin A1 (**2**) in CD3OD in comparison with literature data for pseudomobactin A (*δ* in ppm).

|  | Kineochelin E<sub>1</sub> (**1**) |  | Kineochelin A<sub>1</sub> (**2**) |  | Pseudomobactin A<sup>a</sup> |  |
| --- | --- | --- | --- | --- | --- | --- |
| Position | $\delta_{\text{H}}$ (J in Hz) | $\delta_{\text{C}}$, type | $\delta_{\text{H}}$ (J in Hz) | $\delta_{\text{C}}$, type | $\delta_{\text{H}}$ (J in Hz) | $\delta_{\text{C}}$, type |
| 1 | — | 161.3, C | — | 161.2, C | — | 161.1, C |
| 2 | 6.96, m, ov | 117.9, CH | 6.95, m, br, ov | 117.9, CH | 6.96, d (8.3) | 117.7, CH |
| 3 | 7.41, m | 135.2, CH | 7.41, t (7.8), br | 135.2, CH | 7.41, td (8.3, 1.7) | 135.0, CH |
| 4 | 6.90, m, ov | 120.1, CH | 6.90, m, br, ov | 120.1, CH | 6.90, td (7.4, 1.7) | 119.9, CH |
| 5 | 7.69, dd (7.9, 1.7) | 129.7, CH | 7.68, m, br | 129.6, CH | 7.68, dd (7.4, 1.7) | 129.5, CH |
| 6 | — | 111.8, C | — | 111.8, C | — | 111.6, C |
| 7 | — | 168.3, C | — | 168.0, C | — | 167.8, C |
| 8 | 1.57, d (6.3) | 21.6, CH<sub>3</sub> | 1.56, m, ov | 21.6, CH<sub>3</sub> | 1.57, d (6.3) | 21.4, CH<sub>3</sub> |
| 9 | 4.91, dq (7.5, 6.3) | 80.8, CH | 4.92, m, br | 80.5, CH | 4.90, qd (7.3, 6.3) | 80.6, CH |
| 10 | 4.51, d (7.5) | 75.9, CH | 4.55, d (7.5), br, ov | 75.9, CH | 4.46, d (7.3) | 75.5, CH |
| 11 | — | 173.3, C | — | 173.9, C | — | 175.6, C |
| 12a | 3.93, d (17.0), ov | 43.0, CH<sub>2</sub> | 3.83-4.16, m, ov | 43.4, CH<sub>2</sub> | — | — |
| 12b | 3.87, d (17.5) |  |  |  | — | — |
| 13 | — | 173.8<sup>b</sup>, C | — | 173.0, C | — | — |
| 14 | — | — | 8.28, s, br | 159.6, CH | — | — |
| 15 | — | — | 3.53, m, ov | 50.9, CH<sub>2</sub> | — | — |
| 16 | — | — | 1.71<sup>c</sup>, m, ov | 24.1, CH<sub>2</sub> | — | — |
| 17a | — | — | 1.80<sup>c</sup>, m, ov | 28.8, CH<sub>2</sub>, ov | — | — |
| 17b | — | — | 1.71<sup>c</sup>, m, ov |  | — | — |
| 18 | — | — | 4.34, m, ov | 55.2, CH | — | — |
| 19 | — | — | — | 175.7, C | — | — |
| 20a | — | — | 3.92<sup>c</sup>, m, ov | 62.3, CH<sub>2</sub> | — | — |
| 20b | — | — | 3.81-3.89<sup>c</sup>, m, ov |  | — | — |
| 21 | — | — | 4.27, m | 58.6, CH | — | — |
| 22 | — | — | — | 172.4, C | — | — |
| 23a | — | — | 4.00, m, ov | 62.8, CH<sub>2</sub> | — | — |
| 23b | — | — | 3.81-3.89<sup>c</sup>, m, ov |  | — | — |
| 24 | — | — | 4.41, m | 58.1, CH | — | — |
| 25 | — | — | — | 172.1, C | — | — |
| 26a | — | — | 3.63, m, ov | 52.6, CH<sub>2</sub> | — | — |
| 26b | — | — | 3.59, m, ov |  | — | — |
| 27a | — | — | 2.04, m, ov | 21.5, CH<sub>2</sub> | — | — |
| 27b | — | — | 1.94, m, ov |  | — | — |
| 28a | — | — | 2.02<sup>c</sup>, m, ov | 28.8, CH<sub>2</sub>, ov | — | — |
| 28b | — | — | 1.80<sup>c</sup>, m, ov |  | — | — |
| 29 | — | — | 4.59, dd (9.8, 5.2) | 51.5, CH | — | — |
| 30 | — | — | — | 167.0, C | — | — |

<sup>a</sup> Values obtained from Oluwabusola *et al.* (2021)[50]
<sup>b</sup> Value determined from HMBC spectrum
<sup>c</sup> Value determined from HSQC spectrum

**Figure 3.**
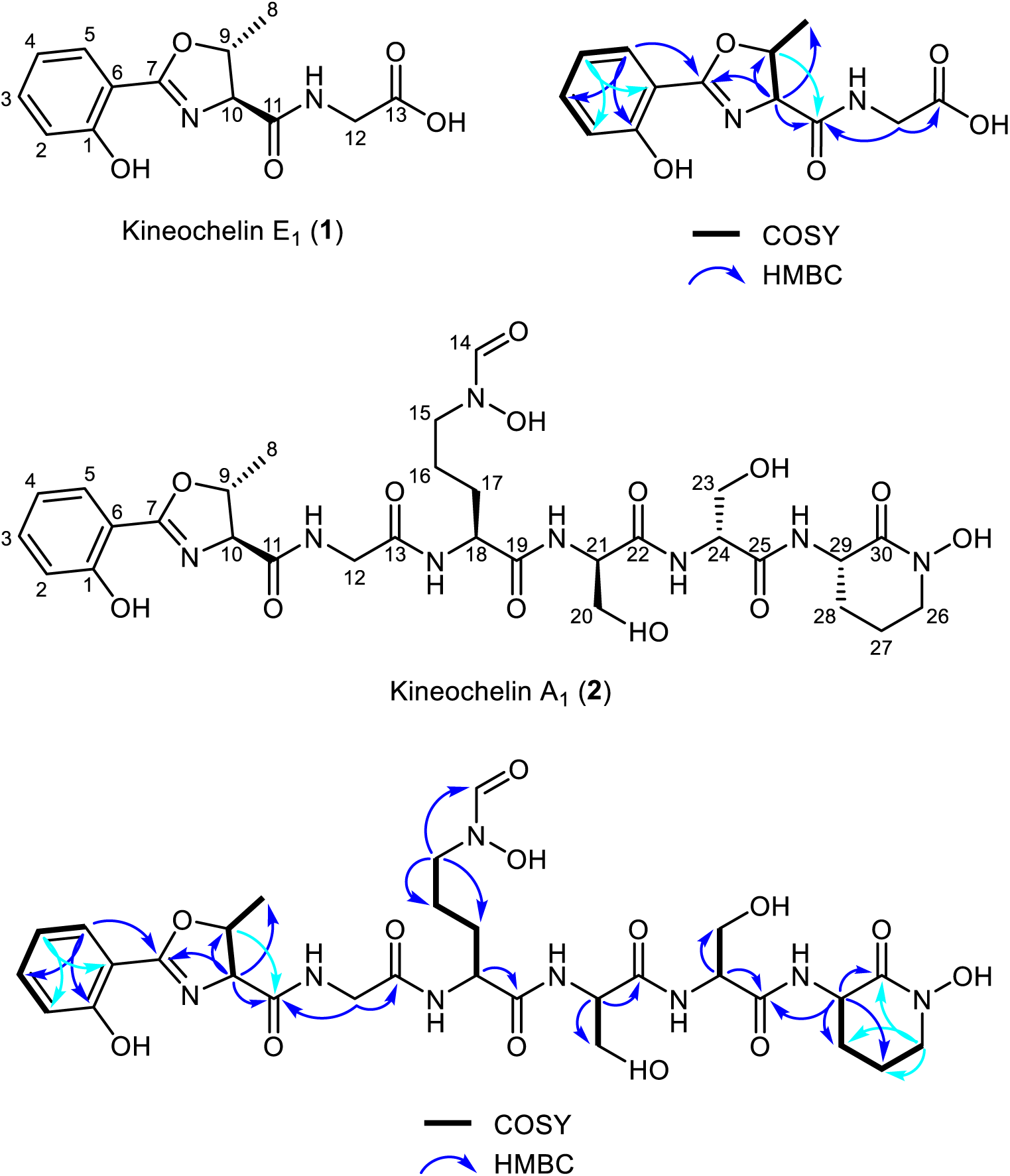
Structure of kineochelin E1 (1) and kineochelin A1 (2). Atom numbering and ^1^H-^1^H COSY and key ^1^H-^13^C HMBC correlations are shown.

Fraction 11 (5.8 mg) contained kineochelin A_1_ (**2**), the largest and most abundant of the detected siderophores, with an estimated purity of ∼80% according to the ELSD data (Supplementary Fig. S10). Kineochelin A_1_ has a sum formula of C_30_H_42_N_8_O_13_ (Supplementary Table S1) and was hypothesized to contain kineochelin E_1_ (**1**) as N-terminal substructure. This was confirmed by comparison of the NMR data of fraction 11 (Supplementary Fig. S11-S15) with those of fraction 17. The signals of the 2-(2-hydroxyphenyl)-5-methyl-4,5-dihydro-1,3-oxazole-4-carbonyl substructure of **2** were nearly identical to those in **1** (Table 2). The signals of the adjacent glycine were slightly more affected by the chain extension, but also easily assigned, despite considerable overlap of various signals in the much more complex ^1^H NMR spectrum of **2**. Based on the thorough interpretation of the MS/MS spectra of all detected kineochelins (Supplementary Fig. S16-S26), the following units were sequentially attached to kineochelin E_1_ to give kineochelin D_1_ to A_1_, respectively: C_6_H_10_N_2_O_3_, C_3_H_5_NO_2_, C_3_H_5_NO_2_, and C_5_H_8_N_2_O. The third amino acid after L-Thr and Gly was found to be *N*⁵-hydroxy-*N*⁵-formyl-ornithine, a non-proteinogenic amino acid typical for hydroxamate siderophores. The identification of this amino acid was supported by the characteristic signals of the formyl-group (H-14 and C-14) in the 1D and 2D NMR spectra. The next two units were found to be Ser residues, followed by a cyclized *N*⁵-hydroxy-ornithine as final C-terminal amino acid residue. The latter could either be piperazic acid, as in the cahuitamycins[51], or result from amide bond formation between the C-terminus and the side-chain hydroxylamine group, as in the amychelins and gobichelins[52,53]. An HMBC cross signal of H-26 to the carbonyl C-30 clearly proofs the cyclic amide (Figure 3), which leads to the complete planar structure. To establish the stereochemistry, 750 μg of **2** were hydrolysed for Marfey’s analysis and the reaction products compared with commercial standards. Only L-Thr, L-Orn, and D-Ser were detected, thus fully establishing the absolute configuration of kineochelin A_1_ (Supplementary Fig. S27).

Along with the kineochelins, two 2,5-diketopiperazines were isolated from fractions 2 and 3 (4.6 mg) and fraction 5 (1.2 mg). They were identified as cyclo(Hyp-Leu) and cyclo(Hyp-Phe), respectively, by interpretation of the HRESIMS and NMR data and by comparison to literature values (Supplementary Fig. S28-S36 and Supplementary Tables S1 and S3)[54,55]. However, they were not further studied regarding stereochemistry or biological activity.

### Biosynthetic gene cluster for kineochelins production

While several BGCs in the UV203 genome contain NRPS modules, only BGC 2.11 could be matched to the proposed biosynthesis of the kineochelins (Table 1). It harbours three core genes, *kinA, kinB,* and *kinC,* encoding a total six of NRP modules, two of which contain epimerization-domains (Figure 4). Furthermore, it encodes a salicylate synthase (*kinL*) and displays ∼38% homology to the *gob* BGC that is responsible for the biosynthesis of the structurally related mixed-ligand siderophores gobichelin A and B[53]. The cluster is hereafter referred to as the *kin* cluster.

In the proposed biosynthesis, the aromatic starter unit is derived from chorismate, which is converted into salicylate via a standalone salicylate synthase KinL, homologous to Ccb3 from the celestinticine biosynthetic pathway (59% identity)[56] and AmcL from *Streptomyces* sp. AA4 (49.4% identity) in the amychelin BGC[52] (Supplementary Table S4). The resulting salicylate is activated by the salicylate-AMP ligase KinH, which shares 58.5% identity with AmcH from the amychelin cluster. In contrast to GobJ, AmcG, and CahA, the initiating NRPSs in the biosyntheses of the gobichelins, amychelins, and cahuitamycins, respectively, no aryl carrier protein (ArCP) domain was detected in the corresponding NRPS KinA. We assume that KinH instead loads the activated salicylate to the ArCP domain encoded in the didomain protein KinT. KinA activates and loads L-Thr onto its peptidyl carrier protein (PCP) domain. Its N-terminal cyclization (Cy) domain accepts KinH as the salicylate donor and catalyses the cyclodehydration with L-Thr to give the initiating 2-(2-hydroxyphenyl)-5-methyl-4,5-dihydrooxazole-4-carbonyl unit. This intermediate is then passed to the downstream three-module NRPS KinB (Figure 4B), which extends the chain by a Gly, an *N*⁵-hydroxy-*N*⁵-formyl-L-ornithine, and a D-Ser residue, the latter via epimerization of L-Ser. The growing chain is than passed to the two-module NRPS KinC, which adds another D-Ser and an *N*⁵-hydroxy-L-ornithine (*N*-OH-L-Orn) residue. Like in the amychelin biosynthetic pathway, the final NRPS module in the kineochelin assembly lacks an associated *C*-terminal thioesterase domain[52]. The final NRPS module lacks a C-terminal thioesterase domain, whereas a standalone ArCP–TE didomain protein (KinT) is encoded within the cluster. Structural analysis of kineochelins revealed the presence of a cyclic amide at the C-terminus.

**Figure 4.**
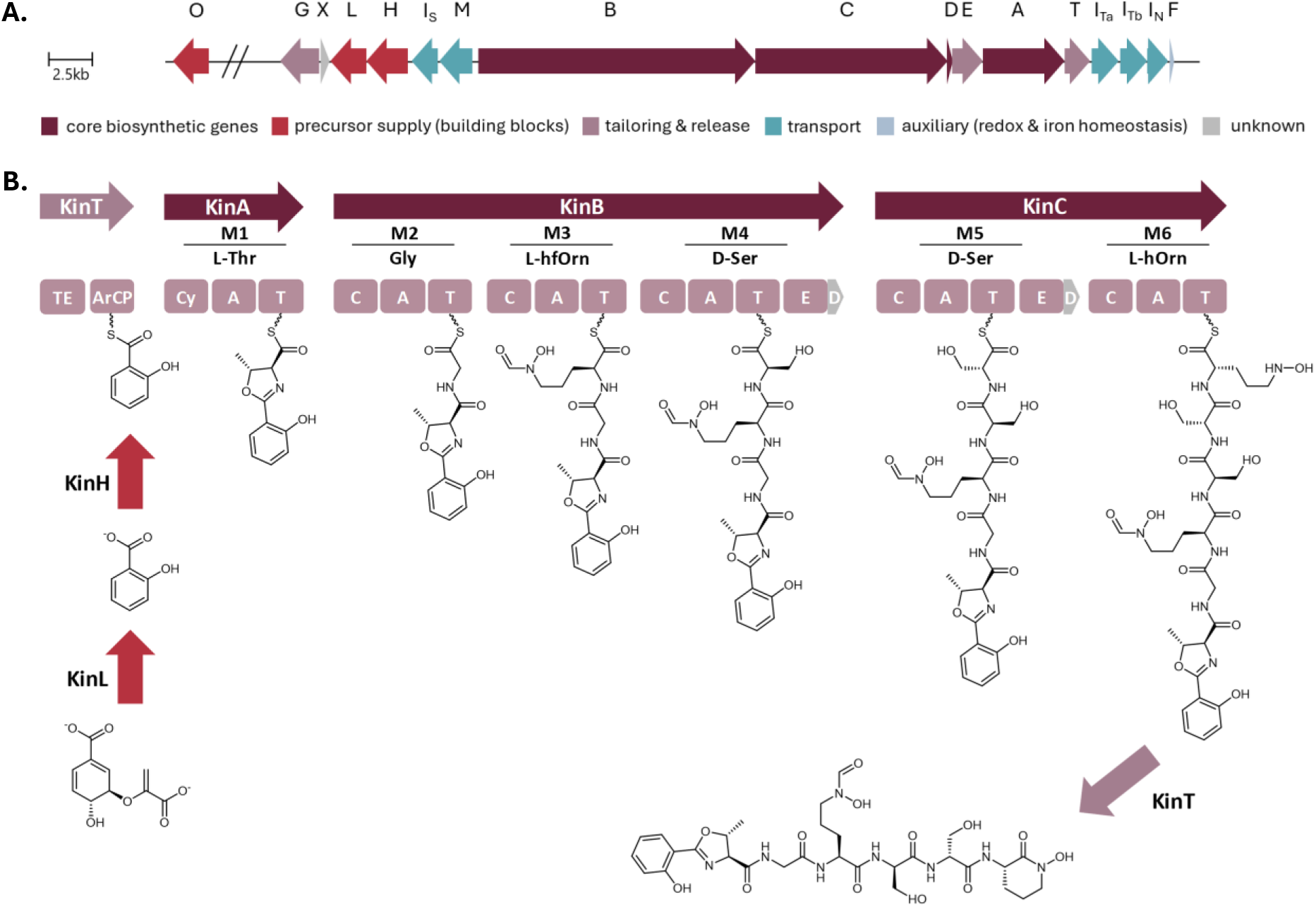
Gene cluster and proposed model for kineochelin biosynthesis. (A.) Kineochelin biosynthetic genes cluster (*kin*) with genes kinA-kinX. Gene colour-coding is indicated in the legend. (B) Proposed biosynthetic model for the kineochelins. Domain abbreviations: C, condensation; A, adenylation; T, thiolation domain (peptidyl carrier protein); E, epimerization; TE, thioesterase (product-release by macrocyclization or hydrolysis); D, docking domain; ArCP, aryl carrier protein.

Comparative network analysis with BiG-SCAPE placed the *kin* cluster of strain UV203 in a separate biosynthetic gene cluster family (GCF) with two uncharacterized BGCs from *Actinokineospora alba* strains (Fig. 5A, 5B). To further contextualize the *kin* cluster, we manually included additional BGCs whose core biosynthetic genes showed the highest similarity to *kin* genes, as identified by NCBI protein BLAST searches and antiSMASH analysis (Supplementary Tables S5, S6). Clinker-based synteny comparisons revealed that full-length (global) alignments of core biosynthetic proteins between the *kin* cluster and related clusters did not exceed 40% amino acid identity[57] (Figure 5C). In contrast, local BLAST-based alignments identified >40% identity for several homologous proteins[58], indicating detectable gene-level similarity restricted to specific domains or regions[59] (Supplementary Table S4).

**Figure 5.**
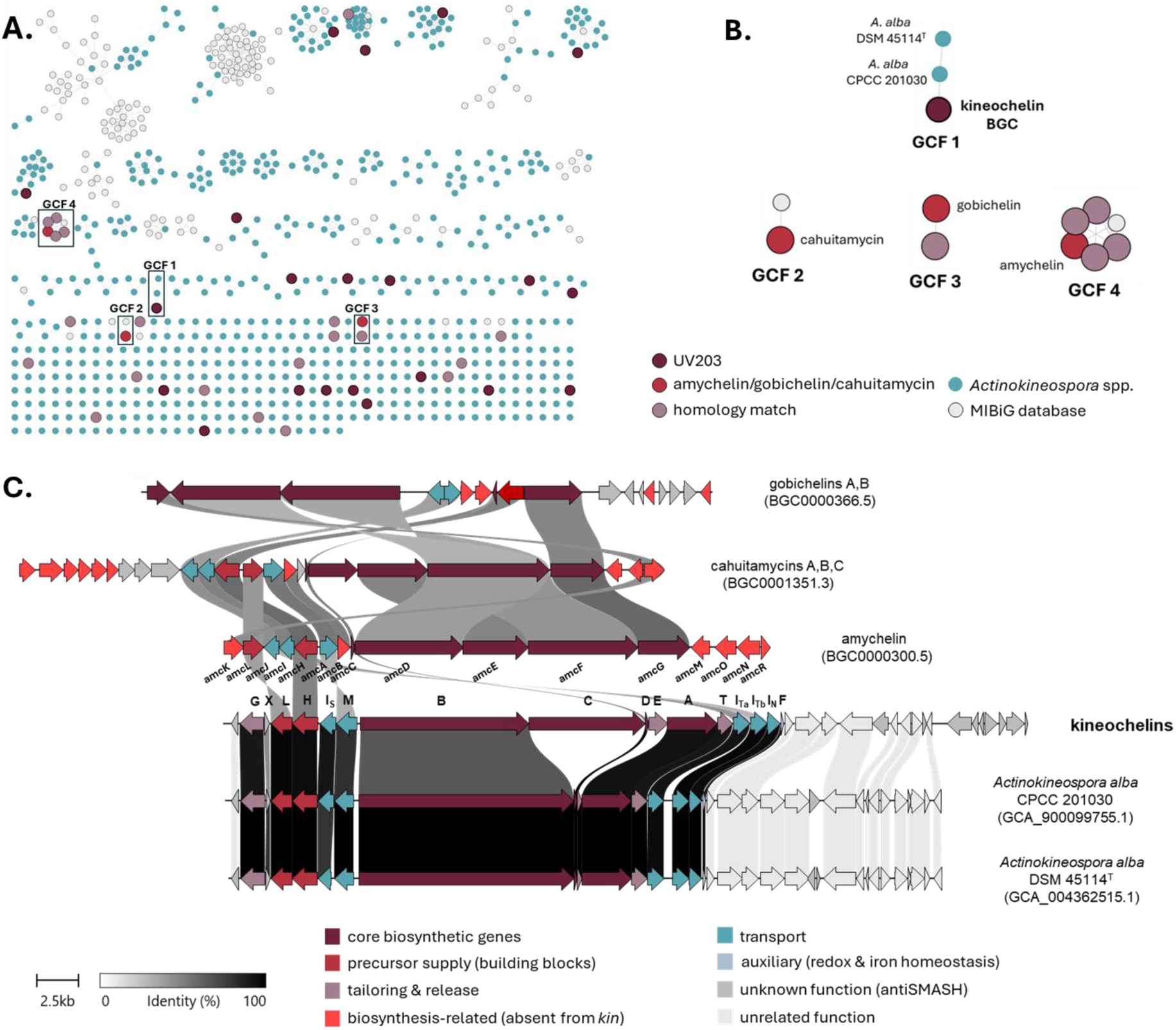
Kineochelin BGCs in *Actinokineospora alba* and strain UV203 represent a new gene cluster family. (A) BiG-SCAPE[60] sequence similarity network (SSN, cutoff 0.4) constructed from BGCs predicted in strain UV203 and all available *Actinokineospora* spp., together with the MIBiG database and three biosynthetically related siderophores. Groups of BGCs are color-coded as indicated in the legend. (B) Enlarged view of the gene cluster families (GCFs) containing the kineochelin, amychelin, cahuitamycin, and gobichelin BGCs. (C) Clinker[57] comparison within kineochelin cluster (member of GCF 1), with the amychelin, gobichelin, and cahuitamycin BGCs. Links between homologous genes are displayed according to percentage identity (see identity scale bar), with only links representing ≥40% amino acid identity shown. Gene arrows are drawn to scale, oriented by transcriptional direction, and colored by functional category.

### Iron-dependent expression links the kin BGC to kineochelin production

To confirm the role of the *kin* cluster in kineochelin biosynthesis, we initially attempted targeted gene inactivation using a single-gene knockout strategy. For both *kinA* and *kinB,* two knockout constructs containing 600 bp and 1000 bp intragenic gene regions, respectively, failed to produce exconjugants despite repeated conjugation attempts, whereas the control plasmid pSET152 integrated successfully, confirming the genetic tractability of *Actinokineospora* sp. UV203. Given these limitations, we shifted to a transcriptomics-based approach to assess the expression of the *kin* biosynthetic genes under iron-modulated conditions.

To select suitable conditions for comparative transcriptome analysis, we initially monitored growth, siderophore production, and antimicrobial activity of the wild-type strain during 10-day cultivation in three variants of SM17 medium: unmodified (SM17), supplemented with FeCl_3_ (SM17-Fe, iron-rich) or with the iron-chelator 2,2′-bipyridine (SM17-BP, iron-limited). In preliminary tests, both 100 µM and 200 µM FeCl_3_ suppressed siderophore production, with stronger inhibition at 200 µM; this concentration was therefore used as the iron-rich condition. Growth differed significantly, depending on the media used (Figure 6A). Growth in SM17 showed the highest yield, whereas both FeCl_3_ and bipyridine supplementation reduced the growth.

Bipyridine supplementation did not increase siderophore production relative to SM17, indicating that the kineochelin pathway is already derepressed under the baseline medium conditions (Figure 6B). Across the 10-day time course for SM17, siderophore titer rose sharply between days 4 and 6 (Figure 6B), coinciding with the first detectable antimicrobial activity (Figure 6C). Siderophore levels peaked at day 6, and antimicrobial activity at day 8, after which both remained stable.

Based on these dynamics, transcriptome sequencing was performed at two time points during growth: day 3, capturing the onset of SM production, and day 7, corresponding to peak activity. We performed comparative transcriptome analysis of strain UV203 grown in unamended and FeCl_3_-supplemented SM17, as iron-rich conditions showed almost complete repression of siderophore activity.

**Figure 6.**
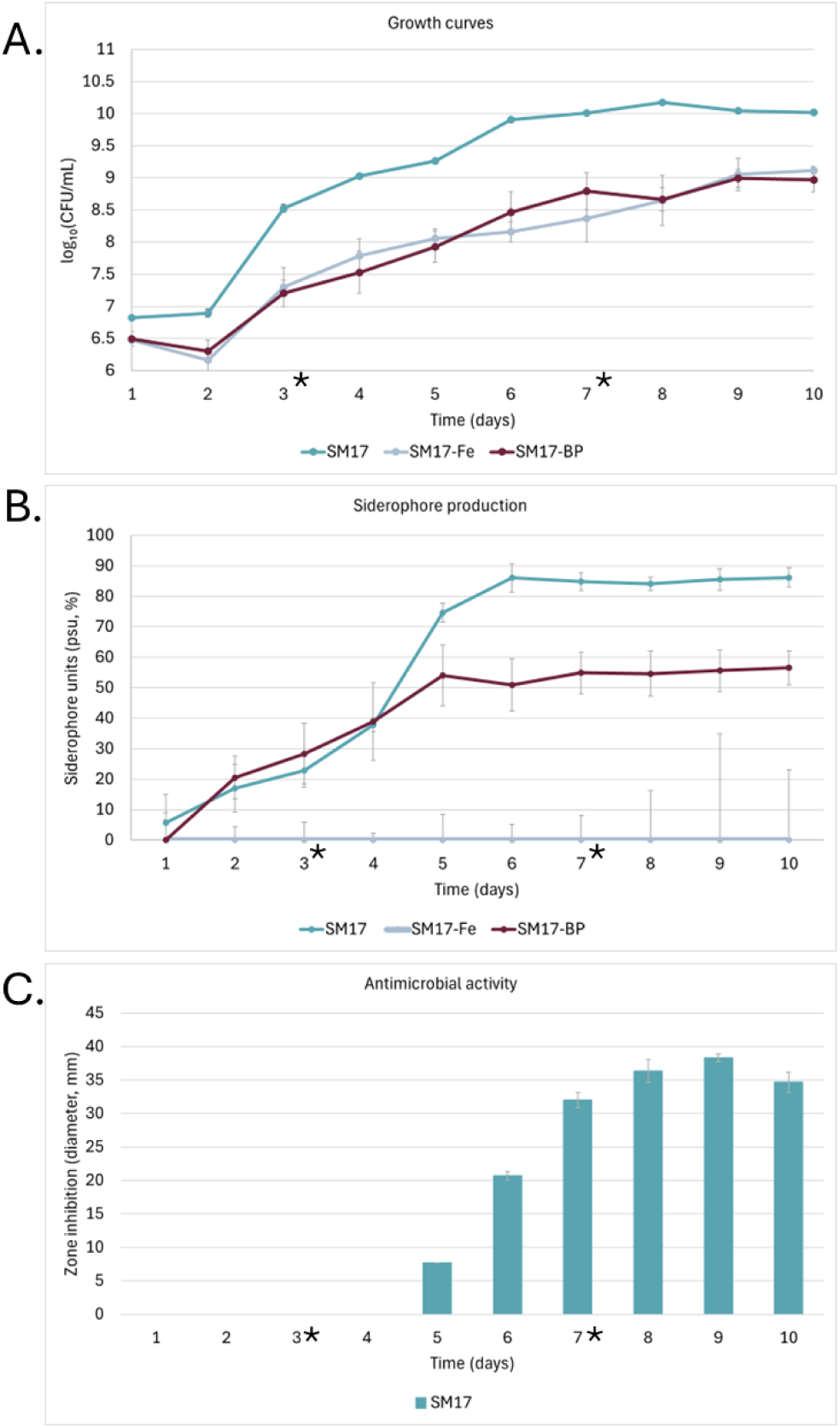
Growth, siderophore production, and antimicrobial activity of *Actinokineospora* sp. UV203 under three iron regimes. (A) Growth curves (log_10_ CFU/mL) in unmodified SM17 medium (SM17), supplemented with FeCl_3_ (SM17-Fe, iron-rich) or with the iron-chelator 2,2′-bipyridine (SM17-BP, iron-limited). Quantification by area under the growth curve (AUC) confirmed this pattern: SM17 = 91.95; SM17-Fe = 80.47; SM17-BP = 80.38. A one-way ANOVA detected significant differences among treatments (F₂,₆ = 18.85, p = 0.0026). Tukey’s HSD (HSD = 6.65) indicated significant reductions in growth in both SM17-Fe (Δ = 11.48) and SM17-BP (Δ = 11.57) relative to unamended SM17, while growth in SM17-Fe and SM17-BP did not differ (Δ = 0.09). (B) Overall siderophore production in culture supernatants measured as percent siderophore units (PSU) by the Chrome Azurol S (CAS)-shuttle assay. SM17-Fe showed no detectable siderophores (CAS values ≤ 0 plotted as 0). (C) Antimicrobial activity of crude extracts is shown as bars (left y-axis), representing inhibition zone diameters (mm) against *Micrococcus luteus* CCM 169^T^. Antimicrobial activity was detected only for the SM17 medium; no activity was observed for SM17-Fe or SM17-BP. Error bars indicate standard deviations (SD) from three biological replicates. Asterisks indicate sampling days for transcriptome analyses.

Principal component analysis (PCA) of the transcriptome data demonstrated clear separation of samples by both the conditions and time points, with tight clustering of biological replicates (Supplementary Fig. S37). Comparative transcriptomics showed that genes in the *kin* cluster were co-transcribed and significantly upregulated under siderophore-producing conditions (SM17 control) compared to iron-replete conditions (SM17 with 200 µM FeCl_3_) at both time points (Figure 7A,7B). Differential transcription of *kin* genes was consistent with selective production of kineochelins only under iron-limited conditions (Figure 7C, Supplementary Fig. S38). We identified an additional co-regulated gene outside the *kin* cluster, annotated as a lysine/ornithine N-oxygenase, and designated it *kinO* (Figure 7A and 7B).

**Figure 7.**
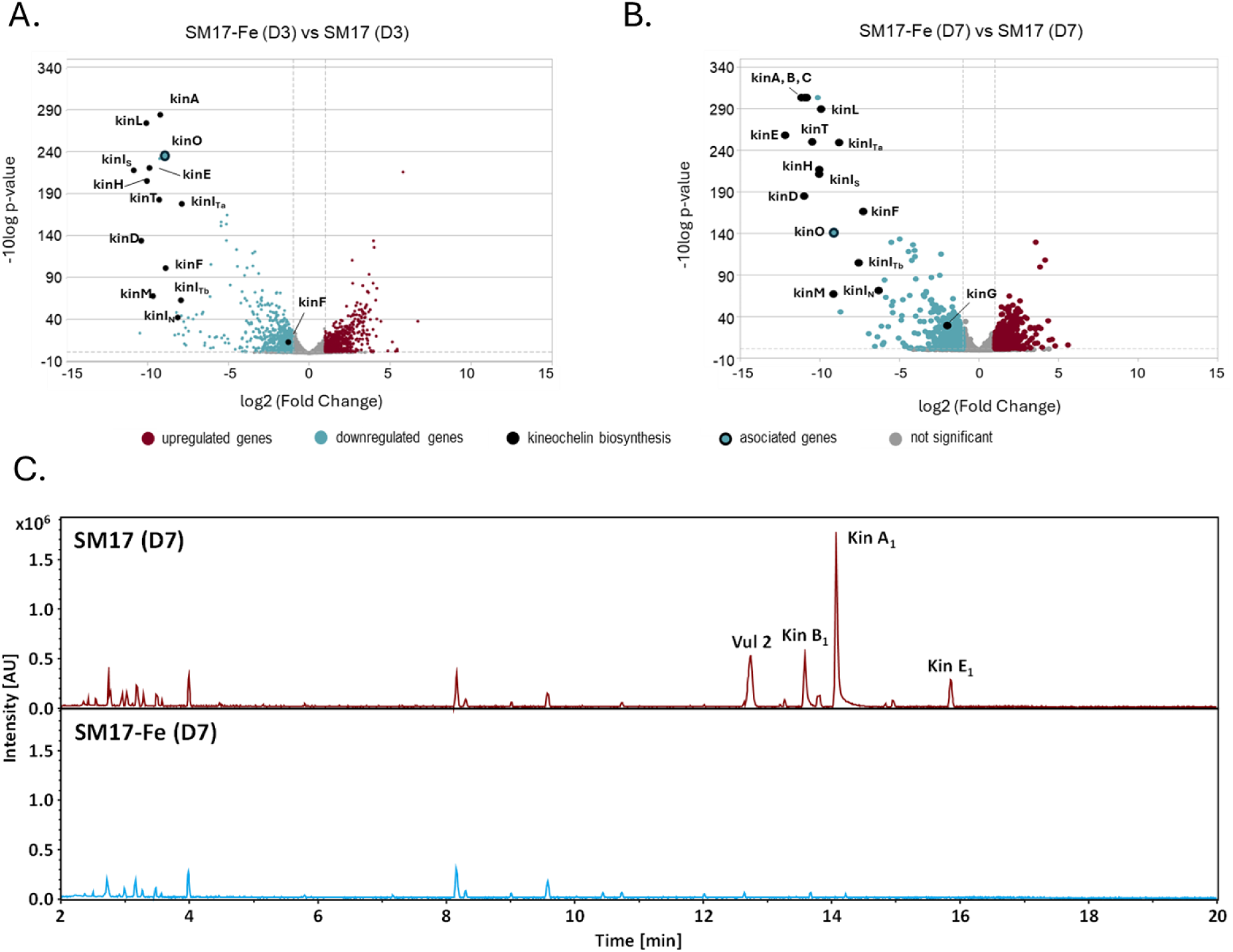
Comparative transcriptomics and metabolomics of *Actinokineospora* sp. UV203 under siderophore-producing and siderophore-depleted growth conditions. Volcano plots for the indicated pairwise comparisons. (A) SM17 supplemented with 200 µM FeCl_3_ versus SM17 (control) at day 3. (B) SM17 supplemented with 200 µM FeCl_3_ versus SM17 (control) at day 7. Each point represents a gene (x-axis: log_2_ fold change; y-axis: −log_10_(FDR)); Vertical dashed lines indicate the log_2_ fold-change threshold (|log_2_FC| ≥ 1), and the horizontal dashed line marks the FDR significance cutoff (FDR < 0.05; −log_10_FDR = 1.3). PCA and volcano plots were generated based on normalized counts from DESeq2. (C) Representative base peak chromatograms of the culture extracts of *Actinokineospora* sp. UV203 grown in SM17 supplemented with 200 µM FeCl_3_ versus SM17 (control) at day 7.

### Kineochelins exhibit metal-binding activity and iron-dependent growth-inhibitory effects

Preliminary bioactivity screening of crude extracts from *Actinokineospora* sp. UV203 against clinically relevant bacteria, yeasts and other isolates from Antarctic soils revealed growth-inhibitory effects, with the most pronounced inhibition observed against the strains from the same environment (Figure 8A, Supplementary Table S7). In addition, inhibitory activity against yeasts was detected, particularly *Nakaseomyces glabratus* and *Saccharomyces cerevisiae* isolates as well as against the bacterial indicator strain *Micrococcus luteus* CCM 169^T^ (Figure 8A, Supplementary Table S7). Follow-up testing with a pre-purified kineochelin-enriched extract, containing predominantly kineochelin A1, B1 and E1 with minor contributions from the co-produced siderophore vulnibactin 2 and trace-level co-occurring metabolites, including pseudomobactin A and asteroic acid (Supplementary Fig. S39), confirmed growth inhibition of both *M. luteus* CCM 169^T^ and yeasts in agar diffusion assays (Supplementary Table S8). However, due to limited extract availability and the greater clinical relevance of the yeast targets, subsequent minimum inhibitory concentration (MIC) and minimum fungicidal concentration (MFC) determinations were restricted to antifungal assays. Inhibition was observed only at relatively high concentrations, with MIC values of 0.5 mg/mL for *Nakaseomyces* strains and 2.5 mg/mL for *S. cerevisiae*, and corresponding MFC values of 2.5 mg/mL and 5 mg/mL, respectively (Supplementary Table S8). Due to these high MIC/MFC values, testing of individual purified kineochelin congeners was not feasible.

To assess the influence of iron availability on growth inhibition, sensitive yeasts and the bacterial indicator strain *M. luteus* CCM 169ᵀ were assayed on media supplemented either with excess iron or with the iron chelator 2,2′-bipyridine (Figure 8B). Iron supplementation markedly reduced growth inhibition, whereas iron depletion in the presence of extracts resulted in a complete growth arrest of sensitive yeasts. A comparable iron-dependent response was observed for *M*. *luteus* CCM 169^T^ (Figure 8B).

**Figure 8.**
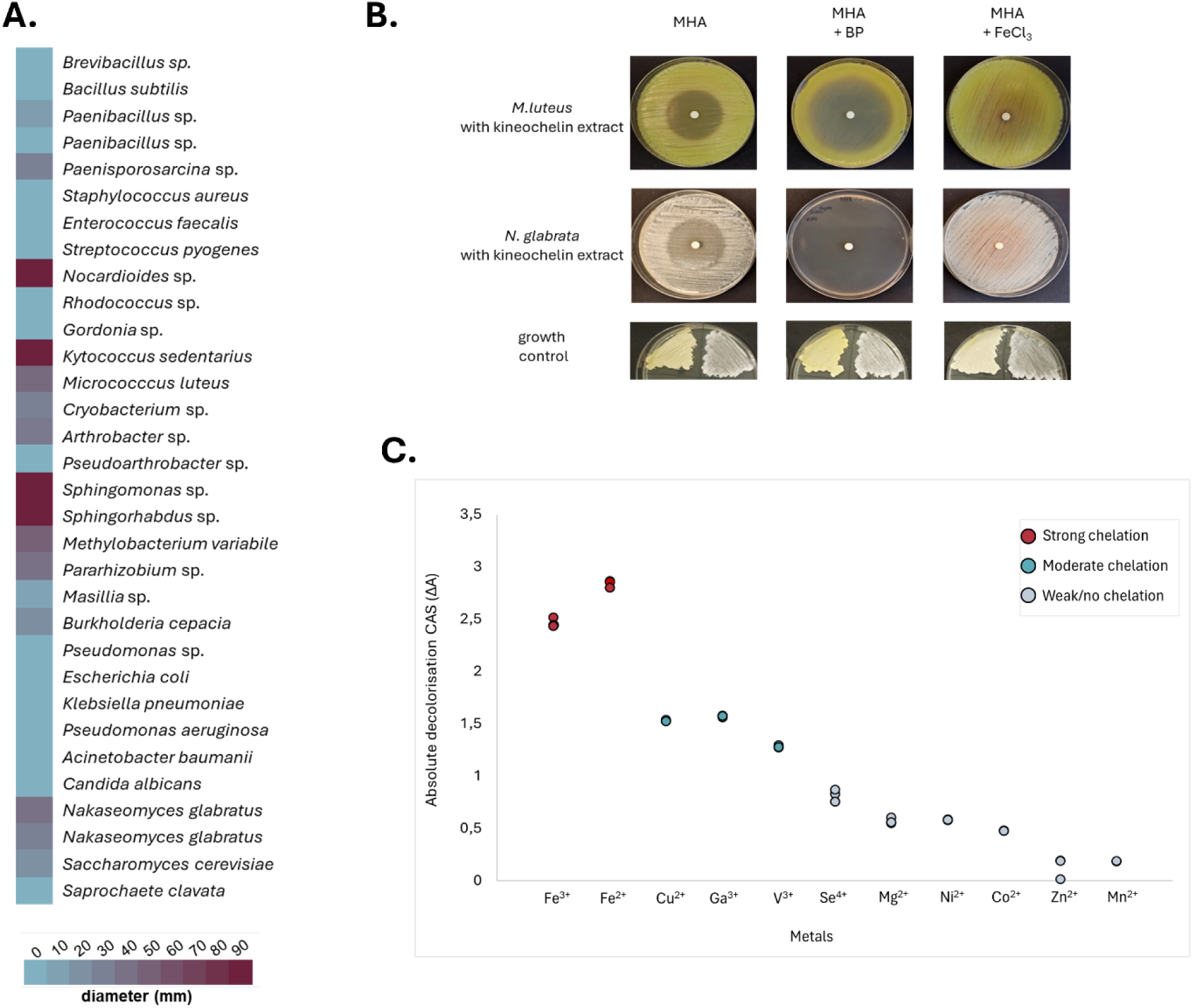
Iron-dependent growth-inhibitory effects and metal-binding properties of UV203-derived extracts. (A) Growth-inhibitory effects of crude extracts from *Actinokineospora* sp. UV203 against Antarctic environmental bacteria, the indicator strain *M. luteus* CCM 169ᵀ, and clinically relevant microbial strains. (B) Iron-dependent modulation of growth inhibition by a pre-purified, kineochelin-enriched extract (Supplementary Figure S39). Assays were performed on Mueller–Hinton agar (MHA), MHA supplemented with 100 µM 2,2′-bipyridine (MHA+BP), or MHA supplemented with 200 µM FeCl_3_ (MHA+FeCl_3_). For *N. glabratus*, media were additionally supplemented with 2% glucose. (C) Metal-binding activity of a pre-purified, kineochelin-enriched extract measured using the Chrome Azurol S (CAS) shuttle assay. Absolute CAS decolorization (ΔA) values for individual metals are shown. Points represent technical replicates (n = 3); colours indicate relative binding strength normalized to Fe³⁺.

To assess the metal-binding properties of kineochelins, we applied a modified Chrome Azurol S (CAS) assay to both whole culture supernatants of *Actinokineospora* sp. UV203 and a pre-purified kineochelin-enriched fraction (Figure 8C; Supplementary Figure S40). Both sample types showed a consistent hierarchy of metal-binding activity, with strongest responses toward ferric and ferrous iron, intermediate activity for Cu²⁺, Ga³⁺, and V³⁺, and weak or near-baseline responses for other tested metals.

We further assessed the antiproliferative activity of the pre-purified kineochelin mixture, tested as two independently handled aliquots (E1 and E2), against a panel of seven human cancer cell lines representing distinct tissue origins (A549, U-87, PaTu 8902, Jurkat, HCT116, A2058, and MDA-MB-231) using the resazurin assay. The calculated IC_50_ values are presented in Supplementary Table S9. Within the tested concentration range of 10 – 750 µg/mL, extract E1 exhibited inhibitory effect on the metabolic activity of A549 (545 ± 32 µg/mL) and A2058 cells (639 ± 32 µg/mL), whereas extract E2 demonstrated inhibitory activity against A549 (715 ± 16 µg/mL), U-87 (742 ± 27 µg/mL), and HCT116 (494 ± 48 µg/mL). The partially overlapping activity profiles observed for E1 and E2 are consistent with expected variability between independently handled aliquots of the same pre-purified extract.

## Discussion

In this study, we investigated the biosynthetic, chemical, and functional basis of siderophore production in *Actinokineospora* sp. UV203, a phylogenetically distinct actinomycete isolated from Antarctic soils. Although UV203 exhibited relatively narrow-spectrum antimicrobial activity in preliminary testing, predominantly against environmental strains, we prioritized it for untargeted secondary metabolome profiling based on the apparent novelty of its genomic biosynthetic potential and its distinct taxonomic placement. In the context of nutrient- and iron-limited Antarctic soil ecosystems, such activity is consistent with localized microbial competitive interactions rather than broad-spectrum antagonism[61]. This strategy proved effective, leading to the identification of kineochelins as the dominant SMs produced under iron-limited conditions.

Early metabolomic analyses of *Actinokineospora* sp. UV203 revealed a pair of low-abundance chlorinated metabolites containing two sulfur atoms. The combination of these structural features is rare and initially allowed tentative linkage of these compounds to a specific BGC encoding both a halogenase and cysteine-incorporating NRPS modules. However, despite targeted overexpression efforts, production levels of these metabolites remained insufficient for isolation and characterization, and this biosynthetic route was not pursued further.

In contrast, subsequent comparative metabolomic analyses revealed a distinct group of metabolites whose production was unexpectedly lost in an engineered regulatory mutant, and whose disappearance coincided with the loss of antimicrobial activity. The tentative structures derived from the LC-MS data did not match to any known compounds in public databases. The absence of database matches, together with the presence of shared mixed-ligand substructures, suggested that kineochelins represent a previously unrecognized family of siderophores, potentially related to other mixed-ligand metallophores such as such as amychelins[52] and gobichelins[53]. Since LC–MS analyses indicated that kineochelins are major metabolites of *Actinokineospora* sp. UV203 under siderophore-producing conditions, and their loss correlated with the disappearance of antimicrobial activity, we proceeded to isolate and structurally characterize representative members of this compound family.

Genome mining and comparative biosynthetic analyses identified the *kin* biosynthetic gene cluster as the most plausible candidate responsible for kineochelin production. Although the *kin* cluster shares biosynthetic logic with pathways directing the biosynthesis of amychelins, gobichelins, and cahuitamycins[51–53], it did not cluster with these pathways in BiG-SCAPE analyses, even under permissive similarity thresholds. This separation, together with clinker-based synteny comparisons, indicates that the *kin* cluster and related clusters in *A. alba* strains represent a distinct architectural lineage. Notably, the *kin* BGC in strain UV203 differs in gene content and organization from its closest homologs, most prominently by the presence of the *kinC* NRPS module, which is absent from the *A. alba* clusters. These differences are consistent with variation in peptide length and composition and likely contribute to the structural diversity of the resulting siderophores. More distant homology to the *gob*, *amc*, and *cah* BGCs, which direct the biosynthesis of gobichelins, amychelins, and cahuitamycins[51–53], respectively, indicates that kineochelin biosynthesis shares selected features with other mixed-ligand siderophore pathways, despite substantial divergence in gene content and organization.

Attempts to genetically validate the role of the *kin* cluster through targeted gene disruption were unsuccessful, likely reflecting low recombination efficiency associated with single-crossover insertion or locus-specific constraints on chromosomal integration in strain UV203[62]. As an alternative, transcriptomic analyses under iron-modulated conditions provided strong regulatory evidence linking the *kin* cluster to kineochelin production. Expression of *kin* genes was strongly upregulated under siderophore-producing conditions and repressed under iron-replete conditions, consistent with canonical siderophore regulation[63,64]. Co-expression of the *kin* genes, together with the identification of the co-regulated N-oxygenase gene *kinO*, supports a coordinated biosynthesis. The predicted function of KinO in supplying *N*-OH-L-Orn provides a mechanistic explanation for the incorporation of hydroxamate moieties into most kineochelins and further substantiates the assignment of the *kin* locus as the biosynthetic source of these compounds[65]. The incorporation of N⁵-hydroxyornithine via external gene is consistent with structurally related mixed-ligand siderophores, including amychelins and cahuitamycins[51,52]. In addition, the absence of a cis-acting thioesterase domain within the NRPS assembly line, together with the presence of a standalone ArCP–TE didomain protein (KinT) and the observed cyclic amide at the C-terminus of kineochelins, suggests that product release occurs via a trans-acting thioesterase. We therefore infer that off-loading proceeds through intramolecular cyclization mediated by KinT, rather than by canonical hydrolytic release[66].

Iron availability had a pronounced impact on growth, siderophore production, and bioactivity in UV203. Both iron excess and iron limitation impaired growth, likely through distinct physiological mechanisms. Growth inhibition under iron-rich conditions is consistent with iron-associated stress and suppression of iron uptake systems reported for other actinomycetes[67,68], whereas iron chelation by bipyridine slows the growth by restricting iron-dependent metabolic processes[69]. Notably, iron supplementation strongly reduced the growth-inhibitory activity of kineochelin-containing extracts, while iron depletion enhanced inhibition, resulting in complete growth arrest of sensitive bacterial indicator strain *M.luteus*. Lower siderophore levels observed at later time points under iron-chelated conditions likely reflect reduced biomass accumulation due to growth inhibition. Collectively, these observations are compatible with extracellular iron sequestration as the primary mechanism underlying kineochelin-mediated inhibition and support a role for kineochelins in iron competition under iron-limited conditions characteristic of Antarctic soils[29,70].

The observed selective sensitivity of *N. glabratus* and *S. cerevisiae*, but not *C. albicans*, to kineochelins likely reflects fundamental differences in their iron acquisition strategies. While all three species may activate reductive iron assimilation under low-iron conditions, *S.* c*erevisiae* relies predominantly on classical reductive iron uptake via surface ferric reduction followed by Fe²⁺ import through the Ftr1/Fet3 system [71,72], whereas *N. glabratus* has a more restricted iron acquisition repertoire, lacking several ferric reductases and depending mainly on ferrous iron and ferritin-derived sources[73,74]. Both species are therefore limited in their ability to exploit complex or protein-bound iron sources. In contrast, *C. albicans* possesses a highly versatile and redundant iron uptake network, including reductive assimilation, heme and hemoglobin utilization, transferrin and ferritin extraction, and siderophore uptake systems[75–79]. This flexibility enables *C. albicans* to switch efficiently between iron acquisition mechanisms when extracellular iron becomes limiting, thereby buffering intracellular iron homeostasis against siderophore-mediated depletion[80]. Collectively, the more constrained iron acquisition strategies of *N. glabratus* and *S. cerevisiae* likely render them more vulnerable to iron sequestration by kineochelins, explaining their selective sensitivity[71–82].

Consistent with this model, CAS assays demonstrated that kineochelins account for the majority of CAS-detectable metal chelation in UV203 cultures and function primarily as iron-chelating siderophores, with limited but reproducible secondary affinities for select non-iron metals. Such secondary metal-binding capabilities may become relevant under competitive or metal-limited environmental conditions but are unlikely to represent the primary biological function of kineochelins[64,83].

Finally, kineochelin-enriched extracts exhibited only weak antiproliferative activity against human cancer cell lines, with IC₅₀ values outside the range typically considered indicative of cytotoxic activity[84]. The higher IC₅₀ values observed for non-malignant fibroblasts compared to cancer cell lines suggest a degree of selectivity but overall indicate that kineochelins are unlikely to act as potent cytotoxins. Together, these findings reinforce the interpretation that kineochelins primarily function as ecological mediators of iron acquisition and competition rather than as broad-spectrum antimicrobial or cytotoxic agents.

## Conclusion and perspectives

Here, we report the discovery of a new group of mixed-ligand siderophores, the kineochelins, from strain UV203, which was isolated from Antarctic soil and represent a potentially novel *Actinokineospora* species. Bioactivity assays revealed selective inhibition of opportunistic yeasts (*N.s glabratus, S. cerevisiae),* whereas *C. albicans* showed resistance, an observation consistent with the divergent iron acquisition strategies of these yeasts. The activity of kineochelins was strongly dependent on iron availability, supporting a mechanism based on siderophore-mediated iron sequestration. Although antimicrobial activity was observed only at high concentrations, kineochelins provide a chemically tractable scaffold for future structure–activity studies aimed at understanding how variation in siderophore architecture influences iron acquisition, competitive interactions, and microbial community dynamics in iron-limited extreme environments.

Beyond their biomedical relevance, the pronounced iron selectivity and mixed-ligand architecture of kineochelins highlight their broader biotechnological potential. Metallophores with defined metal preferences have attracted interest as tools for manipulating metal availability in engineered microbial systems, metal mobilization and recovery, and the study of metal-dependent microbial interactions. While such applications were not addressed here, the chemical tractability and biosynthetic accessibility of kineochelins provide a foundation for future investigations. More broadly, the discovery of kineochelins in an Antarctic bacterium reinforces the value of polar microbiomes as reservoirs of structurally novel NPs and the need to integrate ecological and genomic perspectives in natural product discovery.

## Methods

### Biological resources: Isolation, culture conditions, site description

James Ross Island is part of the James Ross Island group, which comprises four larger islands and several smaller landmasses located near the northernmost tip of the northeastern Antarctic Peninsula (Trinity Peninsula)[85]. Active layers from James Ross Island are regularly sampled as part of microbiological biodiversity surveys conducted under the Czech Antarctic Research Programme, which has maintained a long-term biomonitoring initiative in the area since 2007. *Actinokineospora* sp. UV203 was isolated from an active layer sample at a depth of 10 cm below the surface at Abernathy Flats (63°50’47.5“S, 57°53’23.9”W) collected in the austral summer of 2022 on James Ross Island.

To selectively cultivate sporulating bacteria, one gram of the sediment sample was pre-treated by air-drying for 48 hours, then suspended in 9 mL of sterile saline (0.9%) solution by vortexing. The suspension was incubated in a water bath at 50 °C for 5 minutes[86], followed by serial ten-fold dilutions up to 10⁻⁵. Aliquots of 100 µL of each dilution were spread onto a range of actinomycete-selective media, including starch-casein, starch-nitrate, International Streptomyces Project (ISP) 2, humic acid–vitamin, and Actinomyces agar (Merck, DE) plates[87–90]. Each medium was supplemented with filter-sterilized cycloheximide (50 mg/L) and nalidixic acid (20 mg/L) to suppress the growth of fungi and fast-growing Gram-negative bacteria[86]. The inoculated plates were incubated at 14 °C for up to three months, during which colony development was regularly monitored, and morphologically distinct isolates were selected for classification and preserved at –80 °C in glycerol stocks.

### 16S rRNA gene-based identification of isolates

DNA was isolated using a rapid Chelex-based extraction protocol (Chelex 100 Resin, Bio-Rad), and the nearly full-length 16S rRNA gene was amplified using the 616V and 1492R primer pair[91]. Initial identification of *Actinokineospora* sp. UV203 was based on the 16S rRNA gene sequence similarity using EZBioCloud[92] and the preliminary taxonomic position was confirmed through phylogenetic analysis using MEGA X[93]. Homologous sequences were aligned using MUSCLE[94], and phylogenetic tree was reconstructed using the maximum likelihood method. Tree robustness was assessed by bootstrap analysis with 1000 replicates.

### Whole genome sequencing and genome assembly

DNA extraction and whole genome sequencing were performed by the Joint Microbiome Facility of the Medical University of Vienna and the University of Vienna under the project number JMF-2209-13. High molecular weight DNA was extracted using the Monarch gDNA purification kit (NEB), with the specific protocol for Gram positive bacteria. Pellets were initially treated with an enzymatic cocktail (innuPREP Bacteria Lysis Booster, Innuscreen GmbH) to facilitate chemical cell wall lysis. Samples were diluted and equimolarly barcoded using the SQK-RBK114.96 (Oxford Nanopore Technologies) with the following protocol modifications: we increased the sample input to 230ng/sample, +0.5ul of rapid adaptor to the barcoded library. About 80fmol of a >28Kb library was loaded on a R10.4.1 flowcell (FLO-PRO114, Oxford Nanopore Technologies) and sequenced for 20h on a Promethion P2 solo (Oxford Nanopore Technologies, ONT, UK) using Minknow (v. 23.11.7, ONT, UK). Flowcell light shields were used. Reads were basecalled using Dorado server basecaller v7.8.2 (ONT, UK) using super accuracy mode. The nanopore reads were assembled using flye v. 2.9.3[95], with “–nano-hq” and polished once with medaka (v. 1.11.3, github.com/nanoporetech/medaka). The model used was “r1041_e82_400bps_sup_v4.2.0”. Contigs <1000bp were removed and assemblies were quality checked with QUAST v5.2.0[96], and CheckM v1.2.2[97]. Details on assembled genome are provided in Supplementary Table S10.

### Genomic analyses

Initial genome-based taxonomic classification was performed using the TYGS server and Genome-to-Genome distance calculator v3.0 (GGDC) using formula 2 (TYGS formula d_4_) digital DNA–DNA hybridization (dDDH) to assess relatedness between UV203 and the type strains of all validly described *Actinokineospora* species[32,98]. The average nucleotide identity (ANI) values were calculated using FastANI to further evaluate genome-level similarity[33]. The dDDH was calculated using formula d4 of GGDC. A maximum likelihood phylogenomic tree was constructed based on 92 single-copy marker proteins identified by the Up-to-date bacterial core gene set (UBCG v2) pipeline[31]. Genome annotation was performed using Bacta v1.11.3[99] and the resulting GenBank (.gbk) files were used to predict BGCs via the standalone version of antiSMASH v7.1[100]. Functional annotations for genes within each cluster, as well as flanking regions where possible, were refined using BLAST and InterPro, in addition to initial predictions[101,102]. For sequence similarity network (SSN) analysis, multiple *Actinokineospora* genomes and the closest homologs from the MIBiG and antiSMASH databases were retrieved from NCBI. These were included in BiG-SCAPE v2.0.0 analysis using a similarity cut-off of 0.4[60,103]. The resulting networks were visualized in Cytoscape v3.10.3[104]. Selected BGCs were compared using Clinker, applying a minimum sequence alignment identity threshold of 40%[57].

### Cultivation of Actinokineospora sp. UV203 for production of specialised metabolites and small-scale extraction

A seed culture of *Actinokineospora* sp. UV203 was prepared by inoculating 25 ml of tryptone soy broth (TSB; Oxoid™, USA) with 100ul of heat-activated spores from a glycerol stock (40 °C for 10 min). The culture was incubated at 21 °C with shaking at 200 rpm for 2 days. Subsequently, 5% (v/v) of the seed culture was used to inoculate 50 mL of liquid bioproduction media in 250 mL non-baffled Erlenmeyer flasks. The tested media included: TSB, ISP2[88], SM17[105], 5254 (g/L: glucose 15.0, soy meal 15.0, corn steep liquor 5.0, CaCO_3_ 2.0, NaCl 5.0; pH 7.0), 5304 (g/L: glucose 1.0, soluble starch 24.0, tryptone 5.0, meat extract 3.0 and CaCO_3_ 4.0, NaCl 5.0; pH 7.0) and 5288 (g/L: glycerol 15.0, soy meal 10.0, NaCl 5.0, CaCO_3_ 1.0, CoCl_2_x7H_2_O 0.001; pH 6.8). Two parallel cultures were prepared for each medium and incubated at 21 °C and 200 rpm for 5 and 10 days, respectively. Following incubation, the fermented broths were harvested, frozen at -80°C and freeze-dried for 72 hours until completely dry. The dried material was extracted with methanol (1:1, v/v) by shaking at room temperature for 2 hours at 150 rpm. The extracts were centrifuged at 10,000 rpm for 10 minutes to remove undissolved particles. The resulting organic supernatant was collected and evaporated to dryness using a rotary evaporator at 40 °C and 280 mbar, yielding concentrated crude extracts. These were dissolved in 2 mL of methanol, and 250 µL of each extract was filtered through 0.45 µm Whatman® Mini-UniPrep® G2 syringeless filters (Cytiva, USA) prior to mass spectrometry analysis. Remaining extracts were stored at -20 °C until used for bioactivity tests.

### Mass Spectrometry

LC-MS analyses were performed on a Vanquish Horizon UHPLC system (Thermo Fisher Scientific) equipped with an Acquity Premier HSS T3 column, 2.1 x 150 mm, 1.8 µm (Waters) coupled to the ESI source of a timsTOF fleX mass spectrometer (Bruker Daltonics) as described previously[106]. Compass DataAnalysis 5.3 (Bruker Daltonics), GNPS, The Natural Products Atlas, and CAS SciFinder (American Chemical Society) were used for data analysis[37],[107].

^1^H and ^13^C (DEPTq) 1D as well as COSY, HSQC, and HMBC 2D NMR spectra of the isolated compounds **1-4** in CD_3_OD at 298 K were recorded on an Avance NEO 600 NMR spectrometer (Bruker BioSpin) equipped with a N_2_ cryo probe Prodigy BBFO with z-gradient (600.18 MHz for ^1^H, 150.92 MHz for ^13^C). Chemical shifts were calibrated using the ^1^H residual solvent signal at δ = 3.31 and the ^13^C solvent signal at δ = 49.15.

### Isolation and structural elucidation of kineochelins

The upscaled culture broth of *Actinokineospora* sp. UV203, harvested on day 10 of cultivation at 21 °C and 200 rpm, was used for the isolation of kineochelins. The total volume of 300 mL was obtained by pooling six parallel 50 mL subcultures grown under identical conditions. The combined culture broth was centrifuged at 10,000 rpm for 10 minutes, and the resulting supernatant was mixed with 5% (v/v) of methanol-activated and sterilized Diaion® HP-20 resin (Supelco, Sigma-Aldrich, USA)[108]. The mixture was shaken at 21 °C and 200 rpm for 2 hours to facilitate adsorption of secondary metabolites. After incubation, the aqueous phase was discarded, and the resin was extracted with *n*-butanol under the same shaking conditions (200 rpm, 2 hours). The butanol extract was then transferred to a clean glass flask and evaporated to dryness using a rotary evaporator at 40 °C. The resulting crude extract was dissolved in 3 mL of methanol and fractionated using flash chromatography (FC), which was performed on a PuriFlash 4250 from Interchim equipped with both a photodiode array detector (PDA) and an evaporative light scattering detector (ELSD). The run was performed in reversed phase mode using a PuriFlash 15 C18 HQ 35G column (35.0 g, 22 bar). The mobile phase consisted of water + 0.1 % formic acid (FA) (A) and acetonitrile:water (9:1) + 0.1 % FA (B). The following gradient was applied: 5-20 % B in 10 min, 20-75 % B in 60 min, 75-98 % B in 5 min and 98 % B for 10 min. The flow rate was set to 15 ml/min. The collected tubes were analyzed by UHPLC using a Waters Acquity UPLC H-Class system (Waters, USA), equipped with a sample manager, a quaternary solvent manager and a column manager.

For detection the system was coupled with an ELSD and a PDA detector. The UPLC H-class system was controlled using Empower 3 software. For analysis, an Acquity TSS H3 (2.1 x 100 mm; 1.8 μm) (Waters, USA) was used. The chromatographic conditions were set as followed: flow rate 0.3 mL/min, water + 0.1 % FA (A) and acetonitrile:water (9:1) + 0.1 % FA (B), gradient: 5-20 % B in 10 min, 20-75 % B in 15 min and 75 % B isocratic for 3 min. The tubes were pooled according to UHPLC-PDA-ELSD traces into 21 fractions. Following LC-MS analysis, fractions 11 (5.83 mg) and 17 (1.96 mg), containing mainly kineochelin A_1_ and E_1_, respectively, were subjected to NMR analysis.

Marfeýs derivatization was employed to determine the absolute configurations of amino acids in kineochelin E_1_ (fraction 16) and A_1_ (fraction 10). Samples of kineochelin E_1_ (410 μg) and A_1_ (750 μg) were dissolved in 2 M HCl, subjected to acid hydrolysis, dried and redissolved in 1 M NaHCO, followed by derivatization with 1% Marfey’s reagent (1-fluoro-2,4-dinitrophenyl-5-L-alanineamide (L-FDAA)) in acetone. Subsequently, the mixtures were heated at 40 °C for 1h. Thereafter, neutralisation with 1M HCl was carried out, followed by dilution with acetonitrile for LC-MS analysis. For comparison, standard amino acids, L/D-Orn, L/D-Thr, L/D-allo-Thr, and L/D-Ser, were similarly derivatized with L-FDAA.

### Growth and kineochelin production dynamics of Actinokineospora sp. UV203

To assess the growth kinetics and dynamics of kineochelin production under varying iron conditions, we used a 72-hour seed culture and subsequent culture prepared as described above. Cultivation was carried out in 50 mL volumes of the following media: SM17 (control), SM17 supplemented with 100 µM and 200 µM FeCl_3_ (iron-rich condition), and SM17 supplemented with 100 µM 2,2′-bipyridine (iron-limited condition). Both supplements were filter-sterilized and added to the media prior to inoculation. Each condition was tested in four biological replicates. Cultures were sampled at 12-hour intervals for two purposes: (i) colony-forming unit (CFU) determination using 100 µL aliquots, and (ii) analysis of kineochelin production and total siderophore activity using 2 mL samples. CFU determination was performed using the plate count method, with serial dilutions in 0.9% NaCl, and plating onto SFM agar[109]. CFU values were log_10_-transformed prior to visualization. Growth curves were plotted from the mean of three biological replicates, with error bars representing standard deviation (SD). The area under each growth curve (AUC) was calculated using the linear trapezoidal rule[110]. Differences in AUC among conditions were tested by one-way ANOVA, followed by Tukey’s HSD post-hoc test[111].

To quantify iron-chelating activity during growth, we used a universal Chrome Azurol S (CAS) assay in 96-well microplate format[112,113]. Briefly, 100 µL of culture supernatant or extract solution was mixed with an equal volume of CAS reagent containing Fe³⁺, incubated in the dark for 20 min, and absorbance was measured at 630 nm[113]. Siderophore production was quantified as percent siderophore units (PSU), calculated using the formula published by Payne (1993)[114]. All measurements were performed in three independent biological replicates. CAS absorbance values were background-corrected using CAS reagent mixed with uninoculated medium containing the corresponding supplements. Negative values, which can arise from iron carryover in iron-supplemented cultures, were interpreted as baseline (no detectable siderophore activity) and truncated to zero for visualization.

To assess metal-binding specificity beyond ferric iron, a modified CAS assay was performed using CAS complexes prepared with alternative metal ions. CAS reagents were freshly prepared using chloride salts of Fe²⁺, Mg²⁺, Mn²⁺, V³⁺, Zn²⁺, Se⁴⁺, Co²⁺, Cu²⁺, and Ni²⁺, and gallium bromide (Ga³⁺), following published protocols[115,116]. To determine whether kineochelins alone are sufficient to account for the CAS activity observed in culture supernatants, a pre-purified kineochelin-enriched fraction was analyzed. All measurements were performed using technical replicates (n = 3). Because culture media and extract solutions exhibited intrinsic absorbance at 630 nm, all CAS measurements were corrected for sample-specific background signals. Metal-binding activity was quantified as absolute CAS decolorization (ΔA), calculated as the difference in absorbance between the corresponding metal–CAS control and sample after subtraction of relevant background signals.

### Bioactivity assays

Antimicrobial bioactivity was assessed using the disc diffusion method with 6-mm Whatman paper discs. Discs were impregnated with either 50 µL of crude extract for bacterial assays or 100 µL for yeast assays. For tests involving pre-purified kineochelins, extracts were first dried, weighed, and dissolved to a known concentration (mg/mL). Inhibition zones were recorded after 18–48 hours of incubation at appropriate temperatures. Mueller–Hinton agar (MHA) was used for clinically relevant bacterial strains, while yeasts were tested on MHA supplemented with 2% glucose and 0.5 µg/mL methylene blue dye, as recommended by The Clinical & Laboratory Standards Institute (CLSI) guidelines^79,80^. For environmental isolates, media were selected based on organism-specific growth requirements; MHA was used where appropriate (Supplementary Table S1). The sensitivity profiles of the clinical yeast isolates used in this study were determined by the Faculty Hospital in Brno, as listed in Supplementary Table S11.

To determine the minimum inhibitory concentrations (MICs) against sensitive yeast strains, we followed the 96-well microtiter plate protocol according to CLSI guidelines[117,118]. RPMI 1640 medium, supplemented with glucose and buffered with MOPS to pH 7.0, was used for testing *N. glabratus* strains. For *S. cerevisiae*, which failed to grow in both RPMI and MHA with glucose, YPD medium (DSMZ medium 393) was used instead. Pre-purified kineochelins were tested starting from a concentration of 20 mg/mL using two-fold serial dilutions. Amphotericin B served as a positive control, starting at 125 µg/mL.

Human cancer cell lines for the initial screening of potential antiproliferative activity of kineochelins, the were purchased from ATCC (American Type Culture Collection; Manassas, VA, USA) or ECACC (European Collection of Authenticated Cell Cultures, Salisbury, United Kingdom). The Jurkat (human leukaemic T cell lymphoma) and HCT116 (human colorectal carcinoma) were cultured in RPMI 1640 medium (Biosera, Kansas City, MO, United States), while U-87 MG (human glioblastoma), A2058 (human metastatic melanoma) and MDA-MB-231 (human mammary gland adenocarcinoma) cells were maintained in growth medium consisting of high-glucose Dulbecco’s Modified Eagle Medium (DMEM) supplemented with sodium pyruvate (Biosera, Kansas City, MO, United States). Specific medium requirements were necessary for PaTu 8902 (human pancreatic adenocarcinoma) cells, which were maintained in high-glucose DMEM supplemented with sodium pyruvate (Biosera, Kansas City, MO, United States) and 25 mM HEPES (Sigma, Steinheim, Germany). The healthy, non-tumorigenic cell line CCD-18Co (non-malignant intestinal fibroblasts) was cultured in DMEM (Biosera, Kansas City, MO, USA). All culture media were supplemented with 10% FBS, or 15% in the case of the A2058 cell line (fetal bovine serum; Gibco, Thermo Scientific, Rockford, IL, USA), and an antibiotic–antimycotic solution (Merck, Darmstadt, Germany). Throughout the experiment, cell lines were cultured at 37 °C in a humidified atmosphere containing 5% CO_2_. The effects of a pre-purified kineochelin-enriched extract, tested as two independently handled aliquots (designated E1 and E2), were determined using a resazurin reduction-based assay. A549, U-87 MG, PaTu 8902, Jurkat, HCT116, A2058, MDA-MB-231 and CCD-18Co (5x10^3^/well) cell lines were seeded and cultured in 96-well culture plates. After 24 hours, cells were treated with E1, E2 pre-purified kineochelin extracts (concentration range 10 – 750 μg/ml) and incubated for 72 hours. After 72 h incubation, 10 µl of resazurin dye was added to the each well, followed by incubation for minimal 1.5 h. The fluorescent output was measured using the automated Cytation™ 3 Cell Imaging Multi-Mode Reader (Biotek, Winooski, VT, USA).

### Knock-out mutagenesis and initial overexpression approaches

All standard molecular biology techniques, including DNA manipulations, cloning procedures, and plasmid transformation into *Escherichia coli* strains, were performed as previously described [119]. PCR amplifications were carried out using Q5® High-Fidelity DNA Polymerase (New England Biolabs, Ipswich, MA, USA) with oligonucleotides listed in Supplementary Table S12. A complete list of plasmids and bacterial strains used or constructed in this study is provided in Supplementary Table S13. Luria-Bertani (LB) medium was used for routine cultivation of *E. coli*, supplemented with chloramphenicol (30 µg/mL), kanamycin (30 µg/mL), and when necessary, apramycin (Am, 100 µg/mL). *E.coli* DH5α was employed for general cloning, while *E. coli* ET12567 (pUZ8002) was used to mediate intergenic conjugative plasmid transfer into *Actinokineospora* sp. UV203. Initial attempts to activate BGC 2.18 involved overexpression of LuxR- and HxlR-like (Reg2) regulators encoded within the cluster, using pSET152-based constructs driven by either ermE*p[120] or a native *rrn* operon promoter. Following the unsuccessful activation of BGC 2.18, we focused subsequent genetic manipulation efforts on the *kin* BGC. To assess the roles of candidate biosynthetic genes in kineochelin production, two *kin* cluster genes, kin_2475 (*kinB*) and kin_2479 (*kinA*), were selected for inactivation. Each gene was PCR-amplified using primers incorporating *EcoRI* and *HindIII* restriction sites, and cloned into the 3.1 kb *EcoRI/HindIII* fragment of the pSOK201 vector containing ColE1, *oriT* and Am^R^ [121]. The resulting plasmids, pKO_2475 and pKO_2479, were confirmed by sequencing and subsequently conjugated into the wild-type *Actinokineospora* sp. UV203 strain. Apramycin (50 µg/ml) was used for selection of recombinant *Actinokineospora* strains and nalidixic acid (30 µg/mL) as counterselection against *E. coli*. No exconjugants were recovered for either construct under the conditions tested. As a conjugation control, an empty pSET152 vector[122] was introduced into the wild-type strain using the same conjugation procedure.

### Transcriptomics

RNA extraction and sequencing was performed by the Joint Microbiome Facility of the Medical University of Vienna and the University of Vienna (project JMF-2412-11). RNA was extracted using the Monarch Total RNA Miniprep Kit (New England Biolabs) according to the manufacturer’s instructions but including two rounds of DNA digestion with ezDNAse (Thermo Fischer Scientific). Sequencing libraries were prepared from rRNA depleted (Ribo-Zero Plus rRNA Depletion Kit, Illumina) RNA samples using the NEBNext® Ultra™ II Directional RNA Library Prep Kit for Illumina, New England Biolabs according to the manufacturer’s instructions and sequenced in 2x 100 bp paired-end mode ((Illumina NextSeq 6000 SP 1/2 flowcell)), yielding 76 million raw reads per sample. Individual read libraries were quality checked using fastQC v0.12.1[123] and quality statistics were merged using multiQC v1.21[124]. Original bam files were converted to fastq format using samtools[125] v1.12’s bam2fq function. Fastq files were trimmed and filtered using BBDuk, in BBMap v39.10. PhiX and adapter sequences were removed and the chastityfilter option was set to “true”. Reads were also trimmed at a q-score of 28, with an average q-score of 15, keeping a minimum length of 30 nucleotides and the following other options: “ktrim=r k=23 mink=11 hammingdistance=1 qtrim=r”. Reads were mapped to the reference genome UV203 using BBMap with a minimum percent identity of 98% and ambiguous reads were mapped to all locations. The resulting sam file was converted to a bam file using samtools. FeatureCounts, part of SubRead v2.1.1, was used to generate a counts table for reversely-stranded, paired-end reads with a PGAP-generated .gff file as a reference. DESeq2 release v1.48[126] was used to assess differentially-expressed genes and calculated FPKMs for each. Normalized counts from DESeq2 were used to generate principal component analysis (PCA) plots and volcano plots (log_2_ fold change, log_2_FC, vs. −log_10_ false discovery rate, FDR), applying significance thresholds of |log_2_FC| ≥ 1 and FDR < 0.05. antiSMASH BGC predictions were overlaid on PGAP annotations based on genome coordinates using in-house parsing script (https://github.com/kralovaresearch/Kineochelin).

## Declarations

### Data availability

The genome sequence and transcriptomic data of strain *Actinokineospora* sp. UV203 is available on NCBI (BioProject accession number PRJNA1331526). The nearly full-length 16S rRNA gene (1,395 bp) of strain *Actinokineospora* sp. UV203 is available on NCBI (accession number PX090945). The NMR data of kineochelin E_1_ and A_1_ are deposited in the Natural Products Magnetic Resonance Database (NP-MRD) under accession numbers NP0352113 and NP0352114, respectively.

### Authors’ contributions

S.K. and A.L. conceived the study. S.K., P.S., J.G., V.M., and M.B. - performed experiments and analysed data. A.L., M.Z., S.B.Z., U.G., O.N.S., M.B. and T.R, - contributed essential experimental infrastructure and analytical instrumentation, and expert advice. J.S.C.D.S. and J.O. – performed initial processing of genome and transcriptome data. S.K. - performed bioinformatic analyses. S.K., M.Z., U.G., A.L., and S.B.Z. - interpreted the data. S.K. and M.Z. wrote the article. All authors revised and approved the manuscript.

### Ethics approval and consent to participate

Not applicable.

### Consent for publication

Not applicable

## Acknowledgements

The Life Science Compute Cluster LiSC at the University of Vienna provided the high-performance computing infrastructure for this study. We thank Julia Ramesmayer and Sara Malinowski (Joint Microbiome Facility of the Medical University of Vienna and the University of Vienna) for assistance during high molecular weight extraction and RNA extraction. The authors thank Anna Fabisikova and Michael Klemm-Abraham from the Mass Spectrometry Centre and the team of the NMR Centre (both core facilities of the Faculty of Chemistry, University of Vienna, and members of the Vienna Life Science Instruments) for assistance with data acquisition. We are thankful to Dr. Jaime Felipe Guerrero Garzón for helpful discussions on the use of a *rrn* operon promoter strategy. For open access purposes, the authors have applied for a CC BY public copyright license to any author-accepted manuscript version arising from this submission. Dr. Martin Kello (Department of Pharmacology, Faculty of Medicine, Pavol Jozef Šafárik University, Košice, Slovakia) and Dr. Michal Goga (Department of Plant Biology, Faculty of Science, and Center for Interdisciplinary Biosciences, Technology and Innovation Park, Pavol Jozef Šafárik University in Košice, Košice, Slovakia) funded by VEGA 1/0498/23 are acknowledged for their assistance with the antiproliferative assays.

## References

1. Murray CJ, Ikuta KS, Sharara F, Swetschinski L, Aguilar GR, Gray A, et al. Global burden of bacterial antimicrobial resistance in 2019: a systematic analysis. The Lancet. 2022;399:629–55. 10.1016/S0140-6736(21)02724-0

2. Cardona ST, Rahman ASMZ, Novomisky Nechcoff J. Innovative perspectives on the discovery of small molecule antibiotics. npj Antimicrob Resist. 2025;3:19. 10.1038/s44259-025-00089-0

3. World Health Organization. Antibacterial Agents in Clinical and Preclinical Development: An Overview and Analysis. World Health Organization; 2023.

4. Butler MS, Henderson IR, Capon RJ, Blaskovich MAT. Antibiotics in the clinical pipeline as of December 2022. J Antibiot. Nature Publishing Group; 2023;76:431–73. 10.1038/s41429-023-00629-8

5. Kirienko NV, Rahme L, Cho Y-H. Editorial: Beyond Antimicrobials: Non-traditional Approaches to Combating Multidrug-Resistant Bacteria. Front Cell Infect Microbiol. Frontiers; 2019;9. 10.3389/fcimb.2019.00343

6. El-Sayed SE, Messiha AA, Zafer M. Effective alternative strategies to combat challenges associated with MDR bacterial infections: Drug repurposing, role of artificial intelligence, and novel therapeutic options. Journal of Infection and Public Health. 2026;19:103058. 10.1016/j.jiph.2025.103058

7. Masschelein J, Jenner M, Challis GL. Antibiotics from Gram-negative bacteria: a comprehensive overview and selected biosynthetic highlights. Nat Prod Rep. The Royal Society of Chemistry; 2017;34:712–83. 10.1039/C7NP00010C

8. Hegemann JD, Birkelbach J, Walesch S, Müller R. Current developments in antibiotic discovery. EMBO Rep. 2022;24:e56184. 10.15252/embr.202256184

9. Barry SM. Rethinking natural product discovery to unblock the antibiotic pipeline. Future Microbiology. 2025;20:179–82. 10.1080/17460913.2025.2449779

10. Quinn GA, Dyson PJ. Going to extremes: progress in exploring new environments for novel antibiotics. npj Antimicrob Resist. 2024;2:1–9. 10.1038/s44259-024-00025-8

11. Bej AK, Aislabie J, Atlas RM, editors. Polar microbiology: the ecology, biodiversity, and bioremediation potential of microorganisms in extremely cold environments. Boca Raton: 2010.

12. Medeiros W, Kralova S, Oliveira V, Ziemert N, Sehnal L. Antarctic bacterial natural products: from genomic insights to drug discovery. Nat Prod Rep. 2025; 42, 774–787. 10.1039/d4np00045e

13. Ramasamy KP, Mahawar L, Rajasabapathy R, Rajeshwari K, Miceli C, Pucciarelli S. Comprehensive insights on environmental adaptation strategies in Antarctic bacteria and biotechnological applications of cold adapted molecules. Front Microbiol. 2023;14. https://www.frontiersin.org/articles/10.3389/fmicb.2023.1197797.

14. Pearce DA. Extremophiles in Antarctica: Life at low temperatures. In: Stan-Lotter H, Fendrihan S, editors. Adaption of Microbial Life to Environmental Extremes: Novel Research Results and Application. p. 87–118. 10.1007/978-3-211-99691-1_5

15. Sayed A m., Hassan M h. a., Alhadrami H a., Hassan H m., Goodfellow M, Rateb M e. Extreme environments: microbiology leading to specialized metabolites. Journal of Applied Microbiology. 2020;128:630–57. 10.1111/jam.14386

16. Tytgat B, Verleyen E, Sweetlove M, Van den Berge K, Pinseel E, Hodgson DA, et al. Polar lake microbiomes have distinct evolutionary histories. Science Advances. American Association for the Advancement of Science; 2023;9:eade7130. 10.1126/sciadv.ade7130

17. Vyverman W, Verleyen E, Wilmotte A, Hodgson DA, Willems A, Peeters K, et al. Evidence for widespread endemism among Antarctic micro-organisms. Polar Sci. 2010;4:103–13. 10.1016/j.polar.2010.03.006

18. Waschulin V, Borsetto C, James R, Newsham KK, Donadio S, Corre C, et al. Biosynthetic potential of uncultured Antarctic soil bacteria revealed through long-read metagenomic sequencing. ISME J. 2022;16:101–11. 10.1038/s41396-021-01052-3

19. Medeiros W, Hidalgo K, Leão T, de Carvalho LM, Ziemert N, Oliveira V. Unlocking the biosynthetic potential and taxonomy of the Antarctic microbiome along temporal and spatial gradients. Microbiol Spectr. 2024;12:e0024424. 10.1128/spectrum.00244-24

20. Benaud N, Edwards RJ, Amos TG, D’Agostino PM, Gutiérrez-Chávez C, Montgomery K, et al. Antarctic desert soil bacteria exhibit high novel natural product potential, evaluated through long-read genome sequencing and comparative genomics. Environ Microbiol. 2020;23:3646–64. 10.1111/1462-2920.15300

21. Silva LJ, Crevelin EJ, Souza DT, Lacerda-Júnior GV, de Oliveira VM, Ruiz ALTG, et al. Actinobacteria from Antarctica as a source for anticancer discovery. Sci Rep. Nature Publishing Group; 2020;10:13870. 10.1038/s41598-020-69786-2

22. Schalk IJ. Bacterial siderophores: diversity, uptake pathways and applications. Nat Rev Microbiol. Nature Publishing Group; 2025;23:24–40. 10.1038/s41579-024-01090-6

23. Ribeiro M, Simões M. Advances in the antimicrobial and therapeutic potential of siderophores. Environ Chem Lett. 2019;17:1485–94. 10.1007/s10311-019-00887-9

24. Reitz ZL, Medema MH. Genome mining strategies for metallophore discovery. Current Opinion in Biotechnology. 2022;77:102757. 10.1016/j.copbio.2022.102757

25. Frei A, Zuegg J, Elliott AG, Baker M, Braese S, Brown C, et al. Metal complexes as a promising source for new antibiotics. Chem Sci. The Royal Society of Chemistry; 2020;11:2627–39. 10.1039/C9SC06460E

26. Wei Z, Gu S, Vollenweider V, Zuo Y, Li Z, Kümmerli R. Microbial siderophores for One Health. Trends in Microbiology. 2025; 10.1016/j.tim.2025.05.002

27. Saha M, Sarkar S, Sarkar B, Sharma BK, Bhattacharjee S, Tribedi P. Microbial siderophores and their potential applications: a review. Environ Sci Pollut Res. 2016;23:3984–99. 10.1007/s11356-015-4294-0

28. . Gräff ÁT, Barry SM. Siderophores as tools and treatments. npj Antimicrob Resist. Nature Publishing Group; 2024;2:47. 10.1038/s44259-024-00053-4

29. Kramer J, Özkaya Ö, Kümmerli R. Bacterial siderophores in community and host interactions. Nat Rev Microbiol. 2020;18:152–63. 10.1038/s41579-019-0284-4

30. Konstantinidis KT. Sequence-discrete species for prokaryotes and other microbes: A historical perspective and pending issues. mLife. 2023;2:341–9. 10.1002/mlf2.12088

31. Kim J, Na S-I, Kim D, Chun J. UBCG2: Up-to-date bacterial core genes and pipeline for phylogenomic analysis. J Microbiol. 2021;59:609–15. 10.1007/s12275-021-1231-4

32. Meier-Kolthoff JP, Carbasse JS, Peinado-Olarte RL, Göker M. TYGS and LPSN: a database tandem for fast and reliable genome-based classification and nomenclature of prokaryotes. Nucleic Acids Res. 2021;50:D801–7. 10.1093/nar/gkab902

33. Jain C, Rodriguez-R LM, Phillippy AM, Konstantinidis KT, Aluru S. High throughput ANI analysis of 90K prokaryotic genomes reveals clear species boundaries. Nat Commun. 2018;9:5114. 10.1038/s41467-018-07641-9

34. Blin K, Shaw S, Kloosterman AM, Charlop-Powers Z, van Wezel GP, Medema MH, et al. antiSMASH 6.0: improving cluster detection and comparison capabilities. Nucleic Acids Research. 2021; 10.1093/nar/gkab335

35. IUPAC. Compendium of Chemical Terminology, 2nd ed. (the “Gold Book”). Compiled by A. D. McNaught and A. Wilkinson. Oxford: Blackwell Scientific Publications; 1997. 10.1351/goldbook

36. Sidda JD, Song L, Poon V, Al-Bassam M, Lazos O, Buttner MJ, et al. Discovery of a family of γ-aminobutyrate ureas via rational derepression of a silent bacterial gene cluster. Chem Sci. The Royal Society of Chemistry; 2013;5:86–9. 10.1039/C3SC52536H

37. Wang M, Carver JJ, Phelan VV, Sanchez LM, Garg N, Peng Y, et al. Sharing and community curation of mass spectrometry data with Global Natural Products Social Molecular Networking. Nat Biotechnol. Nature Publishing Group; 2016;34:828–37. 10.1038/nbt.3597

38. Poynton EF, van Santen JA, Pin M, Contreras MM, McMann E, Parra J, et al. The Natural Products Atlas 3.0: extending the database of microbially derived natural products. Nucleic Acids Res. 2025;53:D691–9. 10.1093/nar/gkae1093

39. Zhao B, Moody SC, Hider RC, Lei L, Kelly SL, Waterman MR, et al. Structural Analysis of Cytochrome P450 105N1 Involved in the Biosynthesis of the Zincophore, Coelibactin. International Journal of Molecular Sciences. Molecular Diversity Preservation International; 2012;13:8500–13. 10.3390/ijms13078500

40. Bibb MJ, Janssen GR, Ward JM. Cloning and analysis of the promoter region of the erythromycin resistance gene (*ermE*) of *Streptomyces erythraeus*. Gene. 1985;38:215–26. 10.1016/0378-1119(85)90220-3

41. Zhao M, Yang Z, Li X, Liu Y, Zhang Y, Zhang M, et al. Development of Integrated Vectors with Strong Constitutive Promoters for High-Yield Antibiotic Production in Mangrove-Derived Streptomyces. Mar Drugs. 2024;22:94. 10.3390/md22020094

42. Zhou X, Wu H, Li Z, Zhou X, Bai L, Deng Z. Over-expression of UDP-glucose pyrophosphorylase increases validamycin A but decreases validoxylamine A production in *Streptomyces hygroscopicus* var. *jinggangensis* 5008. Metabolic Engineering. 2011;13:768–76. 10.1016/j.ymben.2011.10.001

43. Klumpp S, Hwa T. Growth-rate-dependent partitioning of RNA polymerases in bacteria. Proceedings of the National Academy of Sciences. Proceedings of the National Academy of Sciences; 2008;105:20245–50. 10.1073/pnas.0804953105

44. Lorenzi J-N, Thibessard A, Lioy VS, Boccard F, Leblond P, Pernodet J-L, et al. Ribosomal RNA operons define a central functional compartment in the Streptomyces chromosome. Nucleic Acids Res. 2022;50:11654–69. 10.1093/nar/gkac1076

45. Steinmetz T, Hiller W, Nett M. Amamistatins isolated from Nocardia altamirensis. Beilstein J Org Chem. Beilstein-Institut; 2022;18:360–7. 10.3762/bjoc.18.40

46. Oluwabusola ET, Adebisi OO, Reyes F, Acquah KS, De La Cruz M, Mweetwa LL, et al. Isolation and characterization of new phenolic siderophores with antimicrobial properties from Pseudomonas sp. UIAU-6B. Beilstein J Org Chem. 2021;17:2390–8. 10.3762/bjoc.17.156

47. Okujo N, Saito M, Yamamoto S, Yoshida T, Miyoshi S, Shinoda S. Structure of vulnibactin, a new polyamine-containing siderophore from Vibrio vulnificus. Biometals. 1994;7:109–16. 10.1007/BF00140480

48. Lee S, van Santen JA, Farzaneh N, Liu DY, Pye CR, Baumeister TUH, et al. NP Analyst: An Open Online Platform for Compound Activity Mapping. ACS Cent Sci. American Chemical Society; 2022;8:223–34. 10.1021/acscentsci.1c01108

49. Hoshino S, Ozeki M, Awakawa T, Morita H, Onaka H, Abe I. Catenulobactins A and B, Heterocyclic Peptides from Culturing Catenuloplanes sp. with a Mycolic Acid-Containing Bacterium. Journal of Natural Products. 2018;81:2106–10. 10.1021/acs.jnatprod.8b00261

50. Oluwabusola ET, Adebisi OO, Reyes F, Acquah KS, Cruz MDL, Mweetwa LL, et al. Isolation and characterization of new phenolic siderophores with antimicrobial properties from Pseudomonas sp. UIAU-6B. Beilstein J Org Chem. Beilstein-Institut; 2021;17:2390–8. 10.3762/bjoc.17.156

51. Park SR, Tripathi A, Wu J, Schultz PJ, Yim I, McQuade TJ, et al. Discovery of cahuitamycins as biofilm inhibitors derived from a convergent biosynthetic pathway. Nat Commun. 2016;7:10710. 10.1038/ncomms10710

52. Seyedsayamdost MR, Traxler MF, Zheng S-L, Kolter R, Clardy J. Structure and Biosynthesis of Amychelin, an Unusual Mixed-Ligand Siderophore from Amycolatopsis sp. AA4. J Am Chem Soc. American Chemical Society; 2011;133:11434–7. 10.1021/ja203577e

53. Chen Y, Unger M, Ntai I, McClure RA, Albright JC, Thomson RJ, et al. Gobichelin A and B: mixed-ligand siderophores discovered using proteomics. Med Chem Commun. 2012;4:233–8. 10.1039/C2MD20232H

54. Xiang W-X, Liu Q, Li X-M, Lu C-H, Shen Y-M. Four pairs of proline-containing cyclic dipeptides from Nocardiopsis sp. HT88, an endophytic bacterium of Mallotus nudiflorus L. Natural Product Research. 2020;34:2219–24. 10.1080/14786419.2019.1577834

55. Harish V, Periasamy M. Enantiomerically pure piperazines via NaBH4/I2 reduction of cyclic amides. Tetrahedron: Asymmetry. 2017;28:175–80. 10.1016/j.tetasy.2016.12.002

56. Janata J, Kadlcik S, Koberska M, Ulanova D, Kamenik Z, Novak P, et al. Lincosamide Synthetase—A Unique Condensation System Combining Elements of Nonribosomal Peptide Synthetase and Mycothiol Metabolism. PLoS One. 2015;10:e0118850. 10.1371/journal.pone.0118850

57. Gilchrist CLM, Chooi Y-H. clinker & clustermap.js: automatic generation of gene cluster comparison figures. Bioinformatics. 2021;37:2473–5. 10.1093/bioinformatics/btab007

58. Altschul SF, Gish W, Miller W, Myers EW, Lipman DJ. Basic local alignment search tool. J Mol Biol. 1990;215:403–10. 10.1016/S0022-2836(05)80360-2

59. Mount DW. Bioinformatics: Sequence and Genome Analysis. CSHL Press; 2004.

60. Navarro-Muñoz JC, Selem-Mojica N, Mullowney MW, Kautsar SA, Tryon JH, Parkinson EI, et al. A computational framework to explore large-scale biosynthetic diversity. Nat Chem Biol. 2020;16:60–8. 10.1038/s41589-019-0400-9

61. Cary SC, McDonald IR, Barrett JE, Cowan DA. On the rocks: the microbiology of Antarctic Dry Valley soils. Nat Rev Microbiol. 2010;8:129–38. 10.1038/nrmicro2281

62. Kieser T. Practical Streptomyces Genetics. John Innes Foundation; 2000.

63. Andrews SC, Robinson AK, Rodríguez-Quiñones F. Bacterial iron homeostasis. FEMS Microbiol Rev. 2003;27:215–37. 10.1016/S0168-6445(03)00055-X

64. Miethke M, Marahiel MA. Siderophore-based iron acquisition and pathogen control. Microbiol Mol Biol Rev. 2007;71:413–51. 10.1128/MMBR.00012-07

65. Ge L, Seah SYK. Heterologous Expression, Purification, and Characterization of an l-Ornithine N5-Hydroxylase Involved in Pyoverdine Siderophore Biosynthesis in Pseudomonas aeruginosa. Journal of Bacteriology. 2006;188:7205–10. 10.1128/jb.00949-06

66. Koglin A, Walsh CT. Structural insights into nonribosomal peptide enzymatic assembly lines. Nat Prod Rep. 2009;26:987–1000. 10.1039/B904543K

67. Cheng Y, Yang R, Lyu M, Wang S, Liu X, Wen Y, et al. IdeR, a DtxR Family Iron Response Regulator, Controls Iron Homeostasis, Morphological Differentiation, Secondary Metabolism, and the Oxidative Stress Response in Streptomyces avermitilis. Applied and Environmental Microbiology. 2018;84:e01503–18. 10.1128/AEM.01503-18

68. Abanoz-Seçgin B, Otur Ç, Okay S, Kurt-Kızıldoğan A. The regulatory role of Fur-encoding SCLAV_3199 in iron homeostasis in Streptomyces clavuligerus. Gene. 2023;878:147594. 10.1016/j.gene.2023.147594

69. Santos TMA, Lammers MG, Zhou M, Sparks IL, Rajendran M, Fang D, et al. Small Molecule Chelators Reveal That Iron Starvation Inhibits Late Stages of Bacterial Cytokinesis. ACS Chem Biol. 2018;13:235–46. 10.1021/acschembio.7b00560

70. Vlček V, Pospíšilová Ľ, Uhlík P. Mineralogy and chemical composition of Cryosols and Andosols in Antarctica. Soil and Water Research. Soil and Water Research; 2018;13:61–73. 10.17221/231/2016-SWR

71. Ramos-Alonso L, Romero AM, Martínez-Pastor MT, Puig S. Iron Regulatory Mechanisms in Saccharomyces cerevisiae. Front Microbiol. Frontiers; 2020;11. 10.3389/fmicb.2020.582830

72. Dancis A, Klausner RD, Hinnebusch AG, Barriocanal JG. Genetic evidence that ferric reductase is required for iron uptake in Saccharomyces cerevisiae. Mol Cell Biol. 1990;10:2294–301. 10.1128/mcb.10.5.2294-2301.1990

73. Gerwien F, Safyan A, Wisgott S, Brunke S, Kasper L, Hube B. The Fungal Pathogen Candida glabrata Does Not Depend on Surface Ferric Reductases for Iron Acquisition. Front Microbiol. 2017;8:1055. 10.3389/fmicb.2017.01055

74. Kumar K, Askari F, Sahu MS, Kaur R. Candida glabrata: A Lot More Than Meets the Eye. Microorganisms. 2019;7:39. 10.3390/microorganisms7020039

75. Knight SAB, Vilaire G, Lesuisse E, Dancis A. Iron Acquisition from Transferrin by Candida albicans Depends on the Reductive Pathway. Infect Immun. 2005;73:5482–92. 10.1128/IAI.73.9.5482-5492.2005

76. Almeida RS, Brunke S, Albrecht A, Thewes S, Laue M, Jr JEE, et al. The Hyphal-Associated Adhesin and Invasin Als3 of Candida albicans Mediates Iron Acquisition from Host Ferritin. PLOS Pathogens. 2008;4:e1000217. 10.1371/journal.ppat.1000217

77. Roy U, Kornitzer D. Heme-iron acquisition in fungi. Curr Opin Microbiol. 2019;52:77–83. 10.1016/j.mib.2019.05.006

78. Kuznets G, Vigonsky E, Weissman Z, Lalli D, Gildor T, Kauffman SJ, et al. A Relay Network of Extracellular Heme-Binding Proteins Drives C. albicans Iron Acquisition from Hemoglobin. PLOS Pathogens. 2014 ;10:e1004407. 10.1371/journal.ppat.1004407

79. Heymann P, Gerads M, Schaller M, Dromer F, Winkelmann G, Ernst JF. The siderophore iron transporter of Candida albicans (Sit1p/Arn1p) mediates uptake of ferrichrome-type siderophores and is required for epithelial invasion. Infect Immun. 2002;70:5246–55. 10.1128/IAI.70.9.5246-5255.2002

80. Chen C, Pande K, French SD, Tuch BB, Noble SM. An iron homeostasis regulatory circuit with reciprocal roles in Candida albicans commensalism and pathogenesis. Cell Host Microbe. 2011;10:118–35. 10.1016/j.chom.2011.07.005

81. Kosman DJ. Molecular mechanisms of iron uptake in fungi. Mol Microbiol. 2003;47:1185–97. 10.1046/j.1365-2958.2003.03368.x

82. Roy U, Yaish S, Weissman Z, Pinsky M, Dey S, Horev G, et al. Ferric reductase-related proteins mediate fungal heme acquisition. eLife;11:e80604. 10.7554/eLife.80604

83. Hider RC, Kong X. Chemistry and biology of siderophores. Nat Prod Rep. The Royal Society of Chemistry; 2010; 27: 637–57. 10.1039/B906679A

84. Boik J. Natural Compounds in Cancer Therapy. Oregon Medical Press; 2001.

85. Roman M, Nývlt D, Davies BJ, Braucher R, Jennings SJA, Břežný M, et al. Accelerated retreat of northern James Ross Island ice streams (Antarctic Peninsula) in the Early-Middle Holocene induced by buoyancy response to postglacial sea level rise. Earth and Planetary Science Letters. 2024 ;641:118803. 10.1016/j.epsl.2024.118803

86. Rego A, Raio F, Martins TP, Ribeiro H, Sousa AGG, Séneca J, et al. Actinobacteria and Cyanobacteria Diversity in Terrestrial Antarctic Microenvironments Evaluated by Culture-Dependent and Independent Methods. Front Microbiol. Frontiers; 2019;10. 10.3389/fmicb.2019.01018

87. Hayakawa M, Nonomura H. Humic acid-vitamin agar, a new medium for the selective isolation of soil actinomycetes. Journal of Fermentation Technology. 1987;65:501–9. 10.1016/0385-6380(87)90108-7

88. Shirling EB, Gottlieb D. Methods for characterization of Streptomyces species. International Journal of Systematic Bacteriology. 1966;16:313–40.

89. Waksman SA. Strain specificity and production of antibiotic substances. x. characterization and classification of species within the Streptomyces griseus group. Proc Natl Acad Sci U S A. 1959;45:1043–7. 10.1073/pnas.45.7.1043

90. Sapkota A, Thapa A, Budhathoki A, Sainju M, Shrestha P, Aryal S. Isolation, Characterization, and Screening of Antimicrobial-Producing Actinomycetes from Soil Samples. International Journal of Microbiology. 2020;2020:2716584. 10.1155/2020/2716584

91. Loy A, Lehner A, Lee N, Adamczyk J, Meier H, Ernst J, et al. Oligonucleotide Microarray for 16S rRNA Gene-Based Detection of All Recognized Lineages of Sulfate-Reducing Prokaryotes in the Environment. Appl Environ Microbiol. 2002;68:5064–81. 10.1128/AEM.68.10.5064-5081.2002

92. Chalita M, Kim YO, Park S, Oh H-S, Cho JH, Moon J, et al. EzBioCloud: a genome-driven database and platform for microbiome identification and discovery. International Journal of Systematic and Evolutionary Microbiology. 2024 ;74:006421. 10.1099/ijsem.0.006421

93. Kumar S, Stecher G, Li M, Knyaz C, Tamura K. MEGA X: Molecular Evolutionary Genetics Analysis across Computing Platforms. Mol Biol Evol. 2018;35:1547–9. 10.1093/molbev/msy096

94. Edgar RC. MUSCLE: multiple sequence alignment with high accuracy and high throughput. Nucleic Acids Res. 2004;32:1792–7. 10.1093/nar/gkh340

95. Kolmogorov M, Yuan J, Lin Y, Pevzner PA. Assembly of long, error-prone reads using repeat graphs. Nat Biotechnol. 2019;37:540–6. 10.1038/s41587-019-0072-8

96. Gurevich A, Saveliev V, Vyahhi N, Tesler G. QUAST: quality assessment tool for genome assemblies. Bioinformatics. 2013;29:1072–5. 10.1093/bioinformatics/btt086

97. Parks DH, Imelfort M, Skennerton CT, Hugenholtz P, Tyson GW. CheckM: assessing the quality of microbial genomes recovered from isolates, single cells, and metagenomes. Genome Res. 2015;25:1043–55. 10.1101/gr.186072.114

98. Meier-Kolthoff JP, Göker M. TYGS is an automated high-throughput platform for state-of-the-art genome-based taxonomy. Nature Communications. Nature Publishing Group; 2019;10:2182. 10.1038/s41467-019-10210-3

99. Schwengers O, Jelonek L, Dieckmann MA, Beyvers S, Blom J, Goesmann A. Bakta: rapid and standardized annotation of bacterial genomes via alignment-free sequence identification. Microb Genom. 2021;7:000685. 10.1099/mgen.0.000685

100. Blin K, Shaw S, Augustijn HE, Reitz ZL, Biermann F, Alanjary M, et al. antiSMASH 7.0: new and improved predictions for detection, regulation, chemical structures and visualisation. Nucleic Acids Res. 2023;51:W46–50. 10.1093/nar/gkad344

101. Sayers EW, Bolton EE, Brister JR, Canese K, Chan J, Comeau DC, et al. Database resources of the national center for biotechnology information. Nucleic Acids Res. 2022;50:D20–6. 10.1093/nar/gkab1112

102. Blum M, Andreeva A, Florentino LC, Chuguransky SR, Grego T, Hobbs E, et al. InterPro: the protein sequence classification resource in 2025. Nucleic Acids Res. 2025;53:D444–56. 10.1093/nar/gkae1082

103. Zdouc MM, Blin K, Louwen NLL, Navarro J, Loureiro C, Bader CD, et al. MIBiG 4.0: advancing biosynthetic gene cluster curation through global collaboration. Nucleic Acids Res. 2025;53:D678–90. 10.1093/nar/gkae1115

104. Shannon P, Markiel A, Ozier O, Baliga NS, Wang JT, Ramage D, et al. Cytoscape: a software environment for integrated models of biomolecular interaction networks. Genome Res. 2003;13:2498–504. 10.1101/gr.1239303

105. Zettler J, Xia H, Burkard N, Kulik A, Grond S, Heide L, et al. New aminocoumarins from the rare actinomycete Catenulispora acidiphila DSM 44928: identification, structure elucidation, and heterologous production. Chembiochem. 2014;15:612–21. 10.1002/cbic.201300712

106. Vignolle A, Zehl M, Kirkegaard RH, Vignolle GA, Zotchev SB. Secondary Metabolite Biosynthesis Potential of Streptomyces Spp. from the Rhizosphere of Leontopodium nivale Subsp. alpinum. ACS Omega. 2025;10:7163–71. 10.1021/acsomega.4c10476

107. van Santen JA, Poynton EF, Iskakova D, McMann E, Alsup TA, Clark TN, et al. The Natural Products Atlas 2.0: a database of microbially-derived natural products. Nucleic Acids Res. 2022;50:D1317–23. 10.1093/nar/gkab941

108. Bogdanov A, Salib MN, Chase AB, Hammerlindl H, Muskat MN, Luedtke S, et al. Small molecule in situ resin capture provides a compound first approach to natural product discovery. Nat Commun. 2024;15:5230. 10.1038/s41467-024-49367-x

109. van Dissel D, van Wezel GP. Morphology-driven downscaling of Streptomyces lividans to micro-cultivation. Antonie van Leeuwenhoek. 2018;111:457–69. 10.1007/s10482-017-0967-7

110. Worth RM, Espina L. ScanGrow: Deep Learning-Based Live Tracking of Bacterial Growth in Broth. Front Microbiol. Frontiers; 2022;13. 10.3389/fmicb.2022.900596

111. Agbangba CE, Sacla Aide E, Honfo H, Glèlè Kakai R. On the use of post-hoc tests in environmental and biological sciences: A critical review. Heliyon. 2024;10:e25131. 10.1016/j.heliyon.2024.e25131

112. Schwyn B, Neilands JB. Universal chemical assay for the detection and determination of siderophores. Analytical Biochemistry. 1987;160:47–56. 10.1016/0003-2697(87)90612-9

113. Arora NK, Verma M. Modified microplate method for rapid and efficient estimation of siderophore produced by bacteria. 3 Biotech. 2017;7:381. 10.1007/s13205-017-1008-y

114. Payne SM. Iron acquisition in microbial pathogenesis. Trends in Microbiology. 1993;1:66–9. 10.1016/0966-842X(93)90036-Q

115. Kuzyk SB, Hughes E, Yurkov V. Discovery of Siderophore and Metallophore Production in the Aerobic Anoxygenic Phototrophs. Microorganisms.2021;9:959. 10.3390/microorganisms9050959

116. Patel PR, Shaikh SS, Sayyed RZ. Modified chrome azurol S method for detection and estimation of siderophores having affinity for metal ions other than iron. Environmental Sustainability. 2018;1:81–7. 10.1007/s42398-018-0005-3

117. CLSI. CLSI Antimicrobial and Antifungal Susceptibility Testing Resources (M02, M07, M45, M44, M51. Clinical and Laboratory Standards, Wayne, PA.; 2015.

118. Berkow EL, Lockhart SR, Ostrosky-Zeichner L. Antifungal Susceptibility Testing: Current Approaches. Clin Microbiol Rev. 2020;33:e00069–19. 10.1128/CMR.00069-19

119. Sambrook J, Fritsch EF, Maniatis T. Molecular Cloning: A Laboratory Manual 3rd Edn. Cold Spring Harbor, NY: CSHL Press; 1989.

120. Sioud S, Aigle B, Karray-Rebai I, Smaoui S, Bejar S, Mellouli L. Integrative Gene Cloning and Expression System for Streptomyces sp. US 24 and Streptomyces sp. TN 58 Bioactive Molecule Producing Strains. J Biomed Biotechnol. 2009;2009:464986. 10.1155/2009/464986

121. Zotchev S, Haugan K, Sekurova O, Sletta H, Ellingsen TE, Valla S. Identification of a gene cluster for antibacterial polyketide-derived antibiotic biosynthesis in the nystatin producer Streptomyces noursei ATCC 11455. Microbiology (Reading). 2000;146 (Pt 3):611–9. 10.1099/00221287-146-3-611

122. Flett F, Mersinias V, Smith CP. High efficiency intergeneric conjugal transfer of plasmid DNA from Escherichia coli to methyl DNA-restricting streptomycetes. FEMS Microbiology Letters. 1997;155:223–9. 10.1111/j.1574-6968.1997.tb13882.x

123. Andrews S. FastQC: A Quality Control Tool for High Throughput Sequence Data [Online]. Available online at: http://www.bioinformatics.babraham.ac.uk/projects/fastqc/.

124. Ewels P, Magnusson M, Lundin S, Käller M. MultiQC: summarize analysis results for multiple tools and samples in a single report. Bioinformatics. 2016;32:3047–8. 10.1093/bioinformatics/btw354

125. Li H, Handsaker B, Wysoker A, Fennell T, Ruan J, Homer N, et al. The Sequence Alignment/Map format and SAMtools. Bioinformatics. 2009;25:2078–9. 10.1093/bioinformatics/btp352

126. Love MI, Huber W, Anders S. Moderated estimation of fold change and dispersion for RNA-seq data with DESeq2. Genome Biol. 2014;15:550. 10.1186/s13059-014-0550-8

