## Supplementary Materials for "Kineochelins - a novel group of siderophores from an Antarctic bacterium"

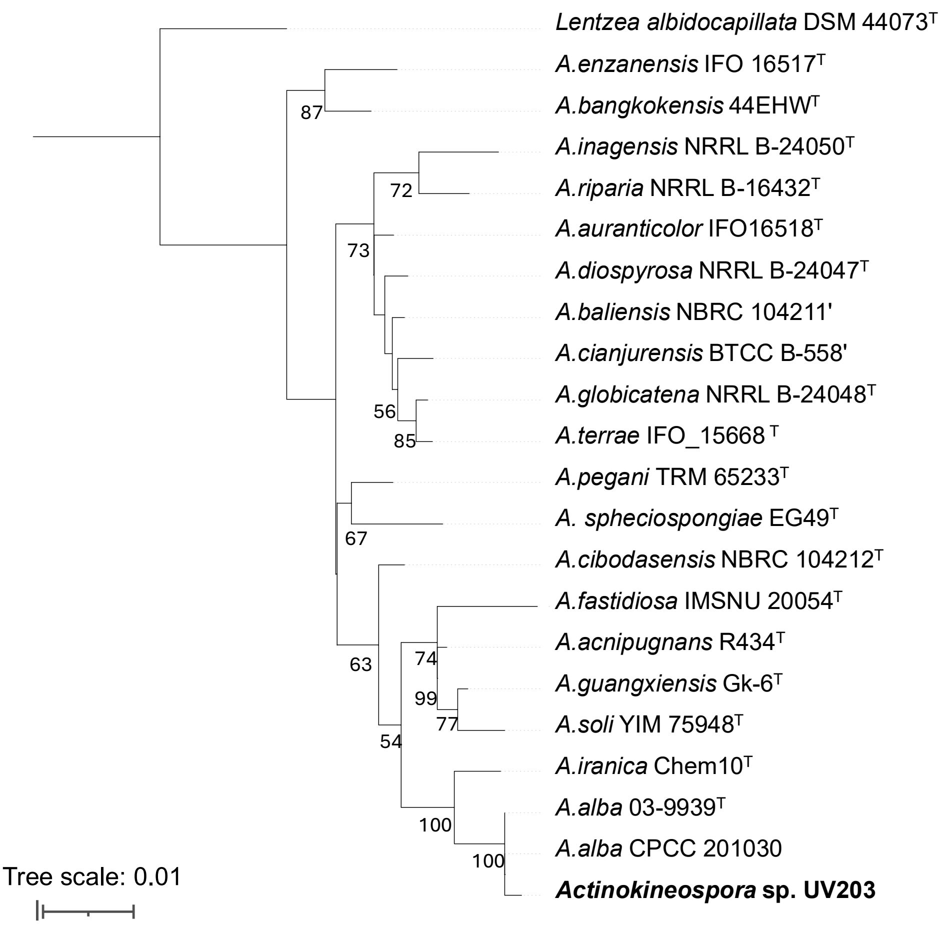


**Supplementary Figure S1.** 16S rRNA gene phylogeny showing the placement of *Actinokineospora* sp. UV203 within the genus; scale bar indicates 0.01 substitutions per site. Numbers at nodes denote the bootstrap values (>50%) for branch points based on 1,000 replications.

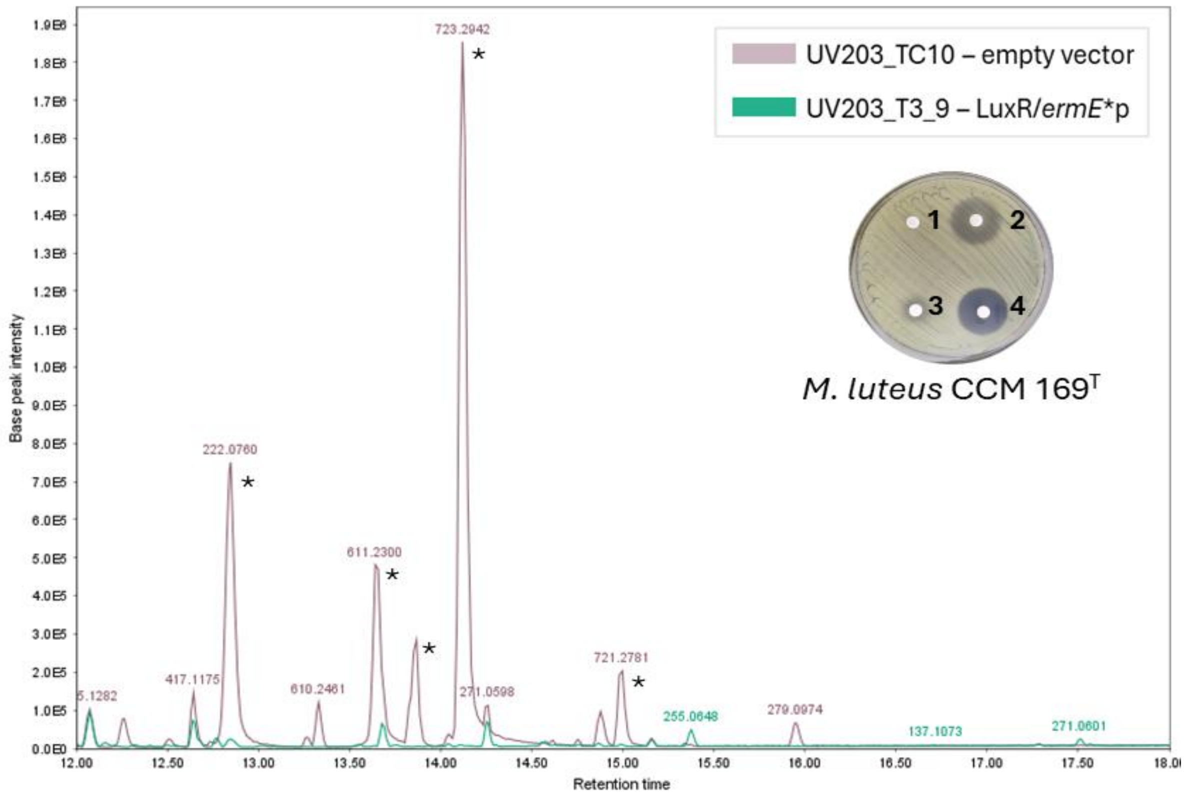


**Supplementary Figure S2.** Base peak chromatograms showing retention time regions with differential metabolite production between UV203 mutants. Inhibition assays on medium with *Micrococcus luteus* as the indicator strain. Zones: (1) negative control; (2) UV203 wild-type; (3) UV203_T3_9 mutant (ermEp with LuxR promoter); (4) UV203_TC10 mutant (empty vector). Asterisks indicate kineochelin congeners.

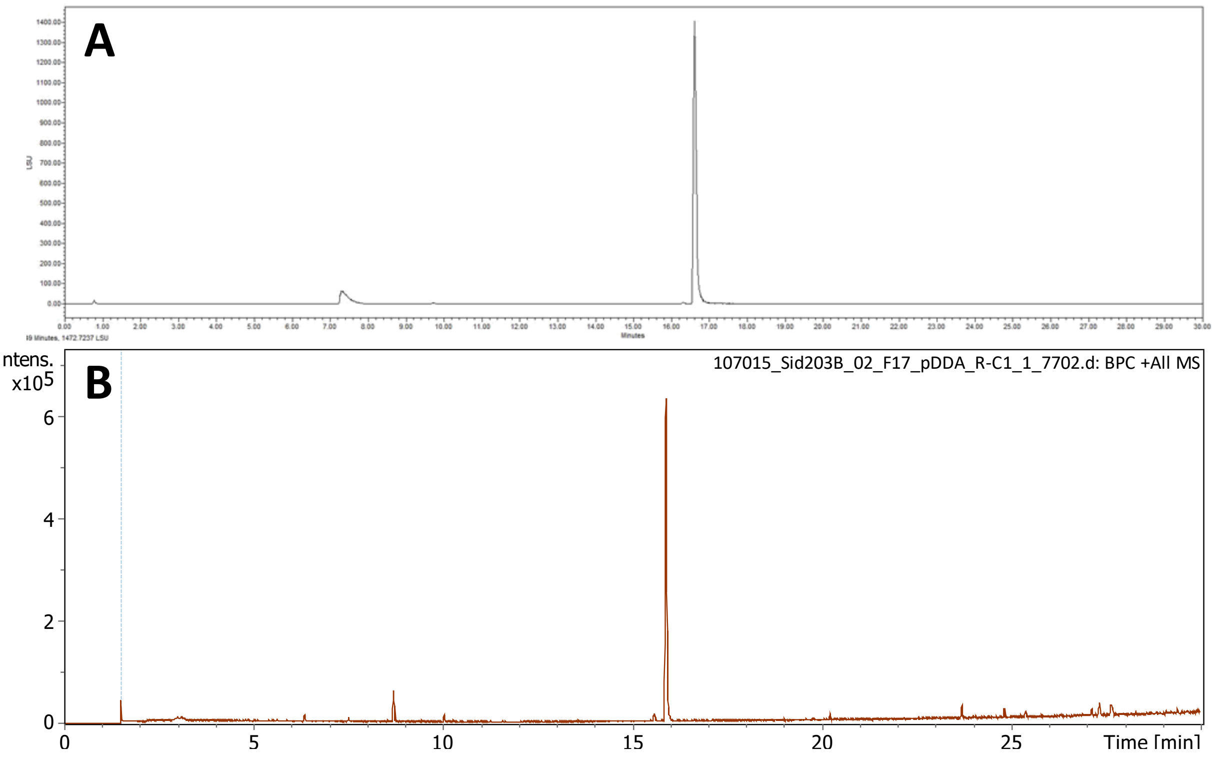


**Supplementary Figure S3.** Purity of isolated kineochelin $E_1$. UHPLC-ELSD chromatogram (A) and UHPLC-MS base peak chromatogram (B) of fraction 17 (2.0 mg) containing kineochelin $E_1$ (**1**) as main compound.

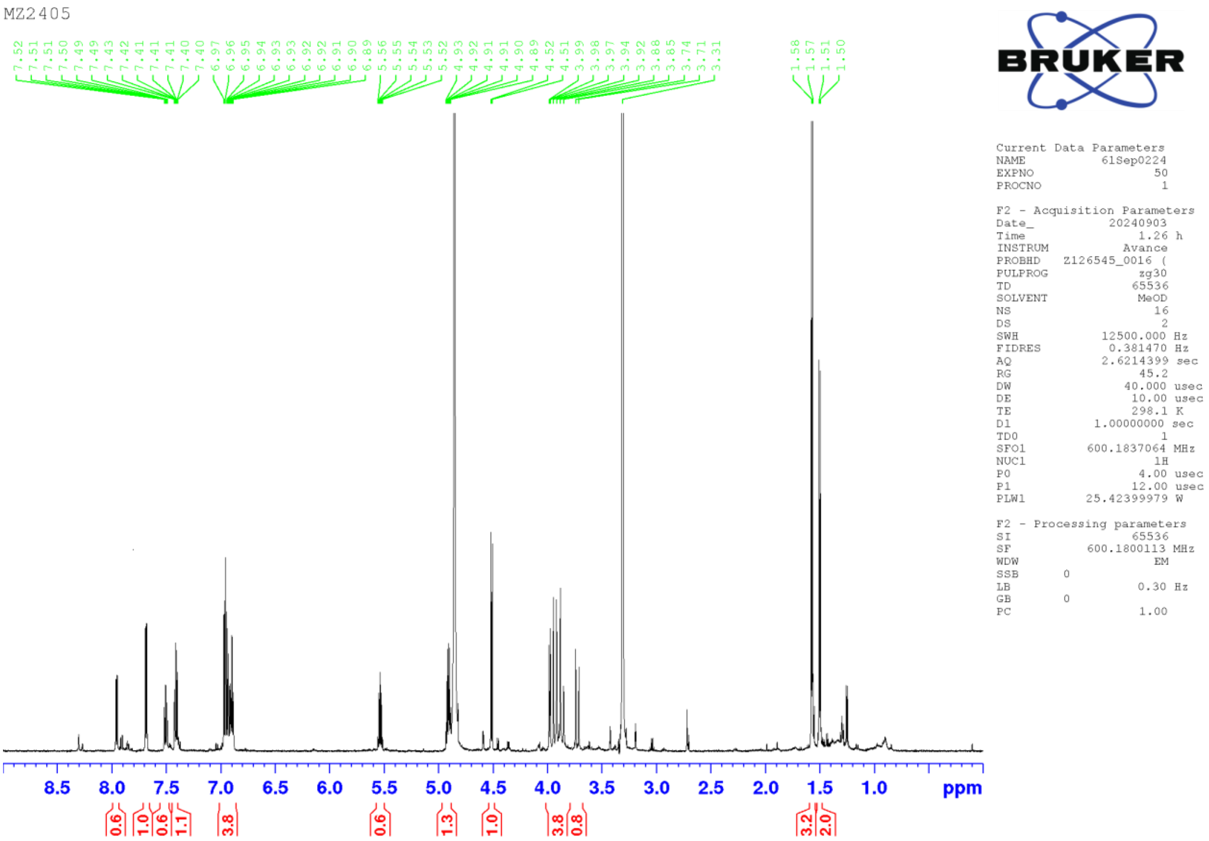


**Supplementary Figure S4.** $^1$H NMR spectrum of kineochelin E$_1$ (**1**) in $CD_3OH$ at 600 MHz.

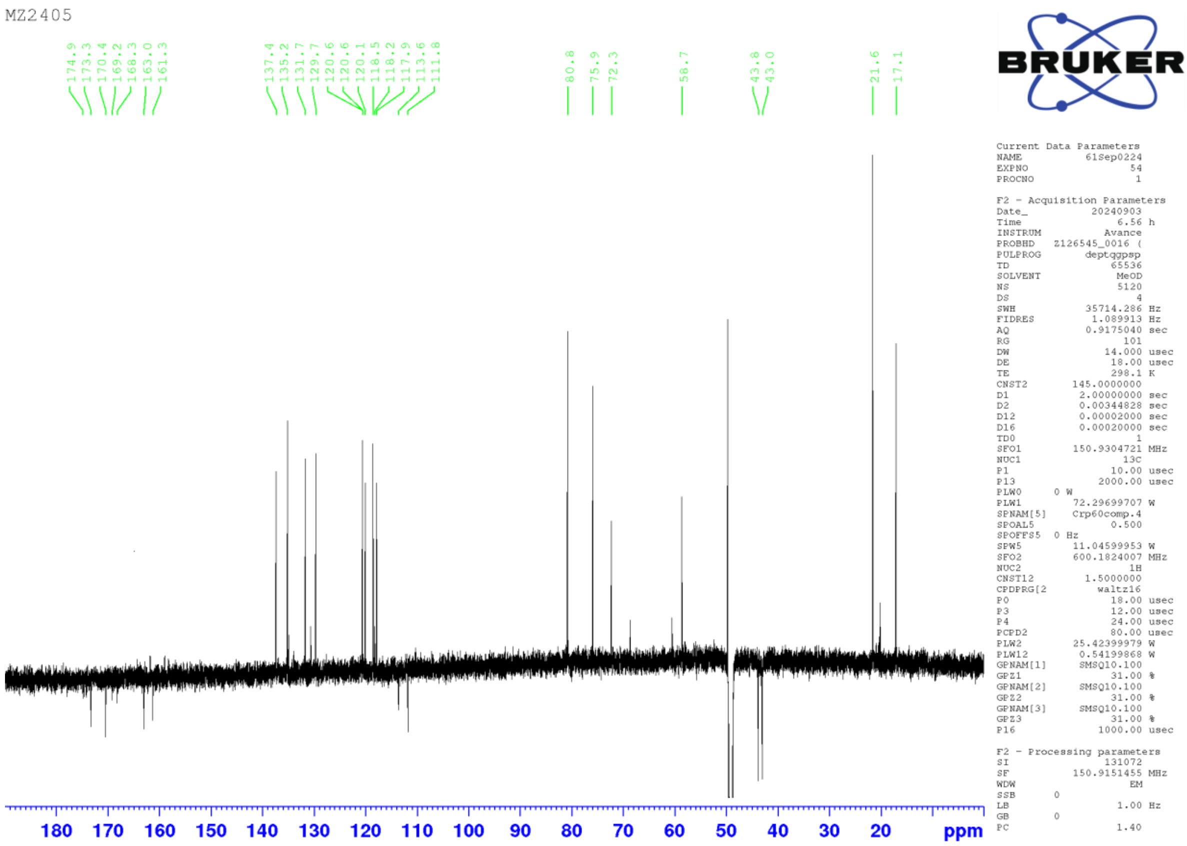


**Supplementary Figure S5.** $^{13}C$ (DEPTq) NMR spectrum of kineochelin $E_1$ (**1**) in $CD_3OH$ at 151 MHz.

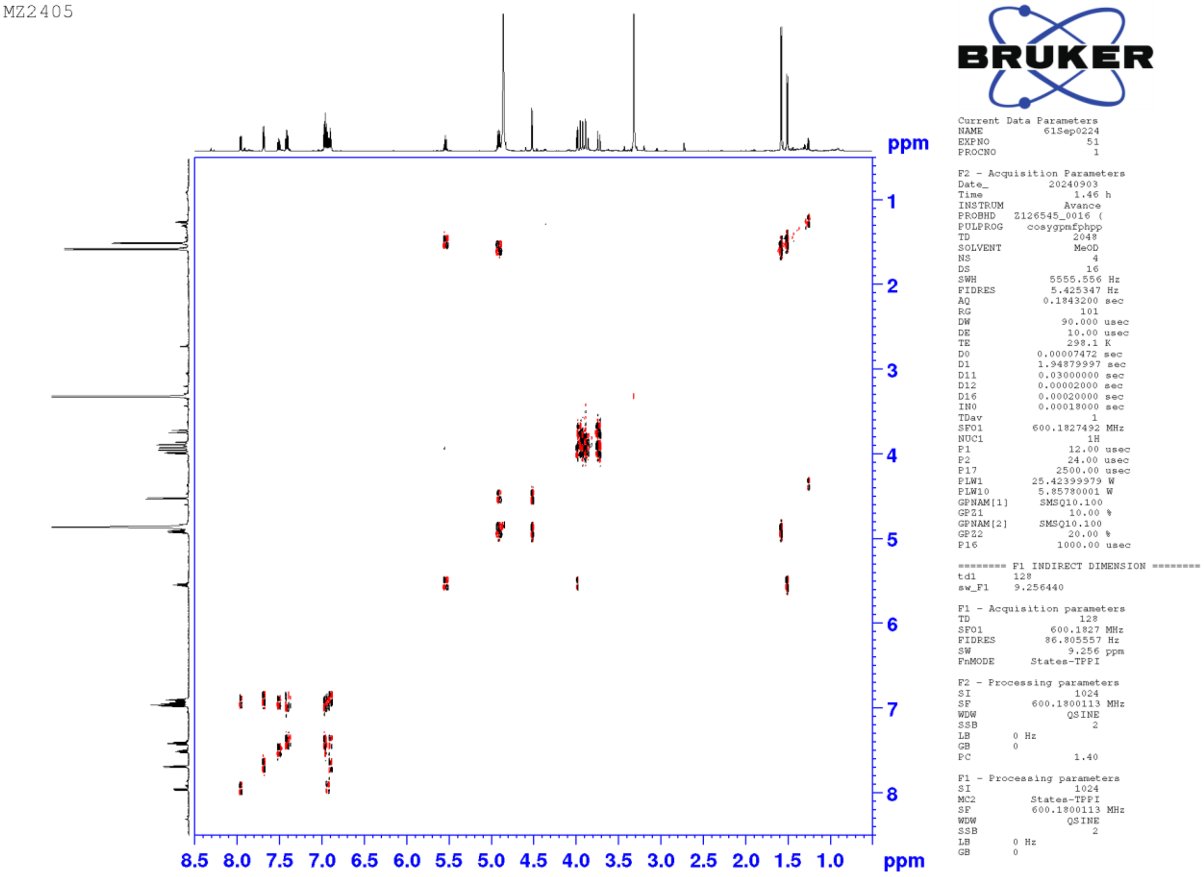


**Supplementary Figure S6.** COSY spectrum of kineochelin $E_1$ (**1**) in $CD_3OH$ at 600 MHz.

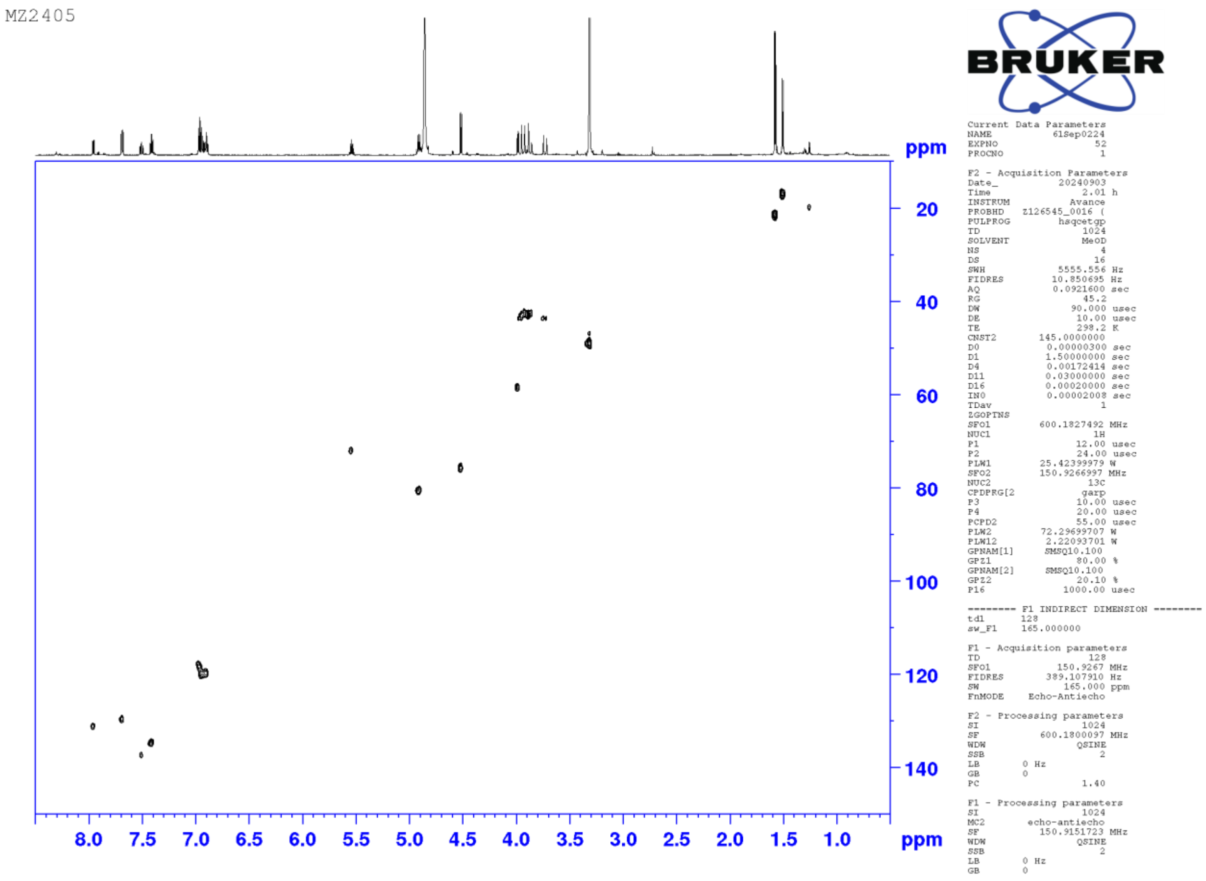


**Supplementary Figure S7.** HSQC spectrum of kineochelin $E_1$ (**1**) in $CD_3OH$ at 600 MHz.

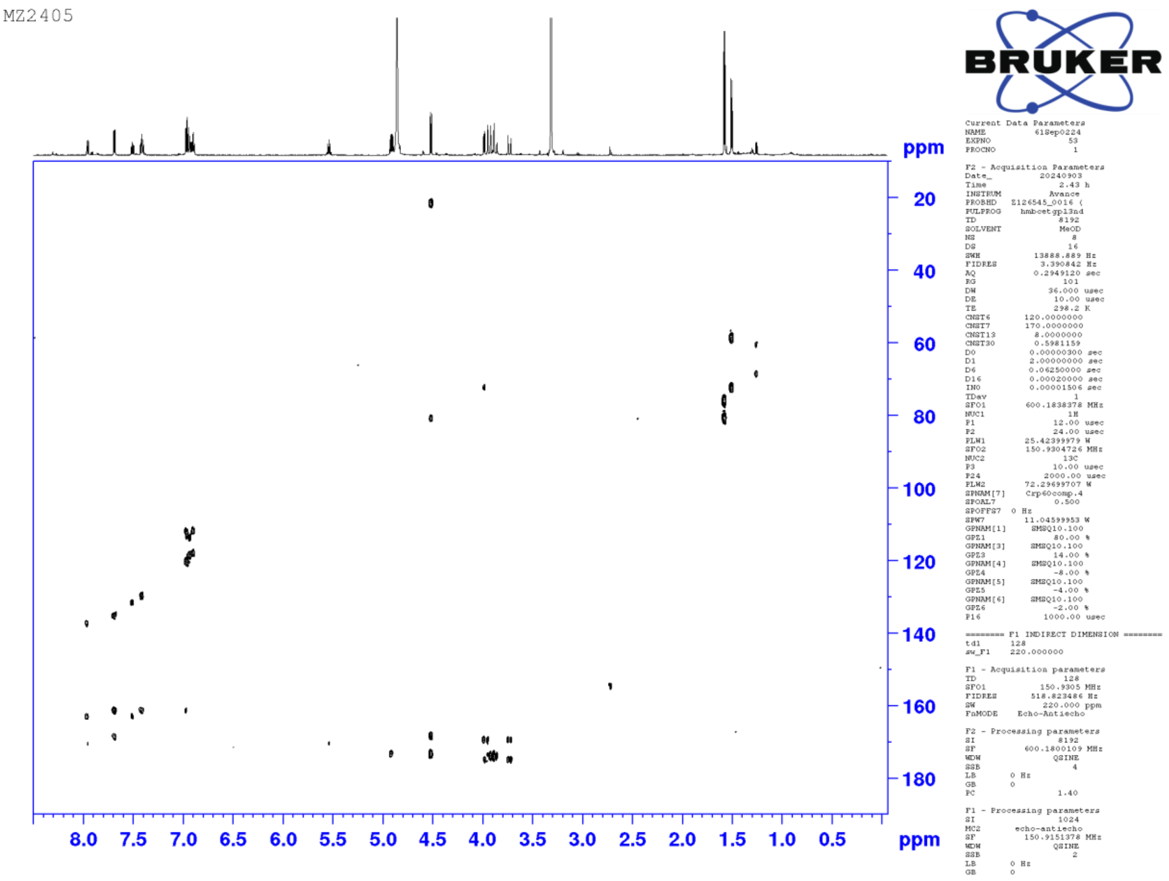


**Supplementary Figure S8.** HMBC spectrum of kineochelin $E_1$ (**1**) in $CD_3OH$ at 600 MHz.

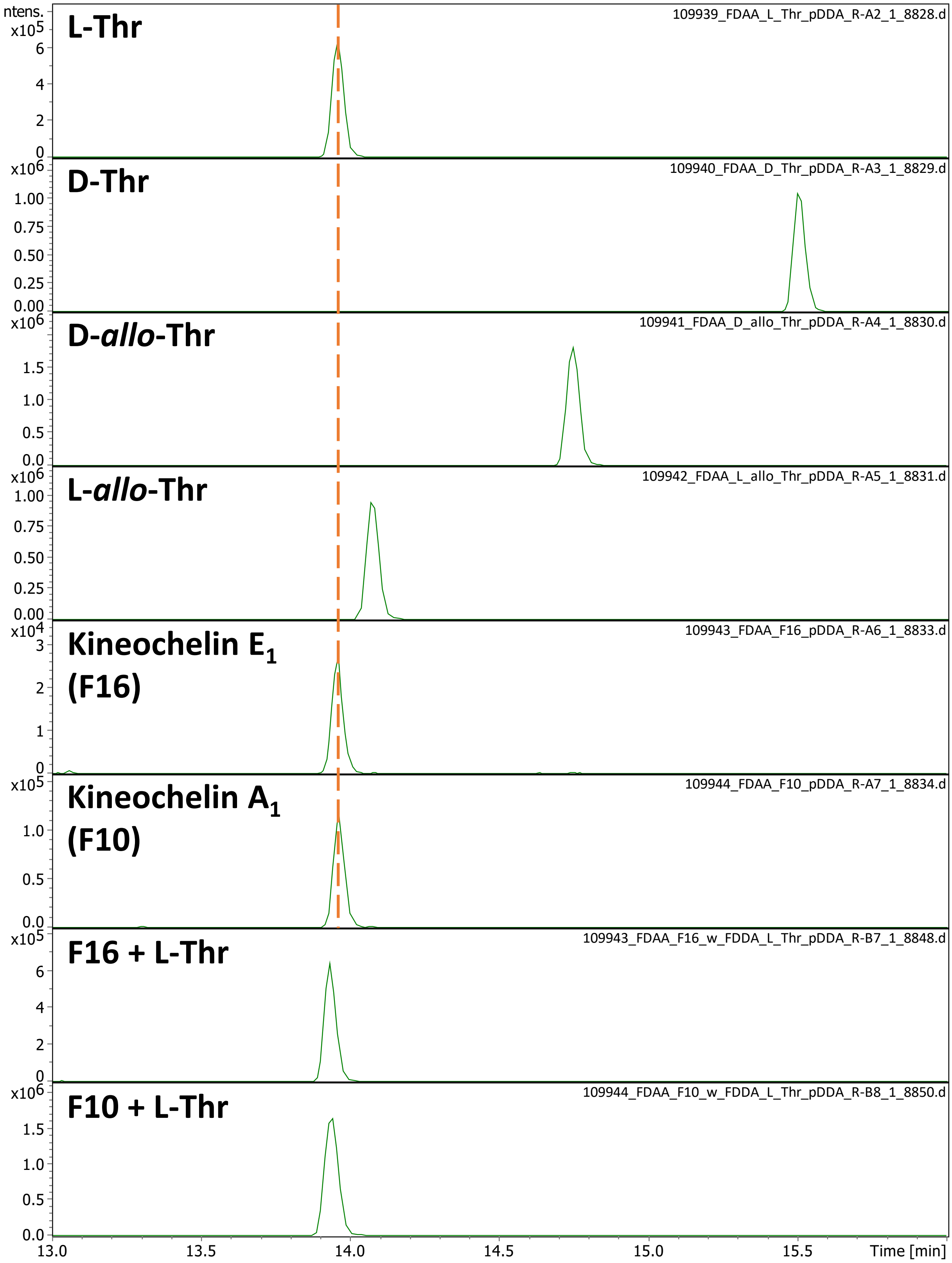


**Supplementary Figure S9.** Marfey's analysis of kineochelin E$_1$ and kineochelin A$_1$. Extracted ion chromatograms (*m/z* 372.1150±0.0050) showing the signals for L-FDAA-derivatised free amino acids L-Thr, D-Thr, D-*allo*-Thr, and L-*allo*-Thr, as well as of L-Thr in the hydrolysed fraction F16, containing mainly kineochelin E$_1$, and hydrolysed fraction F10, containing mainly kineochelin A$_1$. The bottom two EICs show the standard addition experiments confirming presence of pure L-Thr in kineochelin E$_1$ and kineochelin A$_1$.

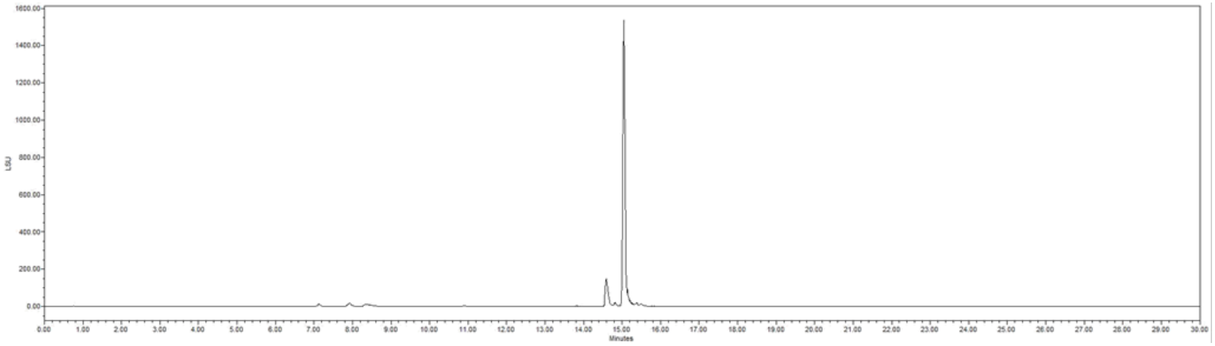


**Supplementary Figure S10.** Purity of isolated kineochelin A$_1$. UHPLC-ELSD chromatogram of fraction 11 (5.8 mg) containing kineochelin A$_1$ (2) as main compound.

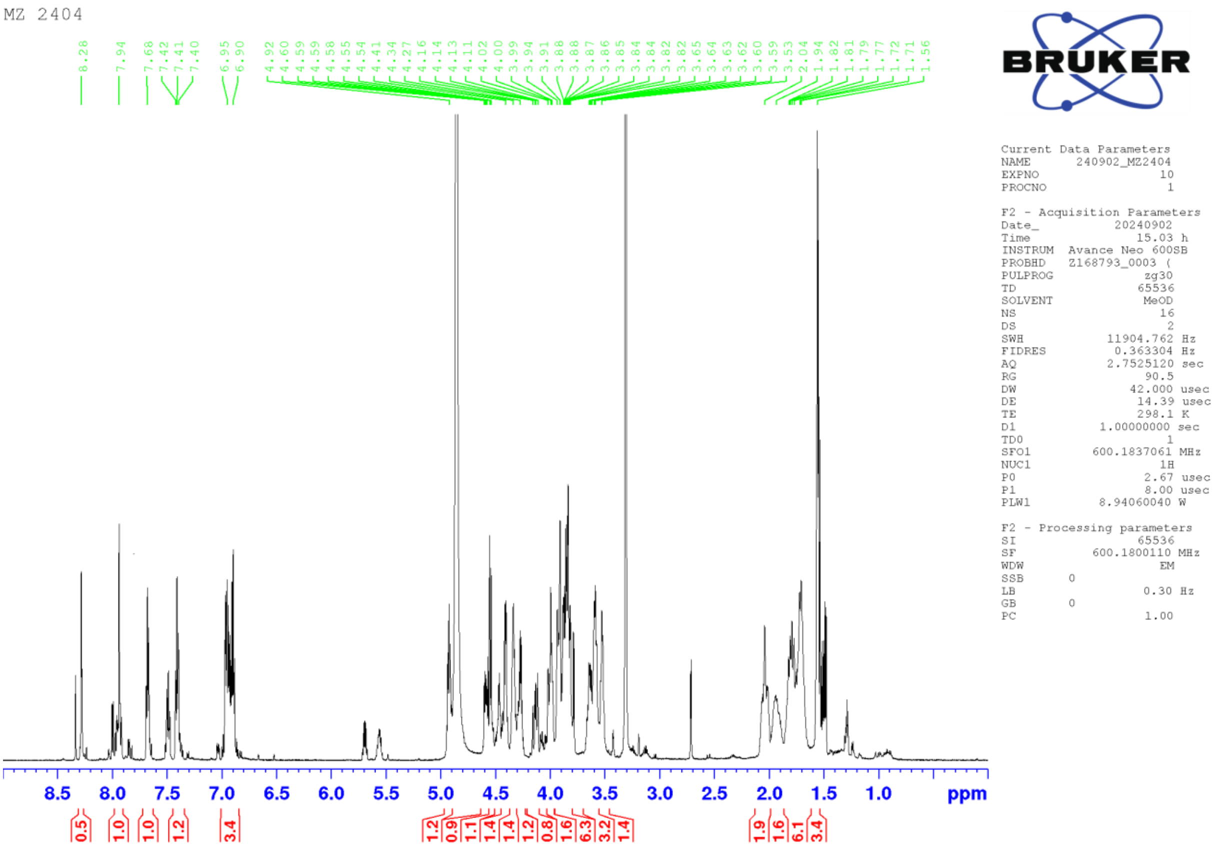


**Supplementary Figure S11.** [1]H NMR spectrum of kineochelin A$_1$ (**2**) in $CD_3OH$ at 600 MHz.

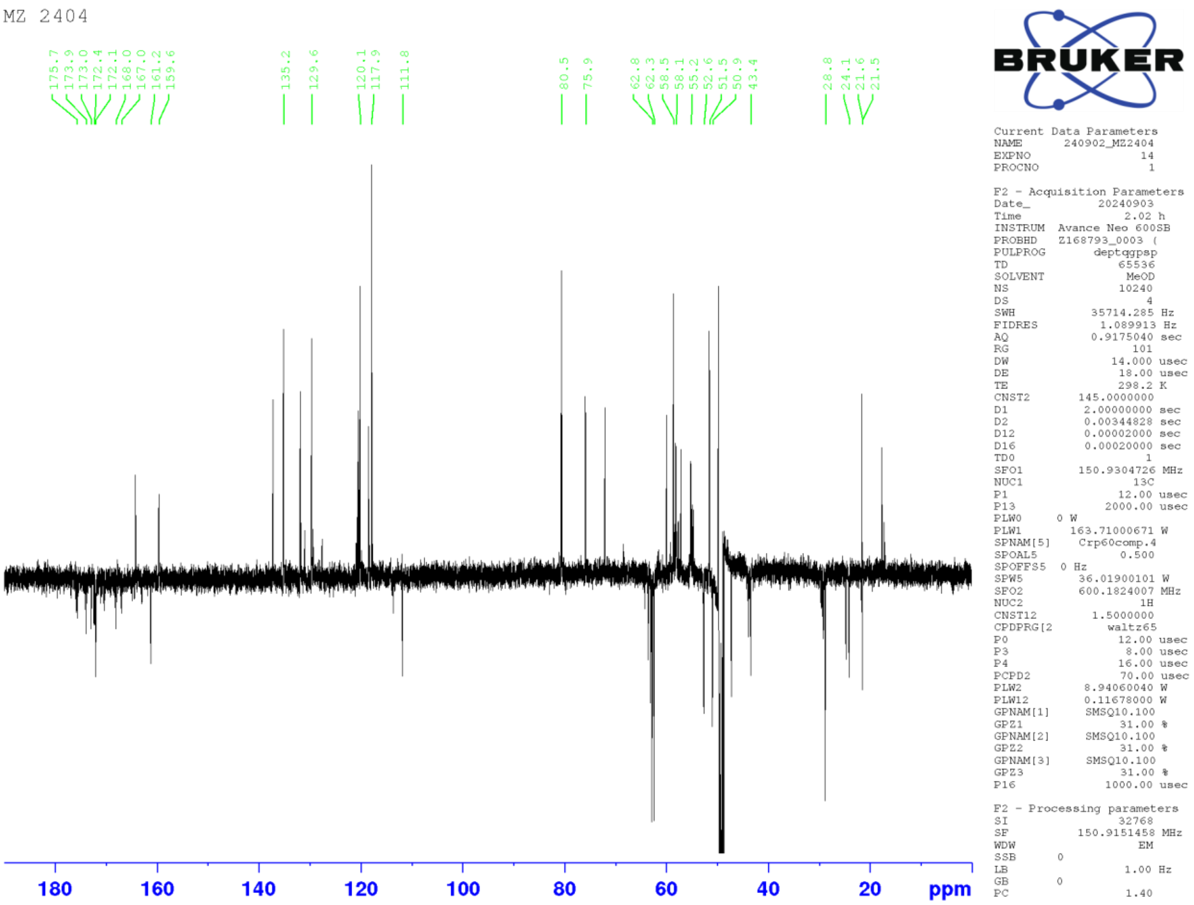


**Supplementary Figure S12.** $^{13}C$ (DEPTq) NMR spectrum of kineochelin $A_1$ (**2**) in $CD_3OH$ at 151 MHz.

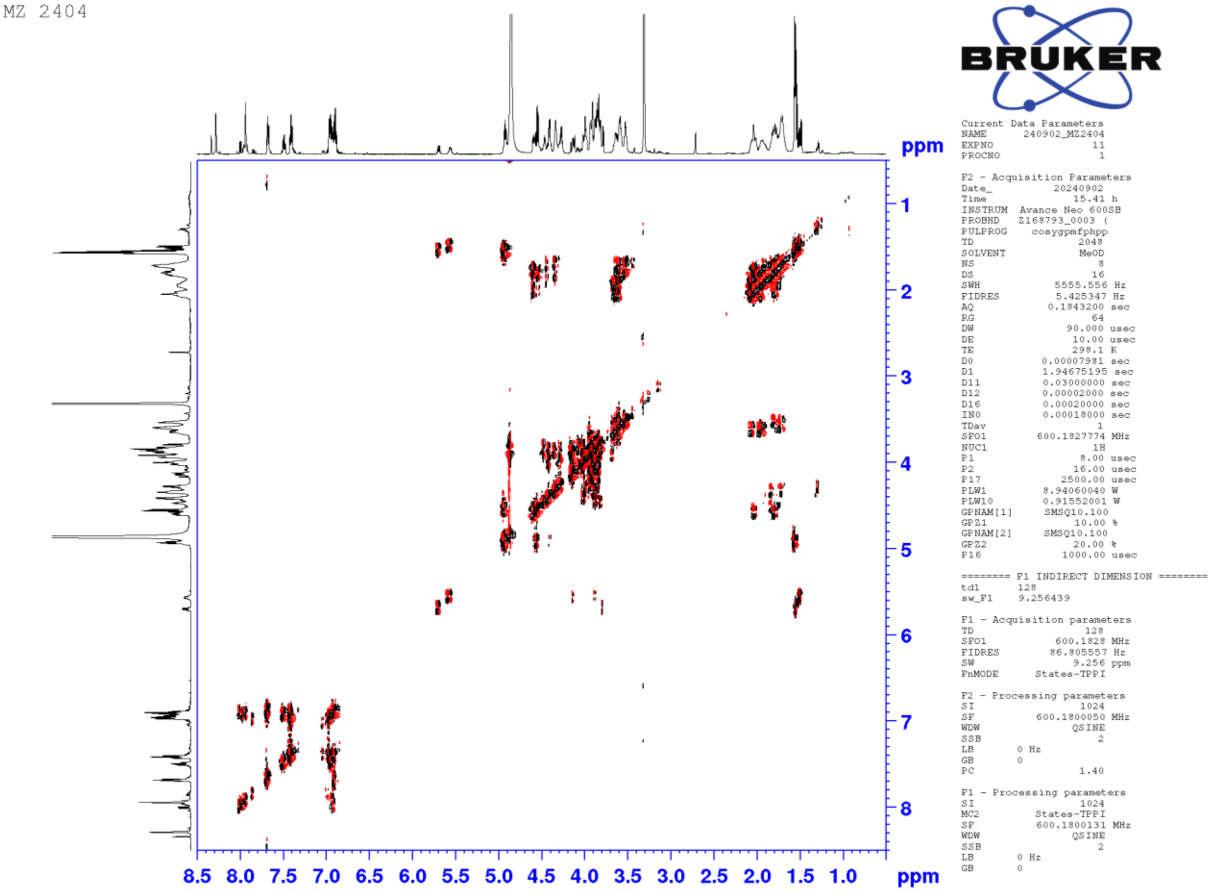


**Supplementary Figure S13.** COSY spectrum of kineochelin $A_1$ (**2**) in $CD_3OH$ at 600 MHz.

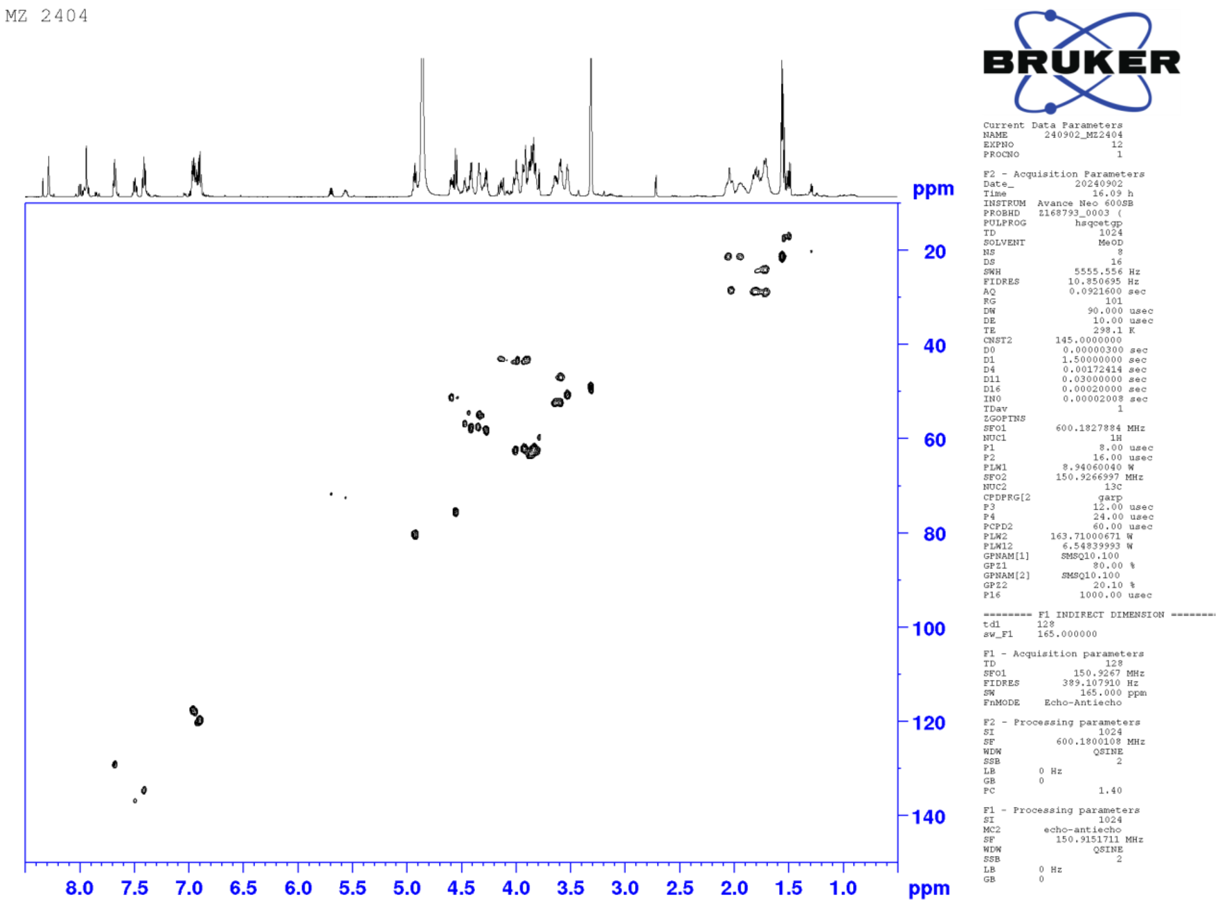


**Supplementary Figure S14.** HSQC spectrum of kineochelin $A_1$ (**2**) in $CD_3OH$ at 600 MHz.

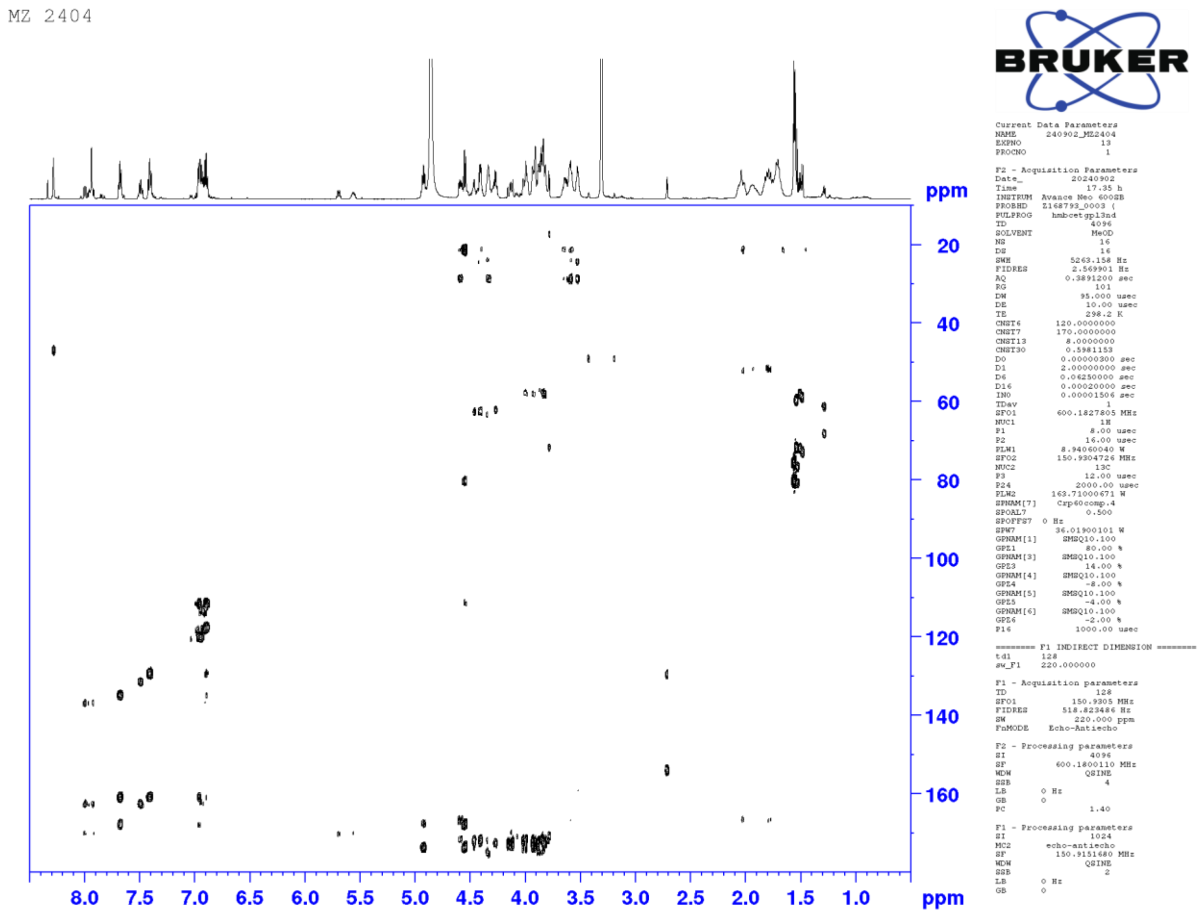


**Supplementary Figure S15.** HMBC spectrum of kineochelin $A_1$ (**2**) in $CD_3OH$ at 600 MHz.

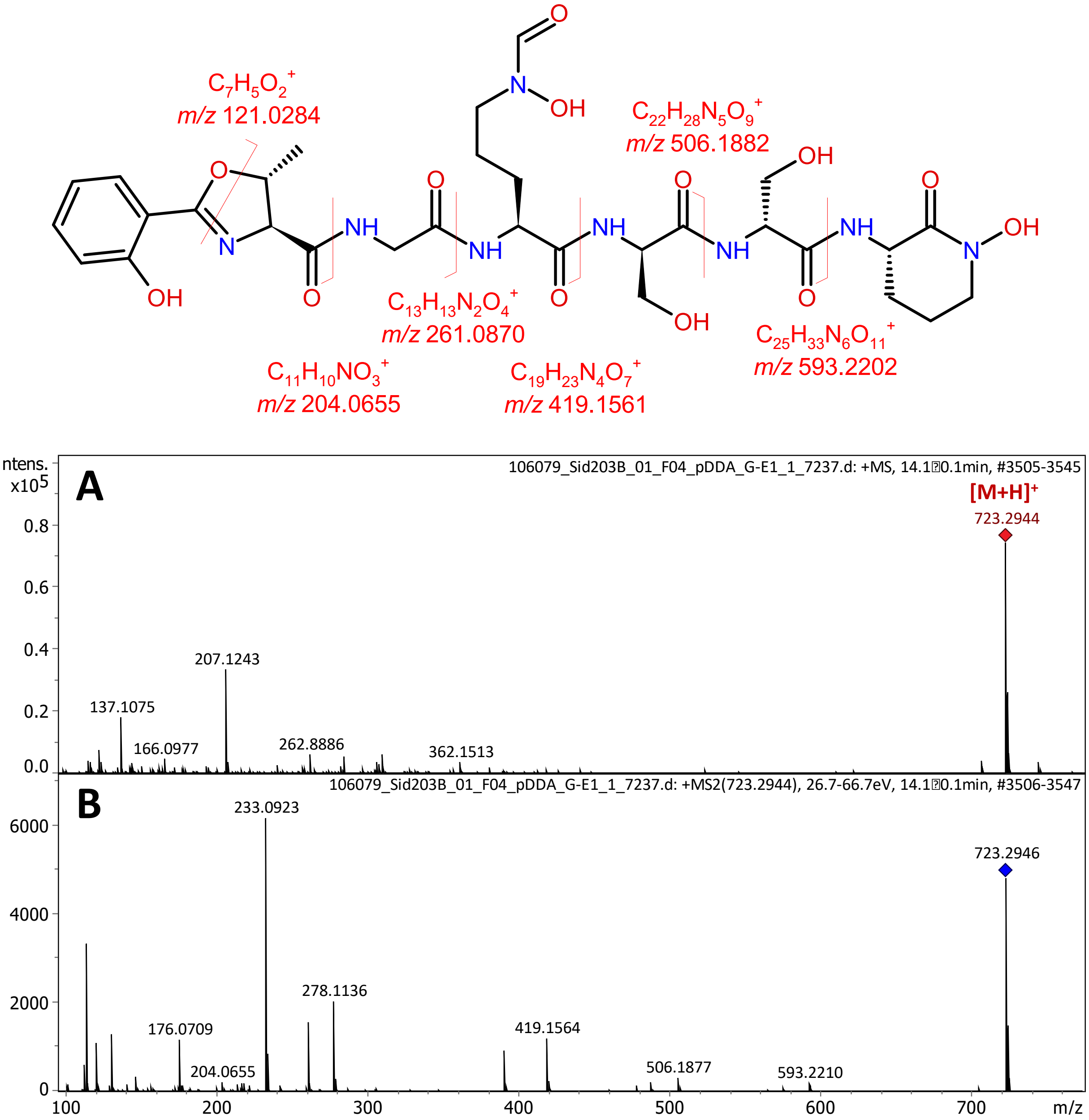


**Supplementary Figure S16.** High resolution ESI-Qq-TOF mass spectrum of kineochelin $A_1$ (A) and high resolution MS/MS spectrum of its $[M+H]^+$ ion (B).

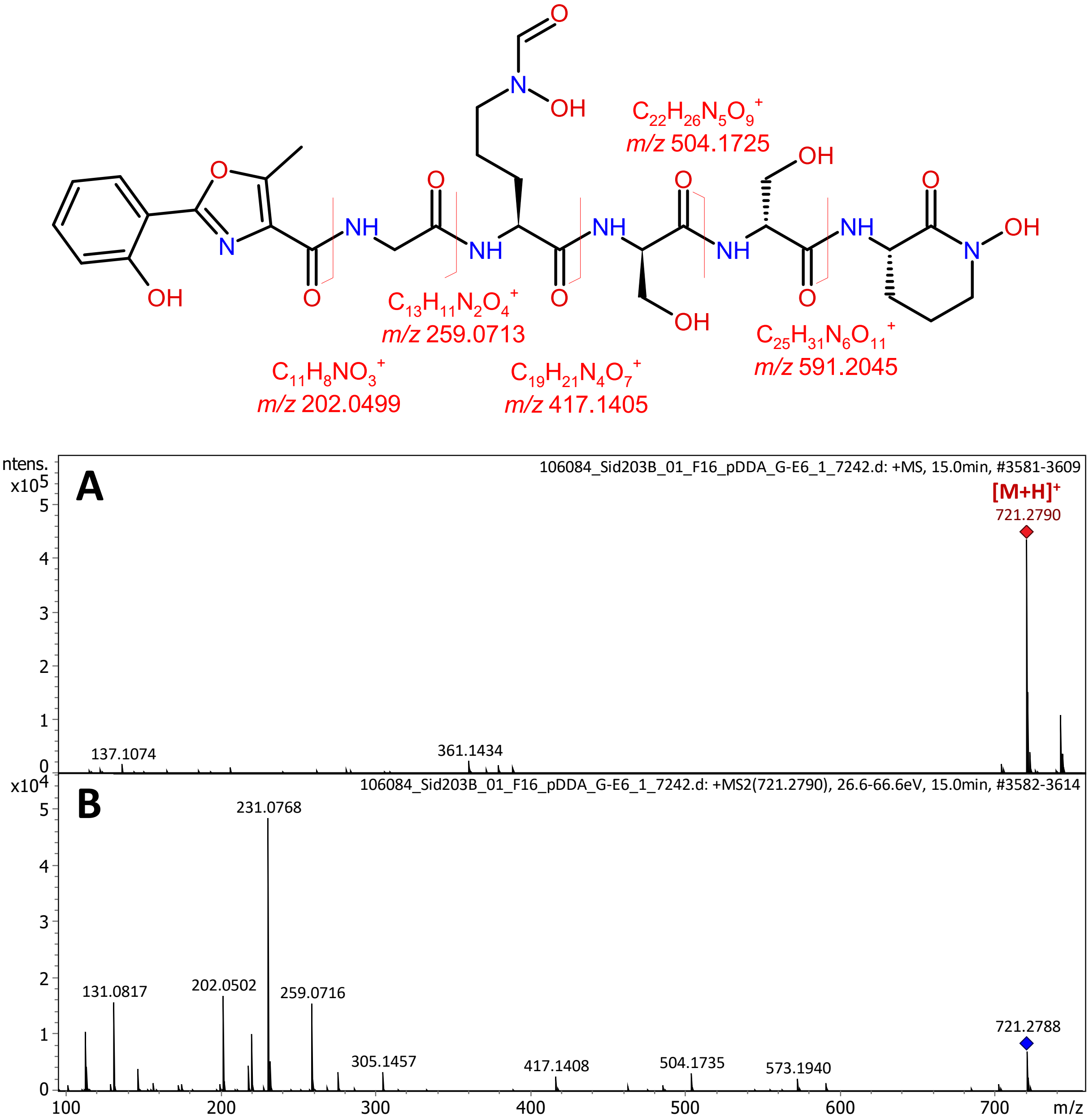


**Supplementary Figure S17.** High resolution ESI-Qq-TOF mass spectrum of kineochelin $A_2$ (A) and high resolution MS/MS spectrum of its $[M+H]^+$ ion (B).

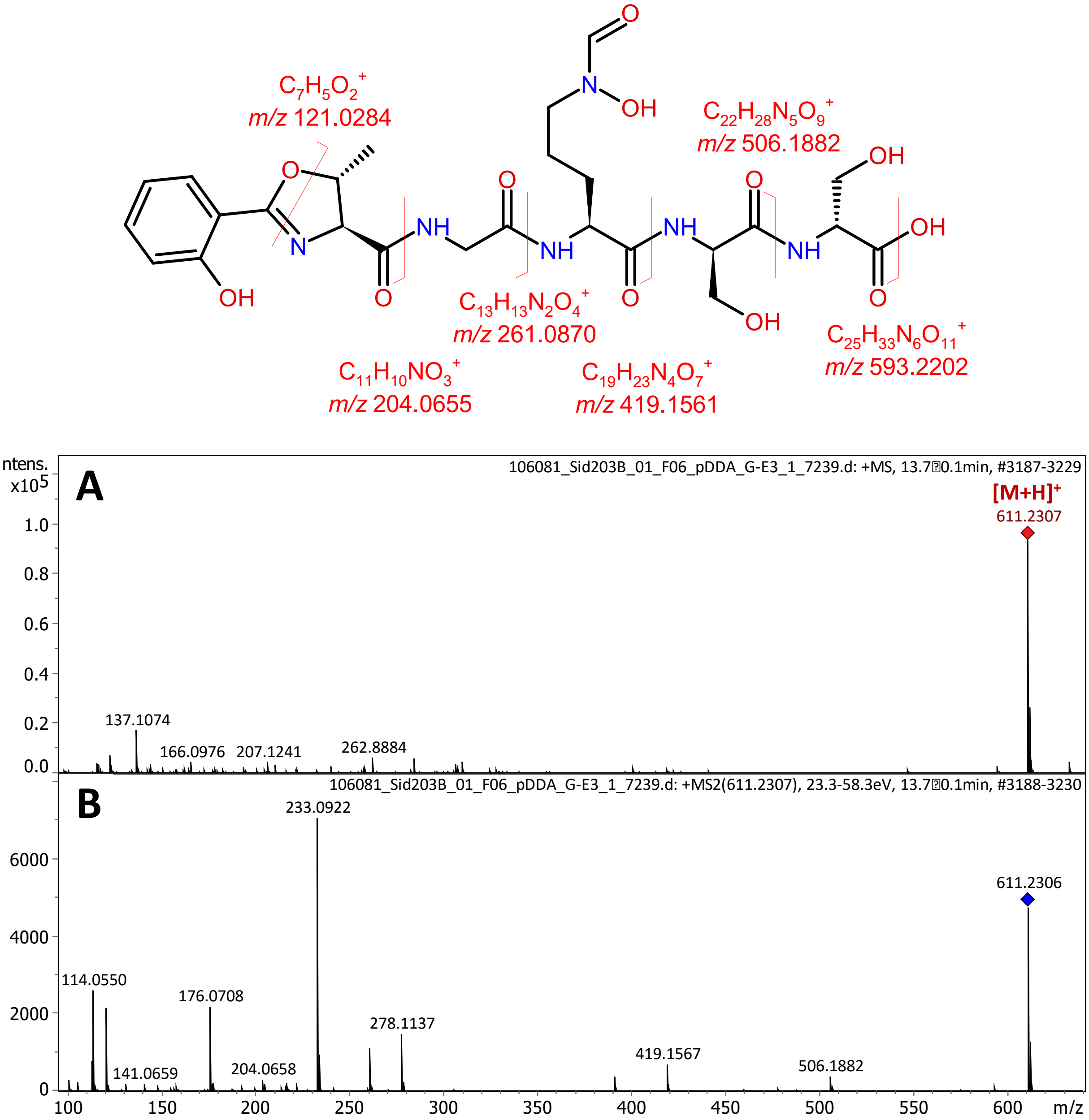


**Supplementary Figure S18.** High resolution ESI-Qq-TOF mass spectrum of kineochelin $B_1$ (A) and high resolution MS/MS spectrum of its $[M+H]^+$ ion (B).

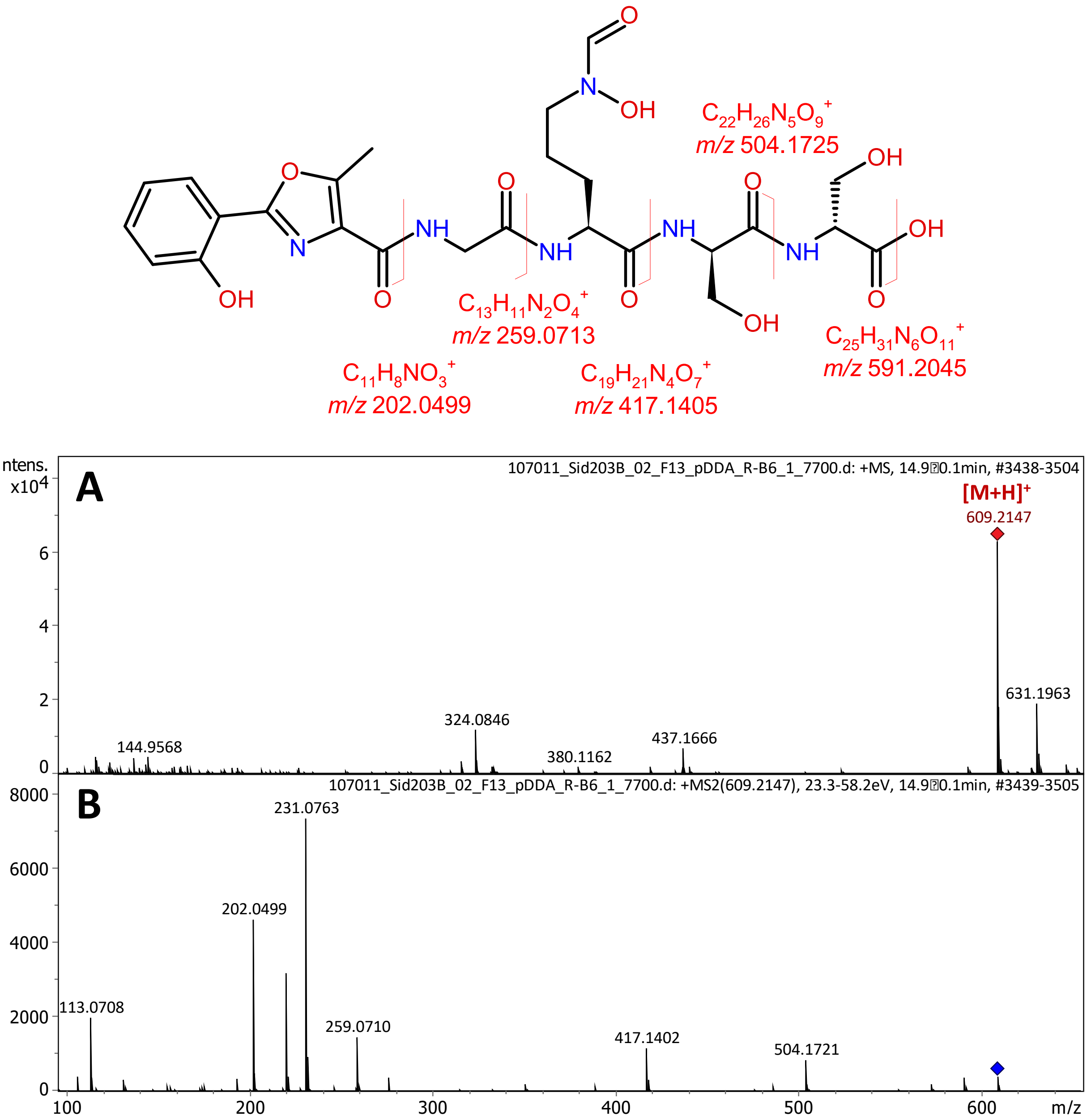


**Supplementary Figure S19.** High resolution ESI-Qq-TOF mass spectrum of kineochelin $B_2$ (A) and high resolution MS/MS spectrum of its $[M+H]^+$ ion (B).

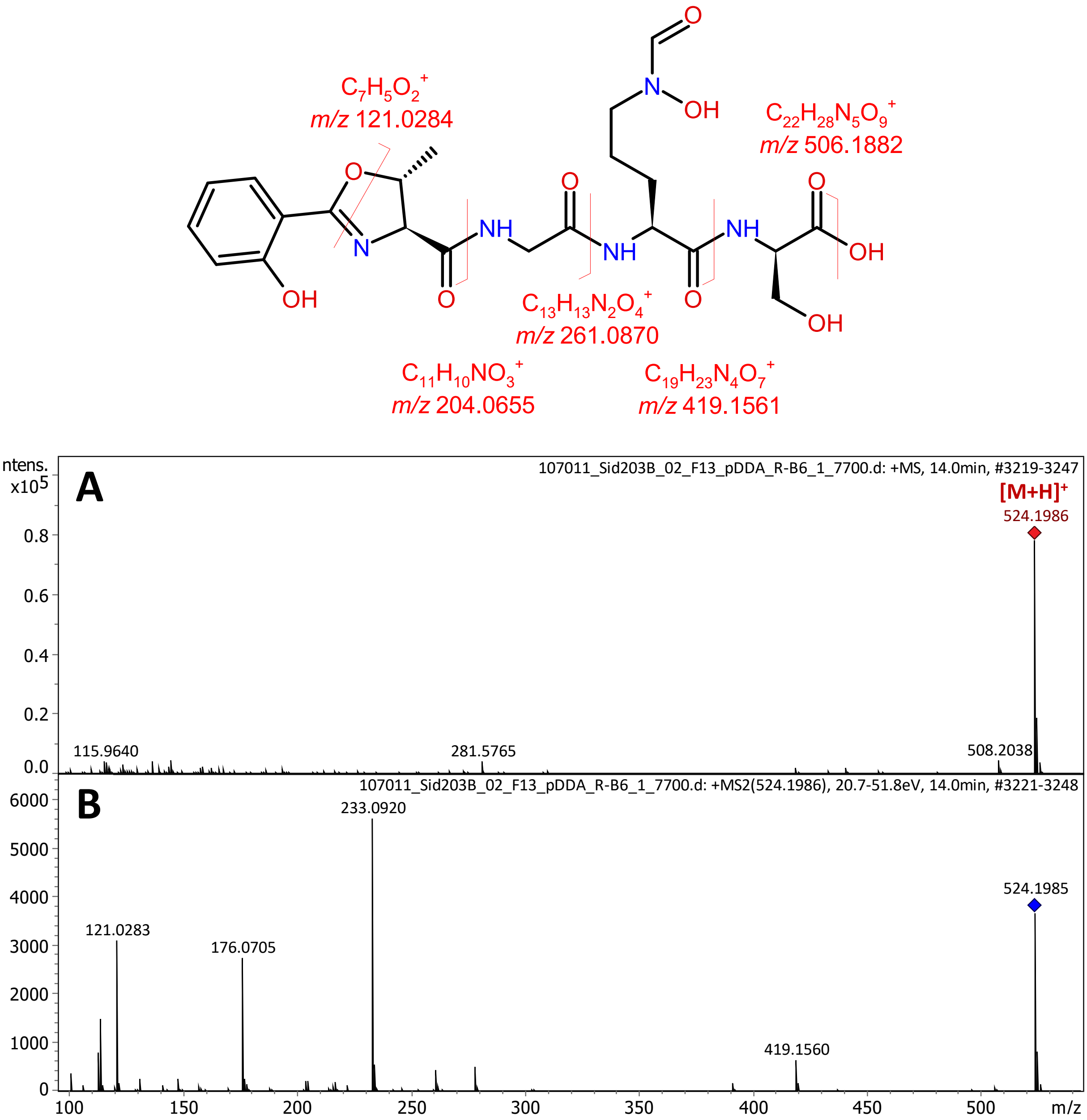


**Supplementary Figure S20.** High resolution ESI-Qq-TOF mass spectrum of kineochelin $C_1$ (A) and high resolution MS/MS spectrum of its $[M+H]^+$ ion (B).

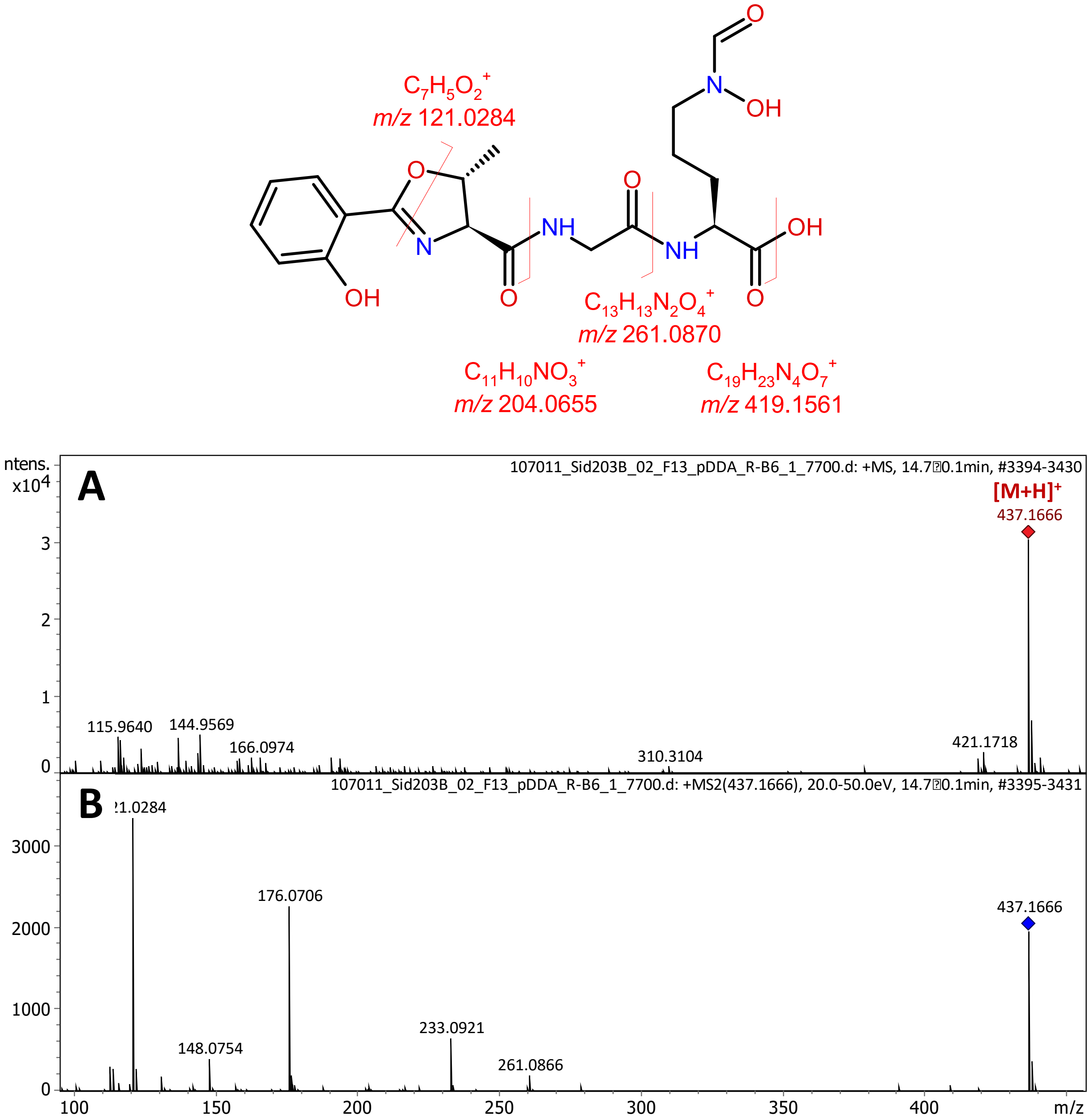


**Supplementary Figure S21.** High resolution ESI-Qq-TOF mass spectrum of kineochelin $D_1$ (A) and high resolution MS/MS spectrum of its $[M+H]^+$ ion (B).

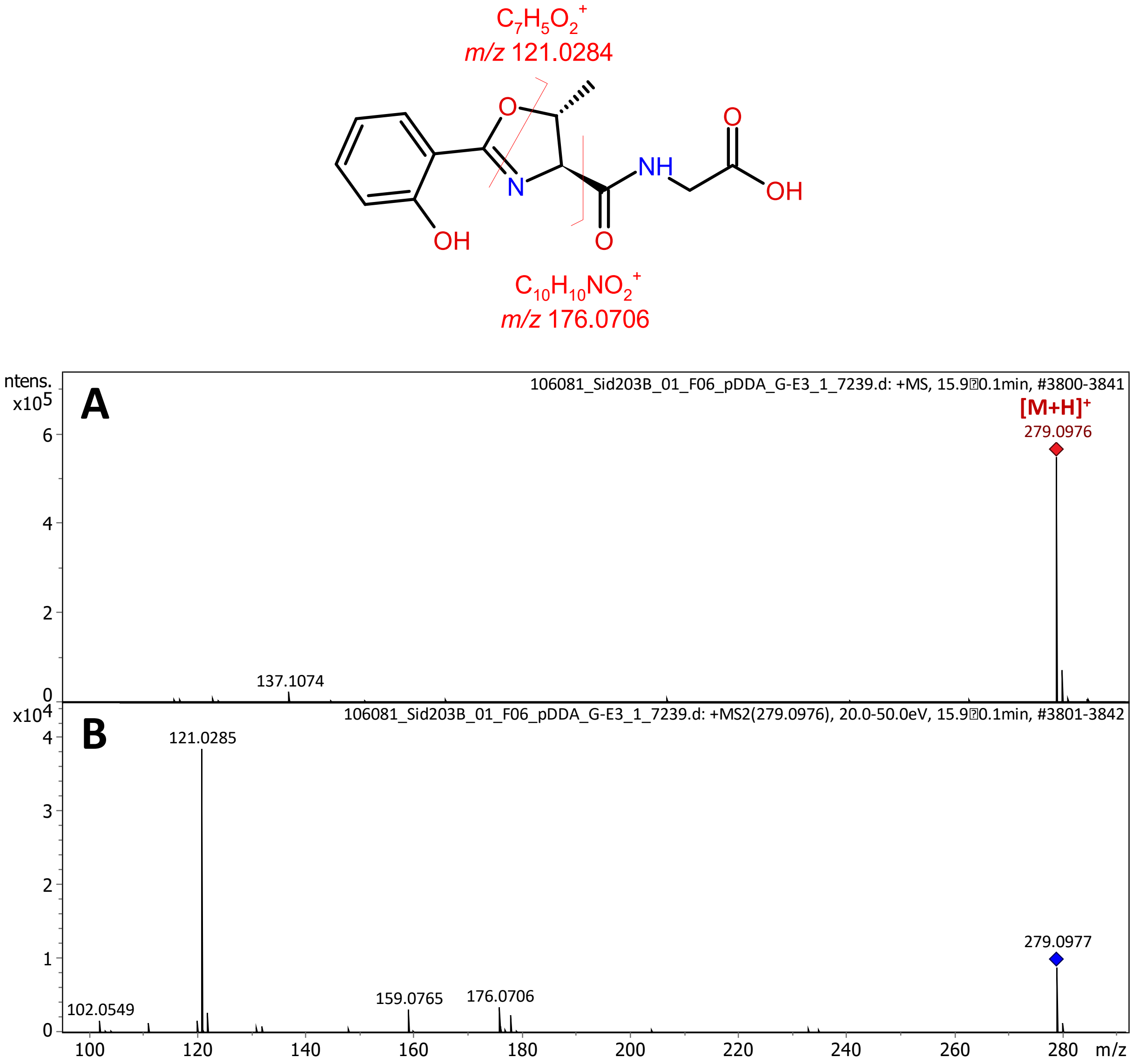


**Supplementary Figure S22.** High resolution ESI-Qq-TOF mass spectrum of kineochelin $E_1$ (A) and high resolution MS/MS spectrum of its $[M+H]^+$ ion (B).

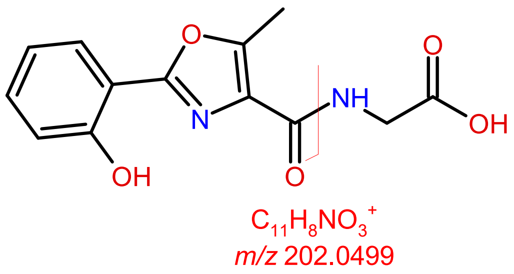


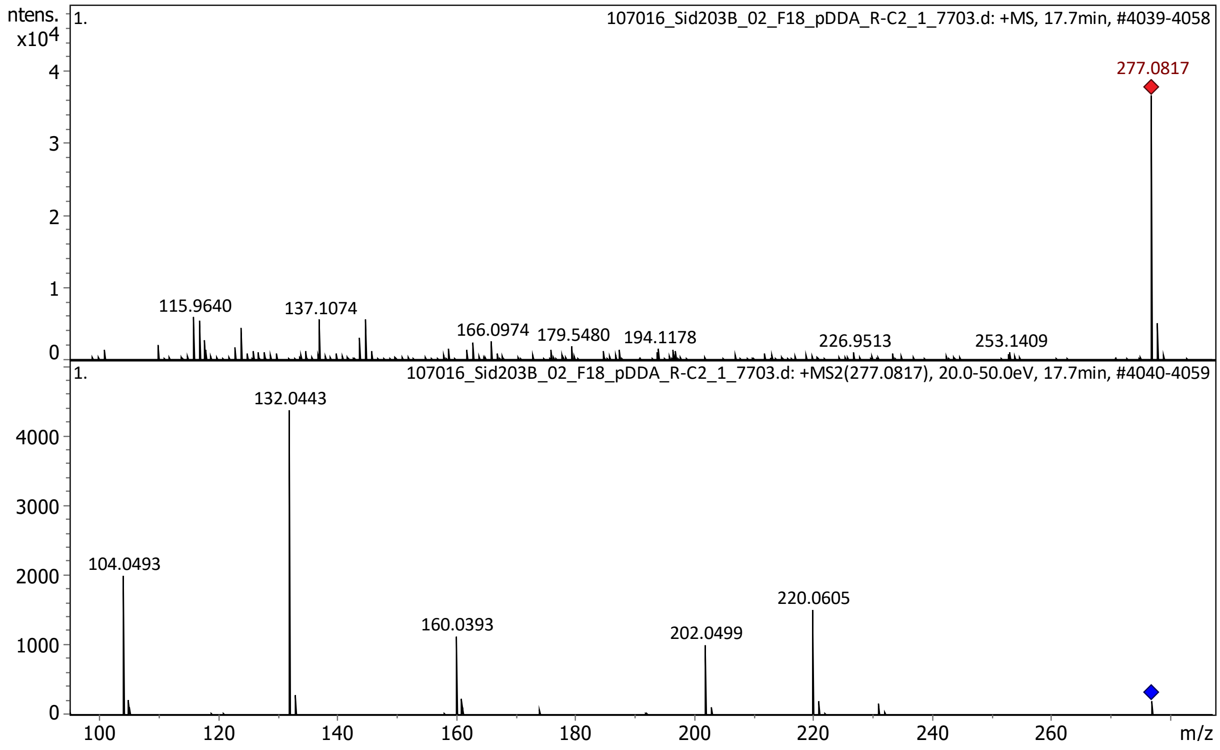


**Supplementary Figure S23.** High resolution ESI-Qq-TOF mass spectrum of kineochelin $E_2$ (A) and high resolution MS/MS spectrum of its $[M+H]^+$ ion (B).

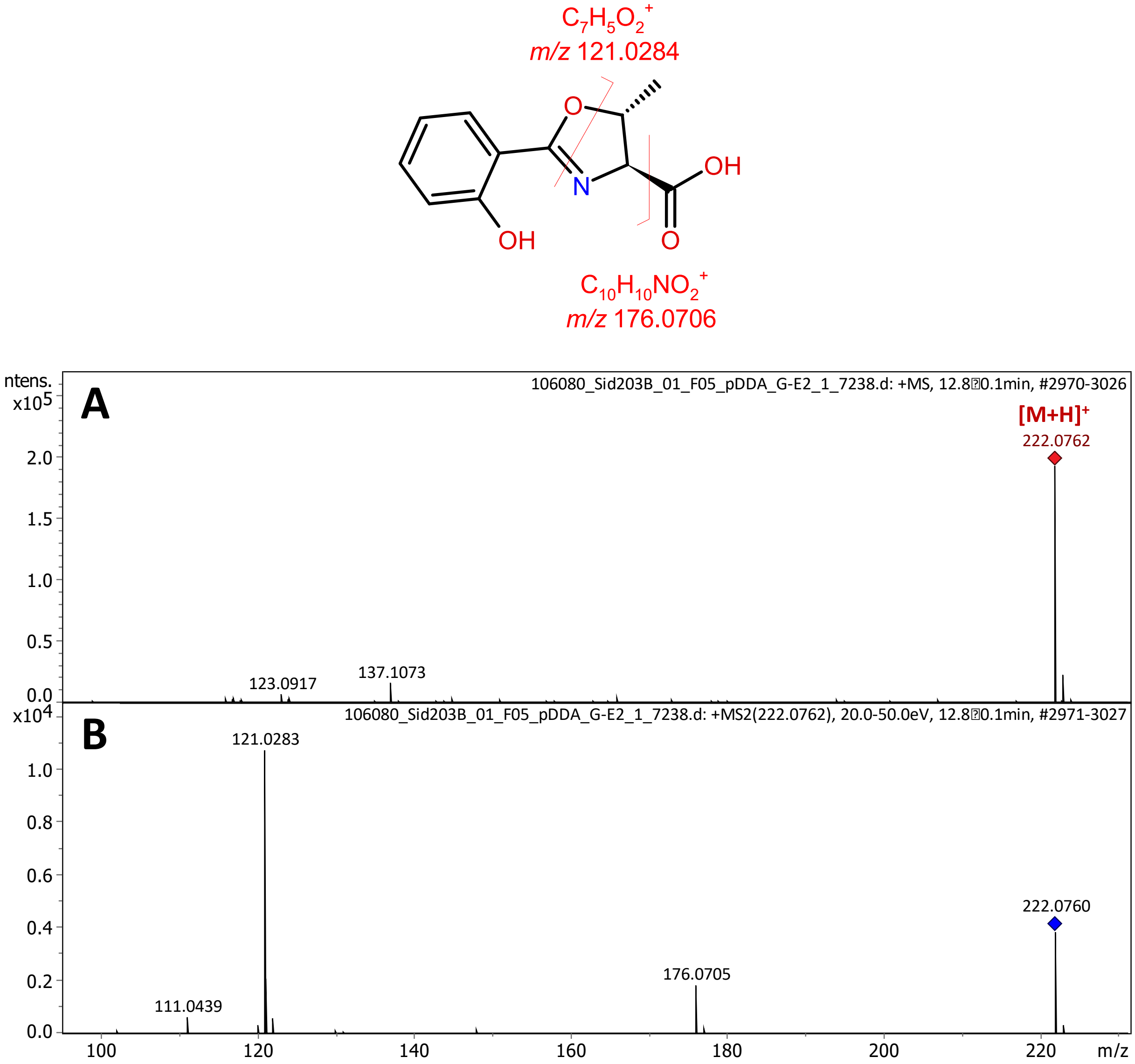


**Supplementary Figure S24.** High resolution ESI-Qq-TOF mass spectrum of vulnibactin 2 (A) and high resolution MS/MS spectrum of its $[M+H]^+$ ion (B).

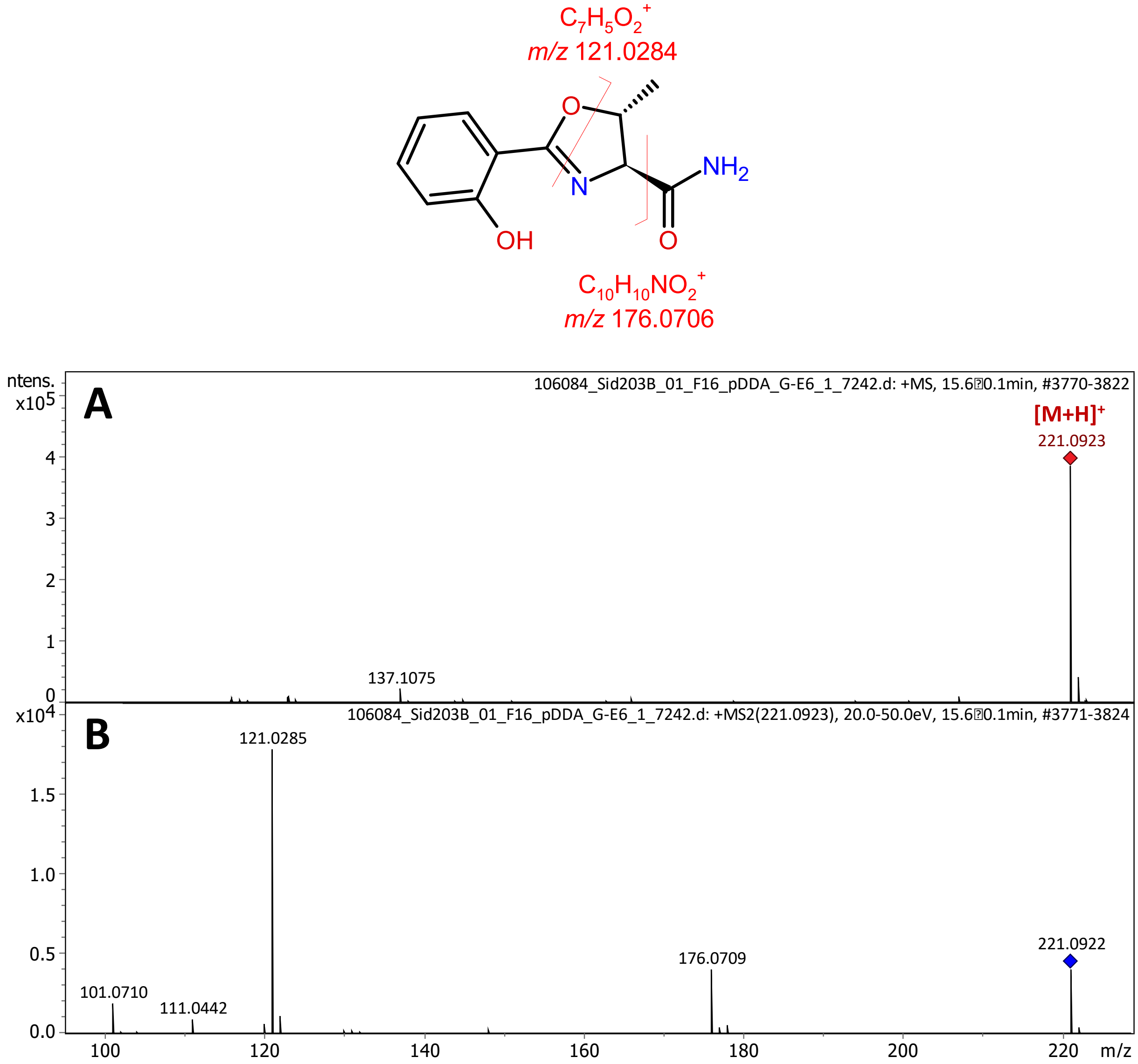


**Supplementary Figure S25.** High resolution ESI-Qq-TOF mass spectrum of pseudomobactin A (A) and high resolution MS/MS spectrum of its $[M+H]^+$ ion (B).

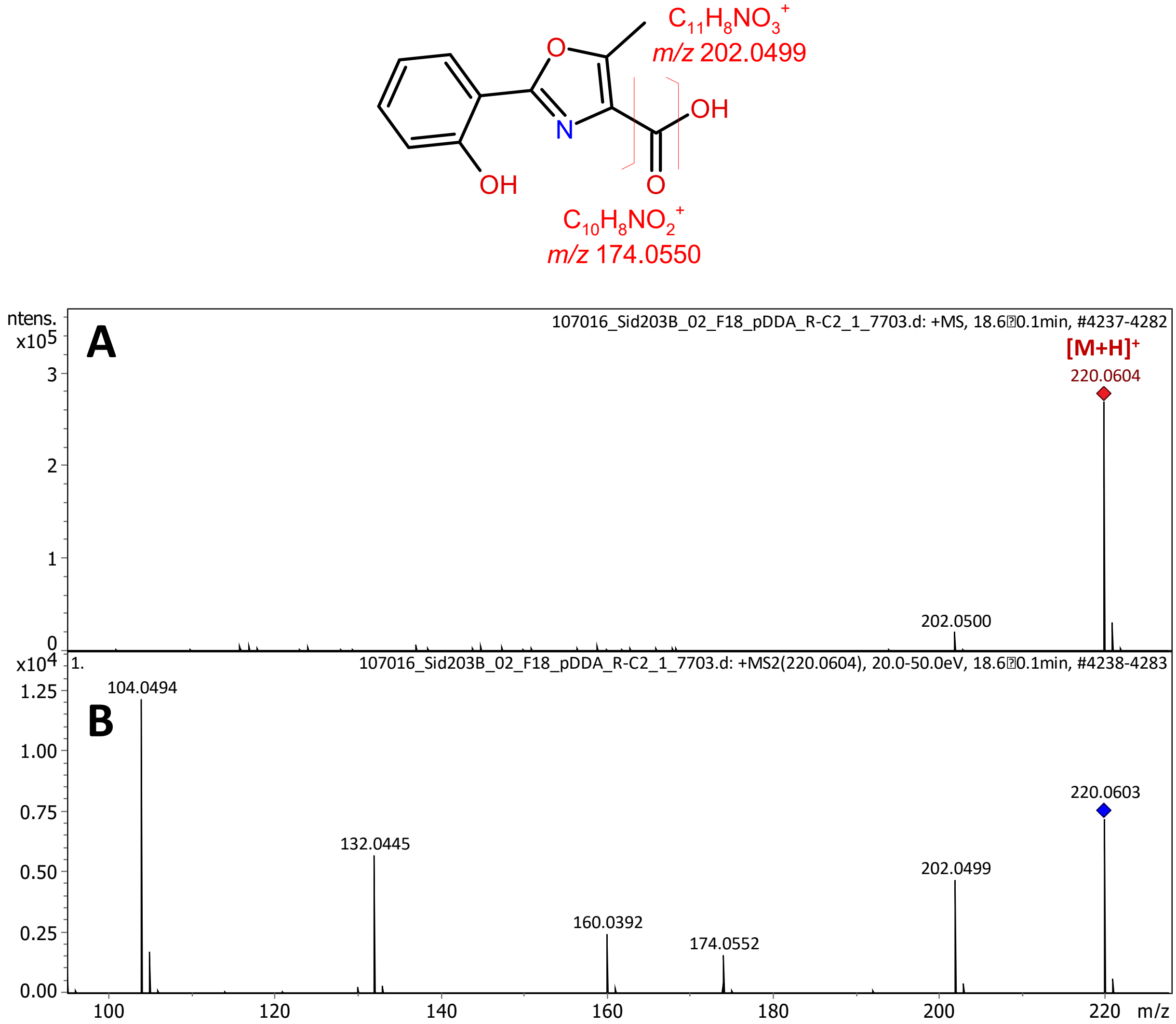


**Supplementary Figure S26.** High resolution ESI-Qq-TOF mass spectrum of asteroidic acid (A) and high resolution MS/MS spectrum of its $[M+H]^+$ ion (B).

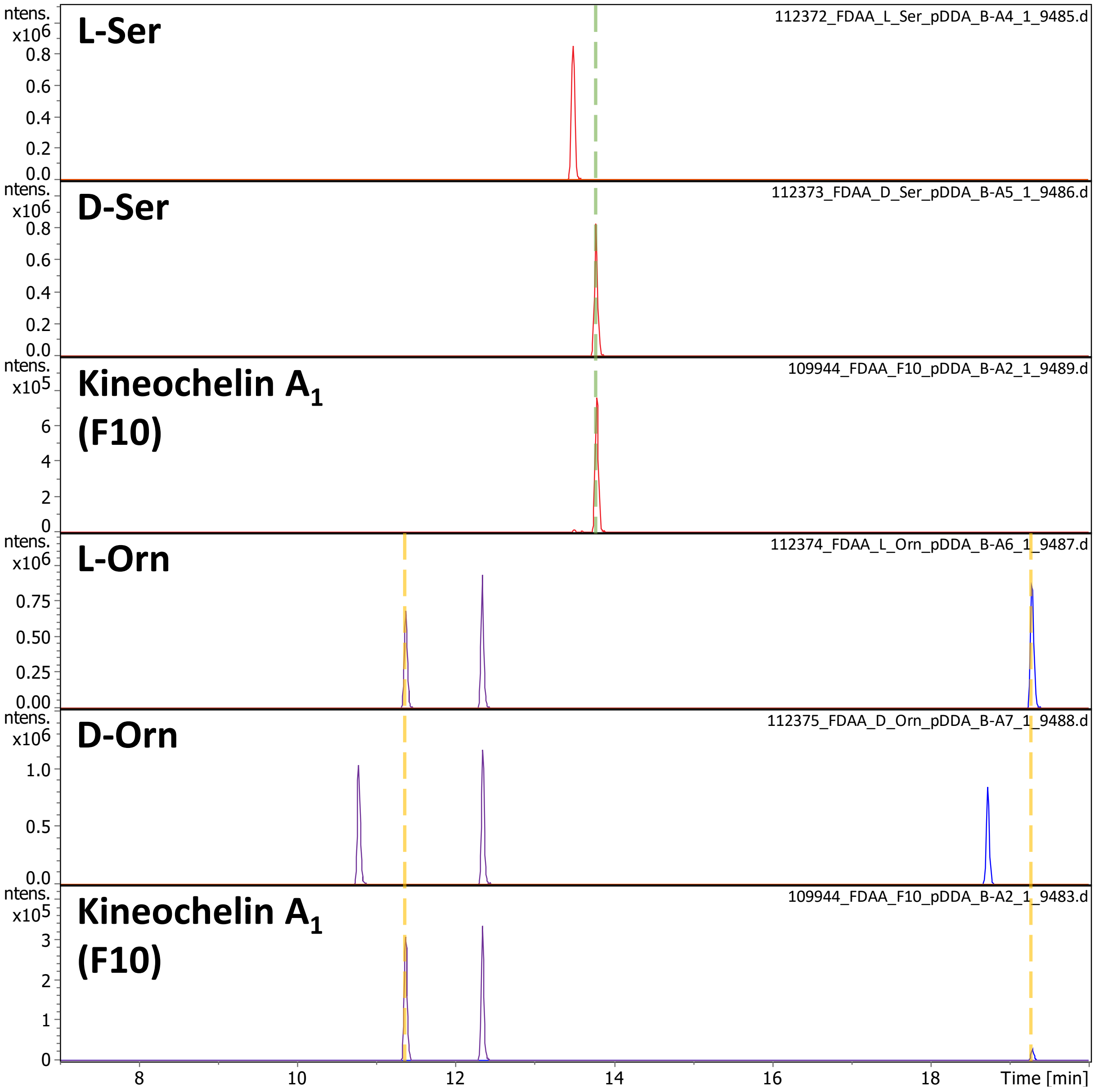


**Supplementary Figure S27. Marfey's analysis of kineochelin $A_1$.** Extracted ion chromatograms (*m/z* 358.0993±0.0050, red; *m/z* 385.1466±0.0050, purple; *m/z* 637.1961±0.0050, blue) showing the signals for L-FDAA-derivatised free amino acids L-Ser, D-Ser, L-Orn, and D-Orn, as well as of D-Ser and L-Orn in the hydrolysed fraction F10, containing mainly kineochelin $A_1$.

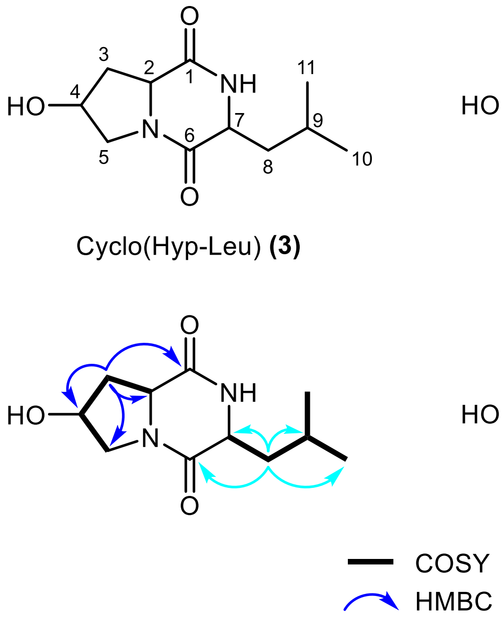


**Figure S28.** Planar structures of cyclo(Hyp-Leu) (**3**) and cyclo(Hyp-Phe) (**4**) with atom numbering and $^1$H-$^1$H COSY and key $^1$H-$^{13}$C HMBC correlations.

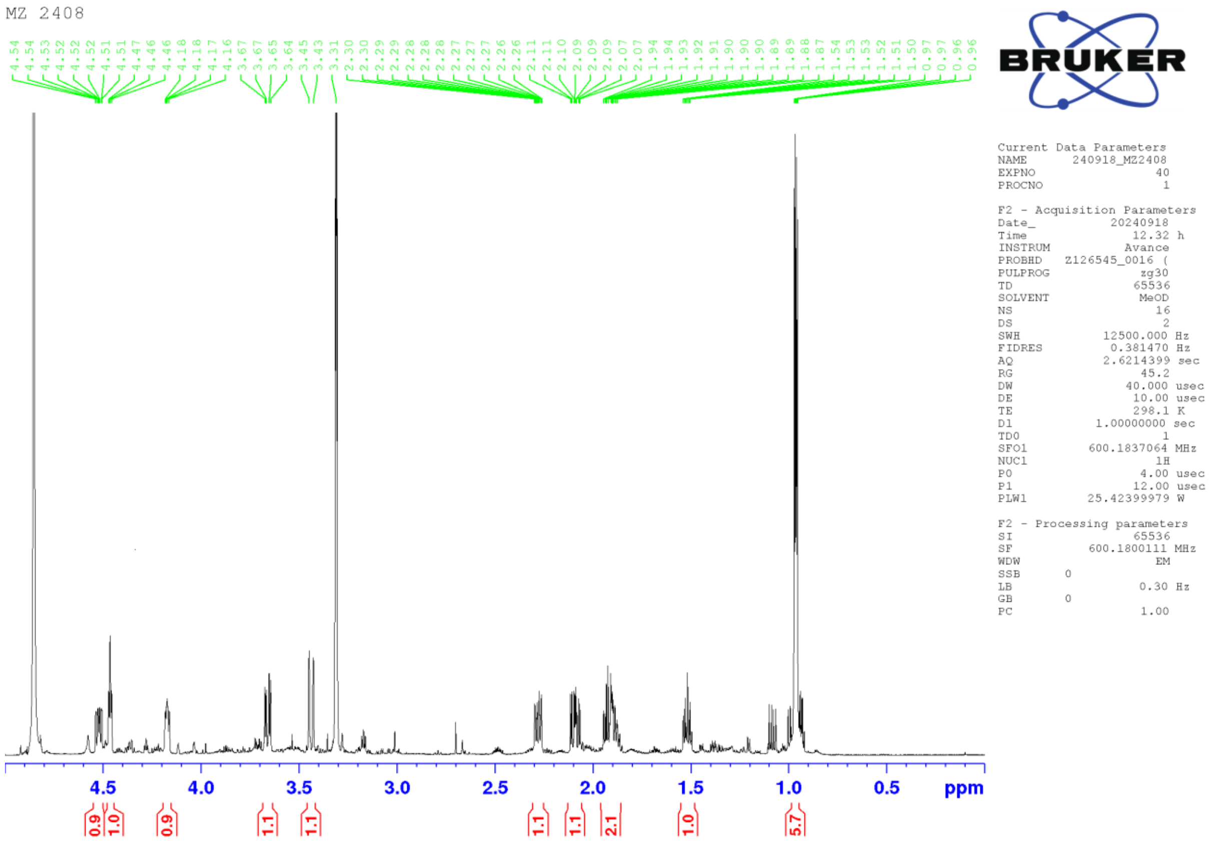


**Figure S28.** $^1$H NMR spectrum of cyclo(Hyp-Leu) (**3**) in $CD_3OH$ at 600 MHz.

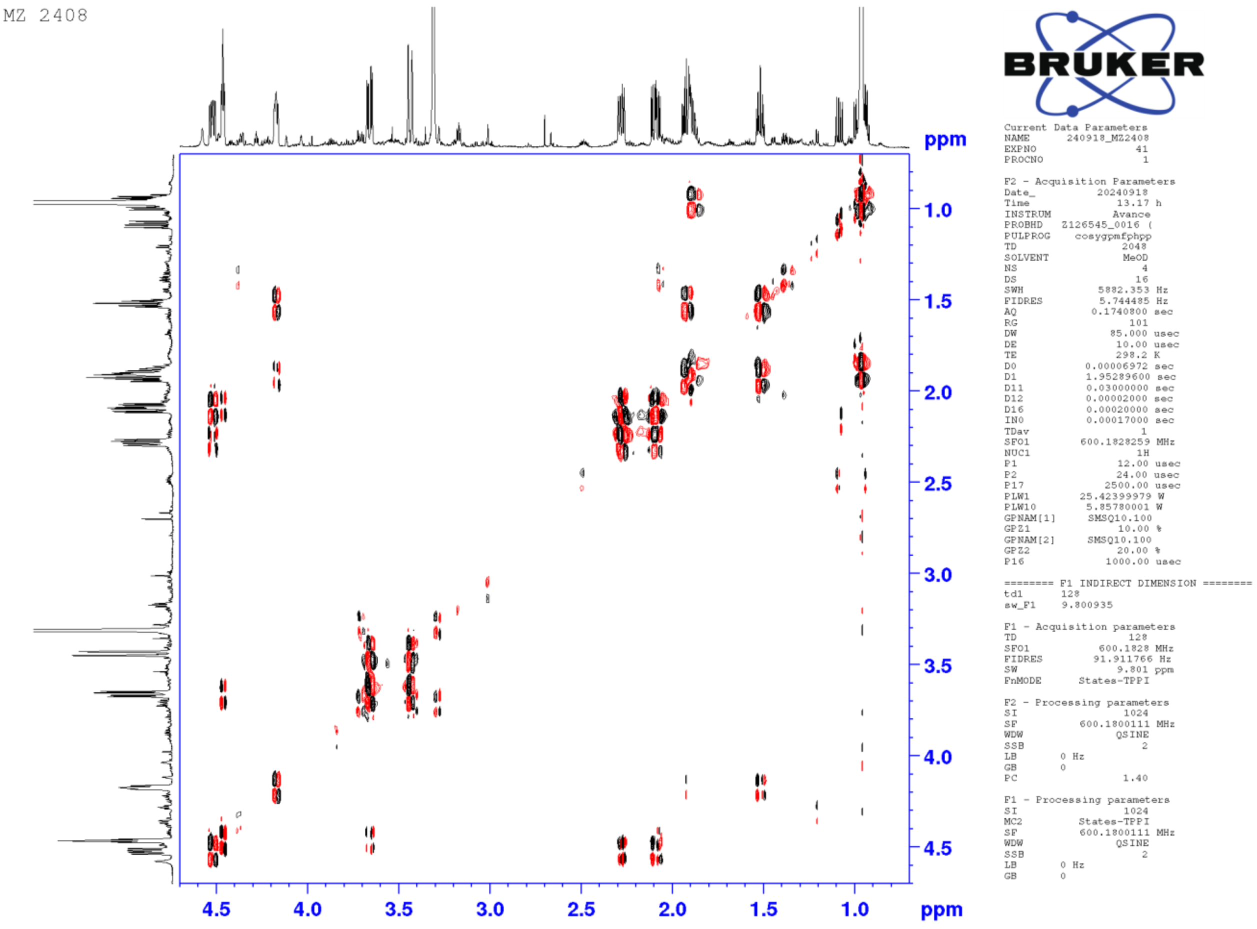


**Figure S30.** COSY spectrum of cyclo(Hyp-Leu) (**3**) in $CD_3OH$ at 600 MHz.

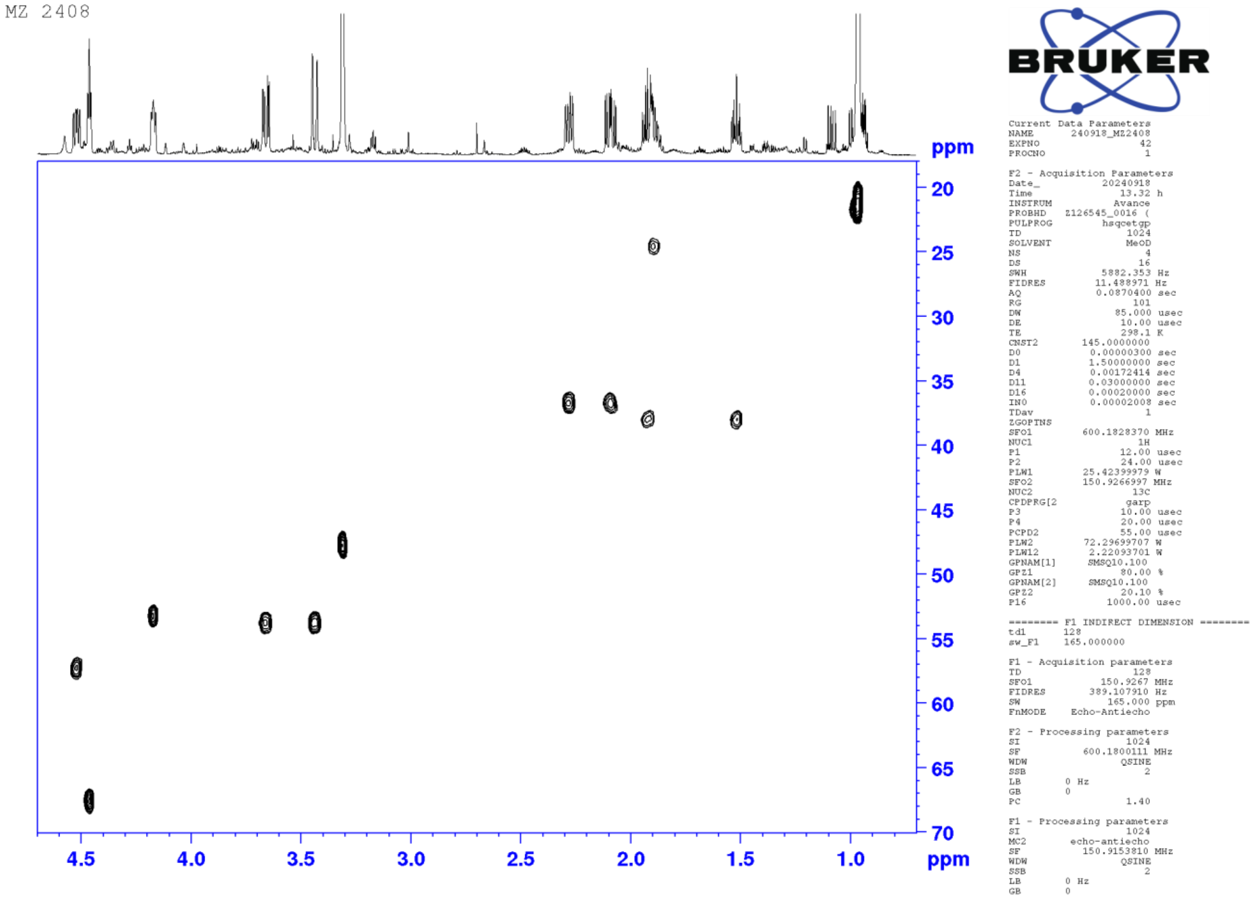


**Figure S31.** HSQC spectrum of cyclo(Hyp-Leu) (**3**) in $CD_3OH$ at 600 MHz.

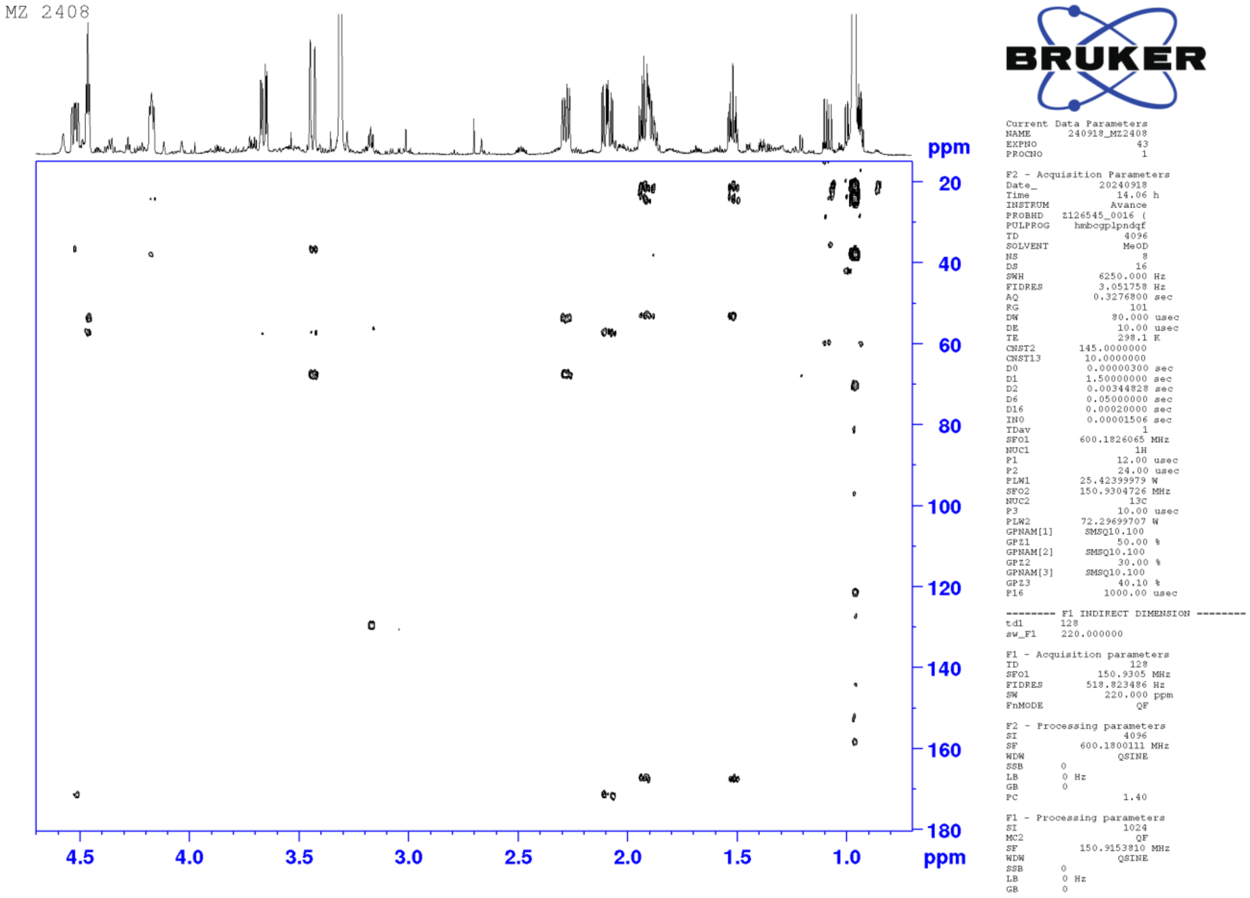


**Figure S32.** HMBC spectrum of cyclo(Hyp-Leu) (**3**) in $CD_3OH$ at 600 MHz.

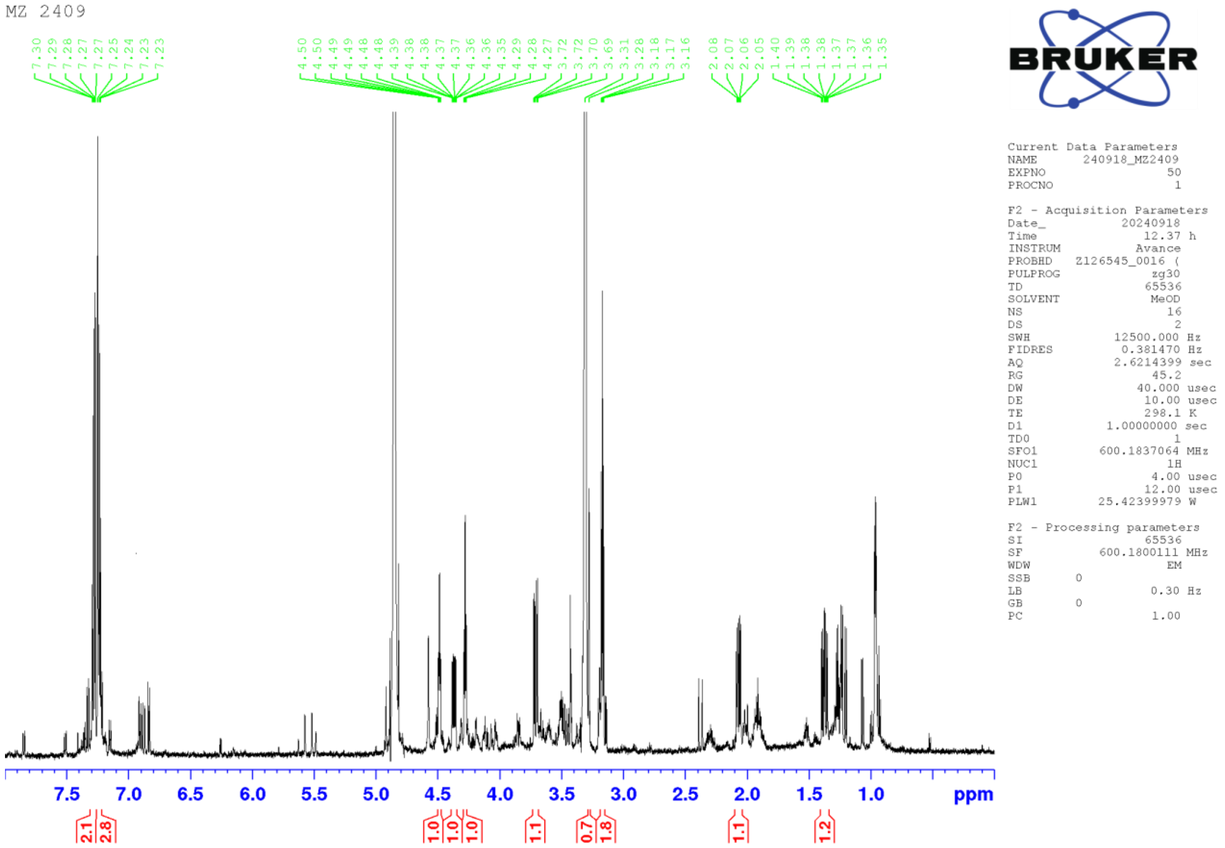


**Figure S33.** $^{1}H$ NMR spectrum of cyclo(Hyp-Phe) (**4**) in $CD_3OH$ at 600 MHz.

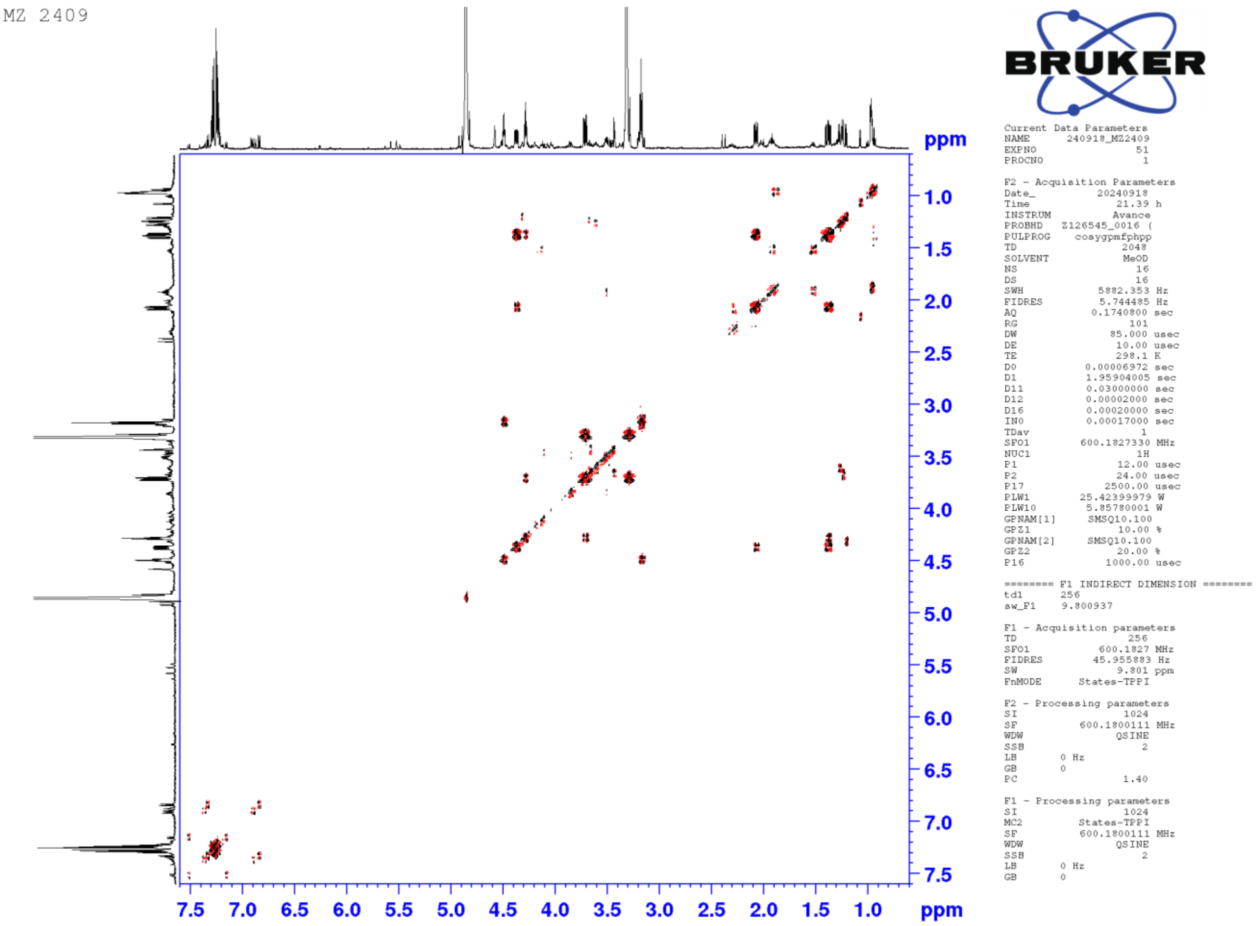


**Figure S34.** COSY spectrum of cyclo(Hyp-Phe) (**4**) in $CD_3OH$ at 600 MHz.

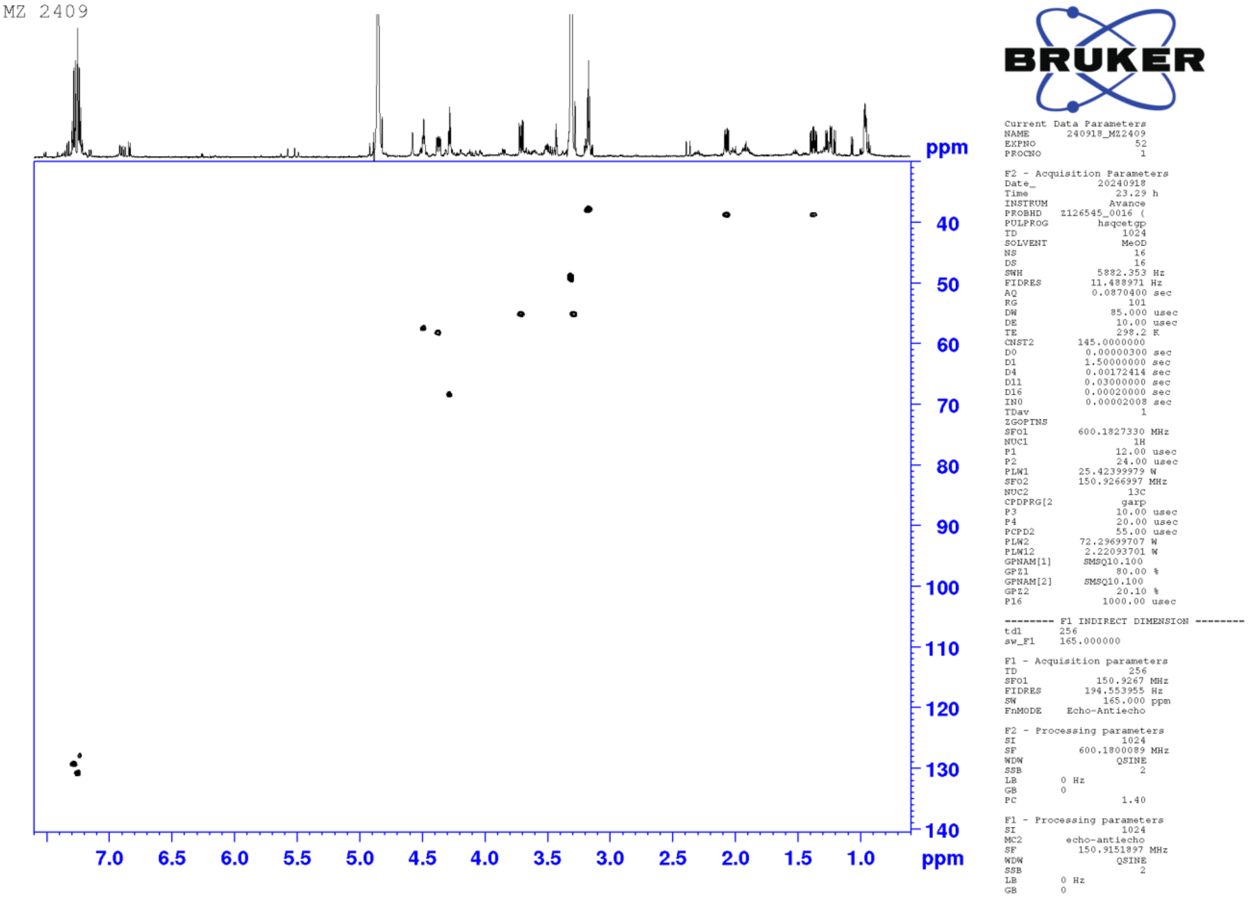


**Figure S35.** HSQC spectrum of cyclo(Hyp-Phe) (**4**) in $CD_3OH$ at 600 MHz.

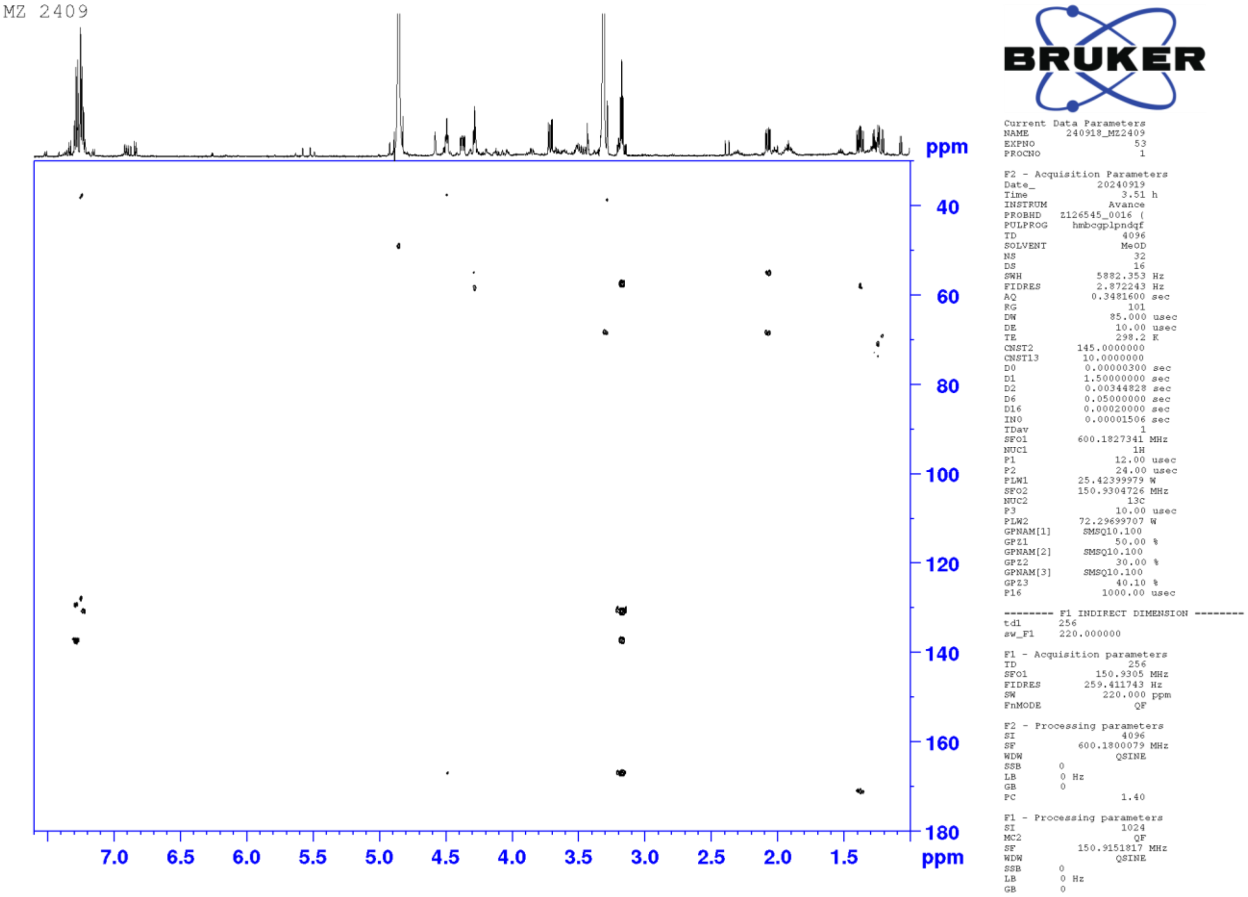


**Figure S36.** HMBC spectrum of cyclo(Hyp-Phe) (**4**) in $CD_3OH$ at 600 MHz.

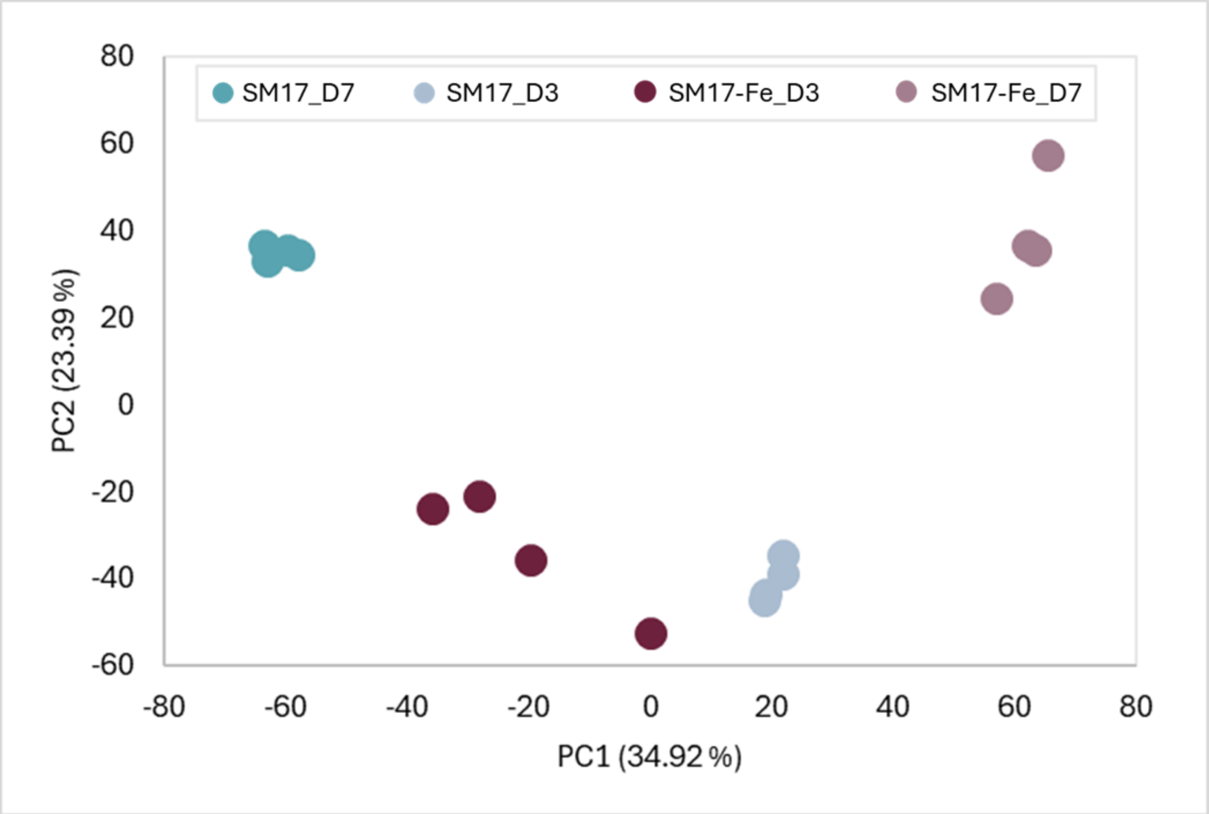


**Supplementary Figure S37. Principal component analysis (PCA) of RNA-seq samples.** PCA of normalized transcript counts from four conditions (SM17 vs. SM17 + 200 μM $FeCl_3$ at day 3 and day 7). Samples separate cleanly (PC1 = 34.9%, PC2 = 21.3%); PC1 distinguishes medium and PC2 captures the time shift. Biological replicates cluster tightly within each group.

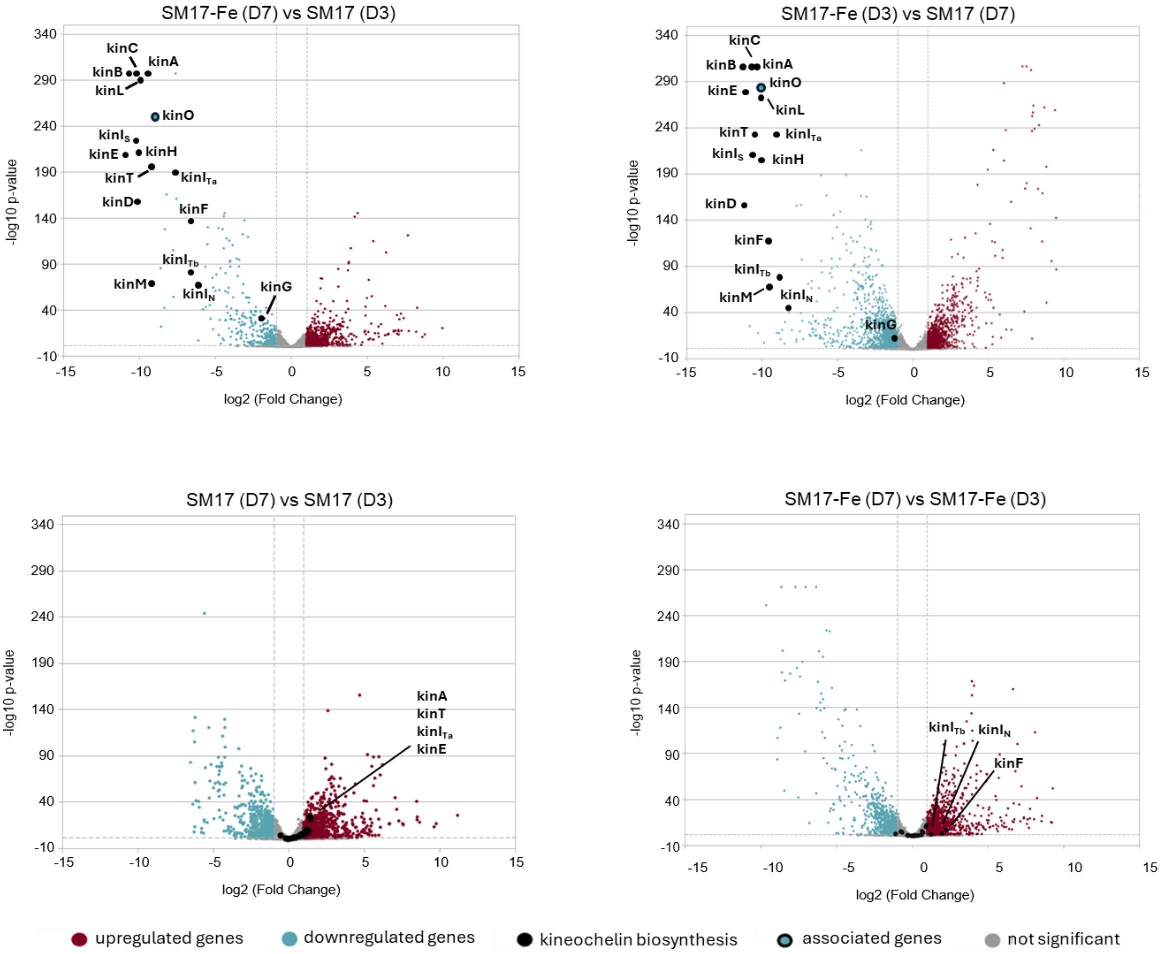


**Supplementary Figure S38. Comparative transcriptomics of *Actinokineospora* sp. UV203 under siderophore-producing and siderophore-depleted growth conditions.** Volcano plots for the indicated pairwise contrasts. Each point is a gene (x-axis: log$_2$ fold change; y-axis: −log$_{10}$(FDR)); dashed lines mark significance thresholds (|log$_2$FC| ≥ 1 and FDR < 0.05). PCA and volcano plots were generated based on normalized counts from DESeq

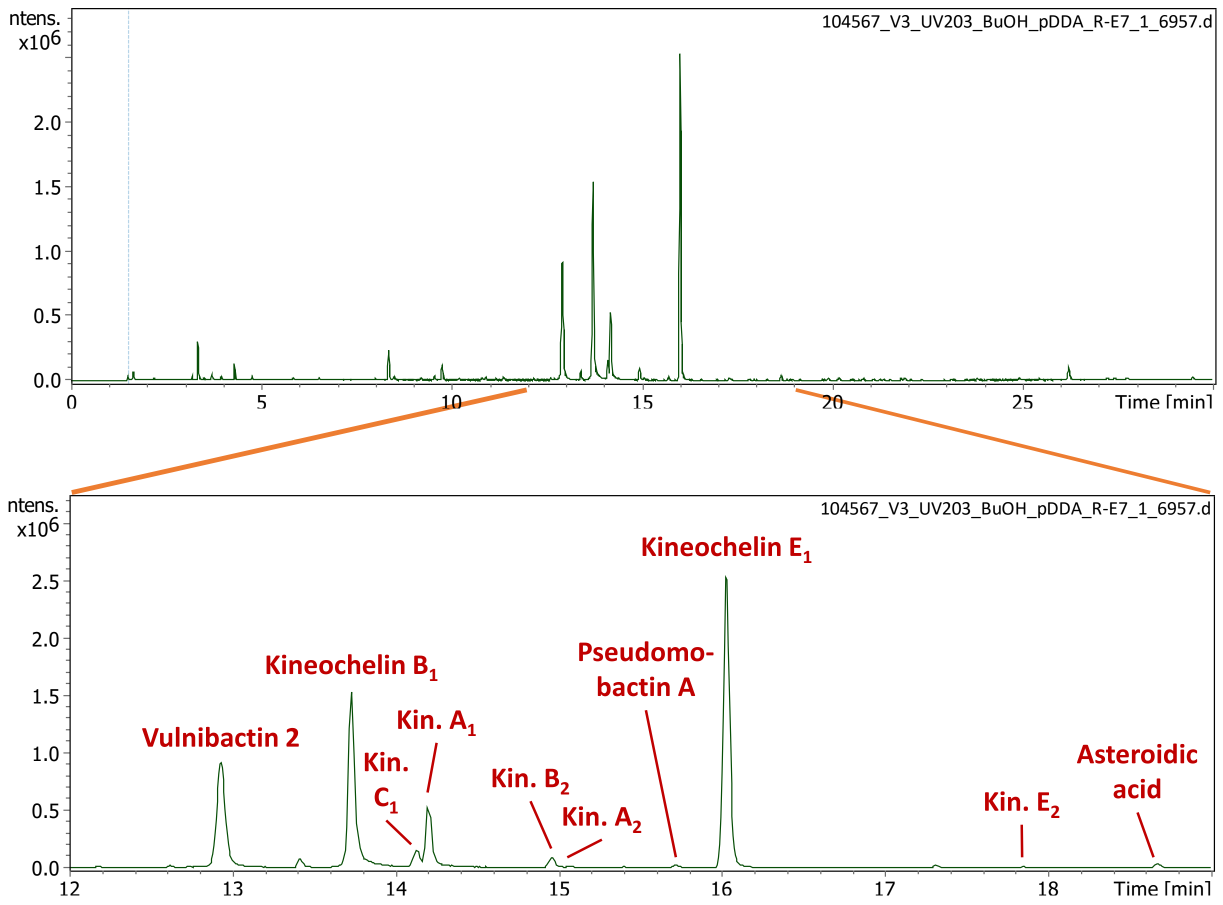


**Supplementary Figure S39.** LC-MS base peak chromatogram of the pre-purified butanol-phase extract showing the kineochelin mixture and co-purified known shunt products.

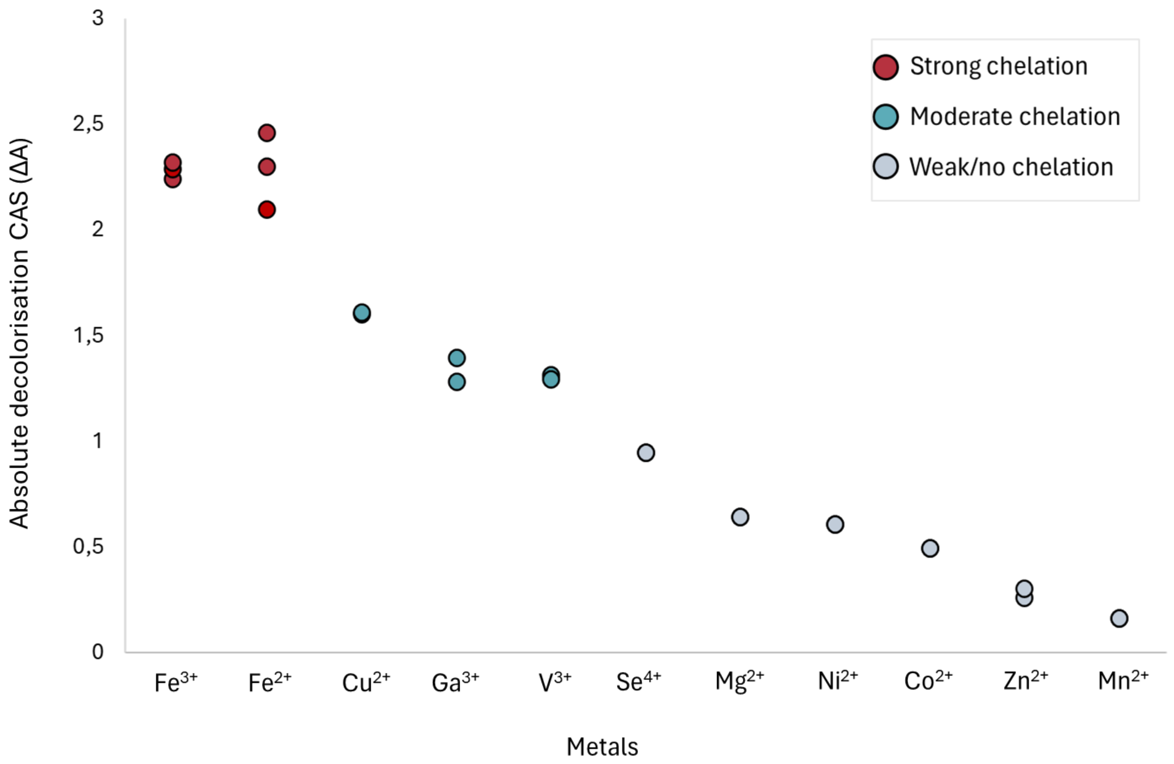


**Supplementary Figure S40. Metal-binding activity of UV203 SM17 culture supernatant measured by Chrome Azurol S (CAS)-shuttle assay.** Absolute CAS decolorization (ΔA) values are shown for individual metal–CAS complexes. Points represent technical replicates (n = 3), and colours indicate relative binding strength normalized to the $Fe^{3+}$ response.

**Supplementary Table S1.** Results from the untargeted LC-MS-based secondary metabolomics analysis of *Actinokineospora* sp. UV203 grown in different media. Groups of secondary metabolites known or presumed to be biosynthetically related are highlighted by the same colour, whereby usually only the most abundant congeners are reported. The listed isoflavone derivatives were only found in cultures from soy-containing media, but not in the media controls. They are thus assumed to be biotransformation products from the soy-isoflavones present in these media.

| # | Rt [min] | *m/z* [M+H]+ | Sum formula (proposed) | *m/z* calcd. | Δ*m/z* [ppm] | Tentative ID | BGC | The Natural Products Atlas | Comment |
|---|---|---|---|---|---|---|---|---|---|
| **1** | 4.0 | 247.1288 | C10H18N2O5 | 247.1288 | 0.0 | Gaburedin D | | NPA028620 | |
| **2** | 4.1 | 279.1009 | C10H18N2O5S | 279.1009 | 0.0 | Gaburedin F | | NPA028622 | |
| **3** | **5.6** | **201.0689** | **C8H12N2O2S** | **201.0692** | **1.6** | **Potentially new natural product** | | | |
| **4** | 6.1 | 261.1446 | C11H20N2O5 | 261.1445 | -0.4 | Gaburedin C | | NPA028619 | |
| **5** | **6.2** | **263.1054** | **C10H18N2O4S** | **263.1060** | **2.1** | **Potentially new natural product** | | | |
| **6** | 6.5 | 261.1445 | C11H20N2O5 | 261.1445 | 0.2 | Gaburedin B | | NPA028618 | |
| **7** | **6.6** | **217.0638** | **C8H12N2O3S** | **217.0641** | **1.6** | **Potentially new natural product** | | | |
| **8** | 7.4 | 295.1288 | C14H18N2O5 | 295.1288 | 0.3 | Gaburedin A | | NPA028617 | |
| **9** | **11.5** | **332.0727** | **C12H17N3O4S2** | **332.0733** | **1.9** | **Potentially new natural product** | | | |
| **10** | **12.6** | **213.0690** | **C9H12N2O2S** | **213.0692** | **1.2** | **Potentially new natural product** | | | |
| **11** | 12.7 | 563.1751 | C27H30O13 | 563.1759 | 1.4 | Genistein-O-deoxyhex-O'-deoxyhex | | | Likely a biotransformation product |
| **12** | 12.8 | 547.1802 | C27H30O12 | 547.1810 | 1.6 | Daidzein-O-deoxyhex-O'-deoxyhex | | | Likely a biotransformation product |
| **13** | 12.8 | 222.0762 | C11H11NO4 | 222.0761 | -0.3 | Vulnibactin 2 | 2.11 | NPA006513 | Shunt product of kineochelins |
| **14** | 12.9 | 693.2379 | C33H40O16 | 693.2389 | 1.5 | Daidzein-O-(deoxyhex-deoxyhex)-O'-deoxyhex | | | Likely a biotransformation product |
| **15** | **13.6** | **611.2307** | **C25H34N6O12** | **611.2307** | **0.0** | **Kineochelin B1** | **2.11** | | **New natural product** |
| **16** | 13.8 | 547.1803 | C27H30O12 | 547.1810 | 1.3 | Daidzein-O-(deoxyhex-deoxyhex) | | | Likely a biotransformation product |
| **17** | **14.0** | **524.1987** | **C22H29N5O10** | **524.1987** | **0.1** | **Kineochelin C1** | **2.11** | | **New natural product** |
| **18** | **14.1** | **723.2944** | **C30H42N8O13** | **723.2944** | **0.0** | **Kineochelin A1** | **2.11** | | **New natural product** |
| **19** | **14.7** | **437.1668** | **C19H24N4O8** | **437.1667** | **-0.2** | **Kineochelin D1** | **2.11** | | **New natural product** |
| **20** | **14.9** | **609.2150** | **C25H32N6O12** | **609.2151** | **0.1** | **Kineochelin B2** | **2.11** | | **New natural product** |
| **21** | **15.0** | **721.2790** | **C30H40N8O13** | **721.2788** | **-0.4** | **Kineochelin A2** | **2.11** | | **New natural product** |
| **22** | 15.6 | 221.0923 | C11H12N2O3 | 221.0921 | -1.1 | Pseudomobactin A | 2.11 | | Shunt product of kineochelins |
| **23** | **15.9** | **279.0976** | **C13H14N2O5** | **279.0975** | **-0.1** | **Kineochelin E1** | **2.11** | | **New natural product** |
| **24** | **17.7** | **277.0817** | **C13H12N2O5** | **277.0819** | **0.5** | **Kineochelin E2** | **2.11** | | **New natural product** |
| **25** | 18.6 | 220.0604 | C11H9NO4 | 220.0604 | 0.1 | Asteroidic acid | 2.11 | | Shunt product of kineochelins |
| **26** | **19.4** | **575.0577** | **C22H24Cl2N4O6S2** | **575.0587** | **1.7** | **Potentially new natural product** | **2.19** | | **Likely NRP containing two Cl-atoms** |
| **27** | **20.2** | **575.0577** | **C22H24Cl2N4O6S2** | **575.0587** | **1.8** | **Potentially new natural product** | **2.19** | | **Likely NRP containing two Cl-atoms** |
| **28** | **23.8** | **276.1589** | **C16H21NO3** | **276.1594** | **1.7** | **Potentially new natural product** | | | |
| **29** | **24.5** | **276.1590** | **C16H21NO3** | **276.1594** | **1.5** | **Potentially new natural product** | | | |

**Supplementary Table S2.** $^{1}H$ (600 MHz) and $^{13}C$ NMR data (151 MHz) of kineochelin $E_1$ (**1**) in $CD_3OD$ in comparison with literature data for pseudomobactin A (*δ* in ppm). A second set of signals (**1'**) was observed in a ratio of approximately 3:5 to the one shown in Table 2. This set of signals was initially hypothesized to belong to a stereoisomer of **1**, but after Marfey's analysis is now assumed to belong to a stable conformer or metal ion complex of kineochelin $E_1$.

| | **Kineochelin $E_1$ (1)** | | **Kineochelin $E_1$ (1')** | | **Pseudomobactin A[a]** | |
|---|---|---|---|---|---|---|
| Position | $\delta_H$ (*J* in Hz) | $\delta_C$, type | $\delta_H$ (*J* in Hz) | $\delta_C$, type | $\delta_H$ (*J* in Hz) | $\delta_C$, type |
| 1 | — | 161.3, C | — | 163.0, C | — | 161.1, C |
| 2 | 6.96, m, ov | 117.9, CH | 6.89-6.97, m, ov | 118.5, CH | 6.96, d (8.3) | 117.7, CH |
| 3 | 7.41, m | 135.2, CH | 7.51, m | 137.4, CH | 7.41, td (8.3, 1.7) | 135.0, CH |
| 4 | 6.90, m, ov | 120.1, CH | 6.89-6.97, m, ov | 120.6, CH | 6.90, td (7.4, 1.7) | 119.9, CH |
| 5 | 7.69, dd (7.9, 1.7) | 129.7, CH | 7.96, dd (8.0, 1.6) | 131.7, CH | 7.68, dd (7.4, 1.7) | 129.5, CH |
| 6 | — | 111.8, C | — | 113.6, C | — | 111.6, C |
| 7 | — | 168.3, C | — | 170.4, C | — | 167.8, C |
| 8 | 1.57, d (6.3) | 21.6, $CH_3$ | 1.50, d (6.5) | 17.1, CH3 | 1.57, d (6.3) | 21.4, $CH_3$ |
| 9 | 4.91, dq (7.5, 6.3) | 80.8, CH | 5.54, qd (6.3, 6.3) | 72.3, CH | 4.90, qd (7.3, 6.3) | 80.6, CH |
| 10 | 4.51, d (7.5) | 75.9, CH | 3.98, d (6.0), ov | 58.7, CH | 4.46, d (7.3) | 75.5, CH |
| 11 | — | 173.3, C | — | 169.2, C | — | 175.6, C |
| 12a | 3.93, d (17.0), ov | 43.0, $CH_2$ | 3.96, d (17.3), ov | 43.8, CH2 | — | — |
| 12b | 3.87, d (17.5) | | 3.73, d (17.3) | | — | — |
| 13 | — | 173.8[b], C | — | 174.9[b], C | — | — |

[a] Values obtained from Oluwabusola *et al.* (2021)[1]

[b] Value determined from HMBC spectrum

**Supplementary Table S3.** $^{1}H$ (600 MHz, $CD_3OD$) data of cyclo(Hyp-Leu) (**3**) and cyclo(Hyp-Phe) (**4**) in comparison with literature data ($\delta$ in ppm).

| | **cyclo(Hyp-Leu) (3)** | **cyclo(*L*-Hyp-*L*-Leu)**[a] | **cyclo(Hyp-Phe) (4)** | **cyclo(*L*-Hyp-*L*-Phe)**[a] |
|---|---|---|---|---|
| Position | $\delta_H$ (*J* in Hz) | $\delta_H$ (*J* in Hz) | $\delta_H$ (*J* in Hz) | $\delta_H$ (*J* in Hz) |
| 1 | — | — | — | — |
| 2 | 4.52, ddd (11.1, 6.5, 1.3) | 4.55, dd (6.6, 1.7) | 4.37, ddd (11.7, 5.9, 1.7) | 4.39, dq (6.0, 1.8) |
| 3 | 2.28, ddt (13.2, 6.5, 1.3)<br>2.09, ddd (13.2, 11.1, 4.3) | 2.30, dd (13.2, 6.5)<br>2.12, m | 2.07, dd (13.0, 6.0), br<br>1.37, ddd (12.9, 11.8, 4.6) | 2.09, dd (13.0, 5.9)<br>1.41, ddd (20.8, 12.0, 0.6) |
| 4 | 4.46, dd (4.3), br | 4.48, t (4.4) | 4.28, dd (4.8), br | 4.29, t (4.8) |
| 5 | 3.66, dd (12.8, 4.5)<br>3.44, d (13.0), br | 3.68, dd (12.8, 4.4)<br>3.45, d (12.0) | 3.71, dd (13.0, 5.1)<br>3.29, m, ov | 3.72, dd (13.0, 5.1)<br>3.34, m |
| 6 | — | — | — | — |
| 7 | 4.17, m | 4.19, m | 4.49, m | 4.50, dt, (5.2, 1.8) |
| 8 | 1.92, m, ov<br>1.52, m | 1.92, m<br>1.55, m | 3.17, dd, br | 3.18, m |
| 9 | 1.89, m, ov | 1.90, m | — | — |
| 10 | 0.97, d (6.4) | 0.97, d (6.4) | 7.28, m | 7.27, m |
| 11 | 0.96, d (6.4) | 0.96, d (6.4) | 7.24, m, ov | 7.30, m |
| 12 | — | — | 7.22-7.25, m, ov | 7.23, m |

[a] Values obtained from Xiang *et al.*(2020)

**Supplementary Table S4.** Local gene identities between *kin* cluster and related amychelin, cahuitamycin, and gobichelin clusters.

| CDS Name | Identifier in BGC | BlastP sequence identity to amychelin (BGC0000300.5) | Identity / Coverage (%) | BlastP sequence identity to cahuitamycin (BGC0001351.) | Identity / Coverage (%) | BlastP sequence identity to gobichelin BGC | Identity / Coverage (%) |
|---|---|---|---|---|---|---|---|
| **kinG** | ctg2_2469 | - | - | - | - | - | - |
| **kinX$_2$** | ctg2_2470 | - | - | - | - | - | - |
| **kinL** | ctg2_2471 | amcL (SSMG_02545) salicylate_synthase | 49.41 / 91.40 | cahI (AMK48233.1) salicylate_synthase | 49.55 / 93.12 | - | - |
| **kinH** | ctg2_2472 | amcH (SSMG_02542) 2,3-dihydroxybenzoate-AMP_ligase | 58.46 / 95.76 | cahJ (AMK48234.1) salicylate-AMP_ligase | 63.16 / 97.24 | gobK (AGE11892.1) 2,3-dihydroxybenzoate-AMP_ligase | 58.59 / 96.49 |
| **kinI$_S$** | ctg2_2473 | amcM (SSMG_02534) iron compound ABC transporter | 27.64 / 75.44 | - | - | gobM (AGE11894.1) iron_siderophore_uptake_ABC_system | 53.43 / 97.08 |
| **kinM** | ctg2_2474 | amcA (SSMG_02541) major facilitator superfamily transporter multidrug resistance protein | 46.60 / 86.99 | cahH (AMK48232.1) siderophore_export_protein | 46.39 / 94.75 | - | - |
| **kinB** | ctg2_2475 | amcF (SMG_02536) NRPS | 44.47 / 71.71 | cahB (AMK48226.1) NRPS | 44.39 / 71.63 | gobR (AGE11898.1) NRPS | 46.39 / 70.97 |
| **kinC** | ctg2_2476 | amcE (SSMG_02537) NRPS | 44.93 / 57.42 | cahD (AMK48228.1) NRPS | 45.40 / 40.34 | gobR (AGE11898.1) NRPS | 38.64 / 58.46 |
| **kinD** | ctg2_2477 | amcC (SSMG_02539) mbtH_protein | 65.67 / 97.10 | cahE (AMK48229.1) MbtH-like_protein | 68.66 / 97.10 | gobL (AGE11893.1) MbtH | 59.38 / 92.75 |
| **kinE** | ctg2_2478 | - | - | | | - | - |
| **kinA** | ctg2_2479 | amcG (SSMG_02535) NRPS | 41.21 / 99.07 | cahA (AMK48225.1) NRPS | 41.00/ 98.23 | gobJ (AGE11891.1) NRPS | 41.52 / 99.25 |
| **kinT** | ctg2_2480 | amcG (SSMG_02535) NRPS | 39.06 / 19.16 | cahA (AMK48225.1) NRPS | 43.06 / 21.56 | gobJ (AGE11891.1) NRPS | 46.56 / 21.86 |
| **kinI$_{Ta}$** | ctg2_2481 | amcI (SSMG_02543) transport system permease protein | 54.55 / 89.80 | cahT2 (AMK48235.1) iron_ABC_transporter_permease | 49.56 / 98.83 | gobP (AGE11897.1) iron_siderophore_transporter | 57.79 / 89.80 |
| **kinI$_{Tb}$** | ctg2_2482 | amcJ (SSMG_02544) transport system permease protein | 42.42 / 91.22 | cahT3 (AMK48236.1) iron_ABC_transporter_permease | 47.18 / 84.99 | gobO (AGE11896.1) ABC_transporter | 58.91 / 98.02 |
| **kinI$_N$** | ctg2_2483 | - | - | cahT4 (AMK48243.1) putative_ABC_transporter_ATPase | 41.11 / 31.90 | gobN (AGE11895.1) ABC_transporter | 64.73 / 92.47 |
| **kinF** | ctg2_2484 | - | - | - | - | - | - |
| **kinO*** | ctg2_1302 | amcP (WP_009078944.1) lysine_N(6)-hydroxylase/L-ornithine_N(5)-oxygenase_family_protein | 67.06 / 94.78 | -[¥] | -[¥] | -[¥] | -[¥] |

(*) gene located outside of *kin* cluster; (-) no match in cluster identified; (-[¥]) no match outside of BGC identified due to genome unavailability

**Supplementary Table S5.** Closest NCBI Blast and AntiSMASH database matches to the core biosynthetic enzymes KinA, KinB, and KinC in the *kin* cluster.

| CDS Name | Identifier in BGC | Hit Record | Organisms | Amino acid Identity [%] | Database |
|---|---|---|---|---|---|
| **kinB** | ctg2_2475 | NZ_LT629701 | *Streptomyces atratus* SCSIO_ZH16 | 47.48 | AntiSMASH v4[a] |
| | | NZ_CP027306 | *Alllokutzneria albata* DSM 44149 | 47.18 | AntiSMASH v4 |
| | | NZ_WOFH01000003 | *Actinomadura litoris* NEAU-AAG5 | 47.78 | AntiSMASH v4 |
| | | WP_091382965.1 | *Actinokineospora alba* | 92.67 | RefSeq non-redundant proteinsb[2] |
| | | WP_187220517.1 | *Actinokineospora xionganensis* | 92.11 | RefSeq non-redundant proteins |
| | | WP_431423574.1 | *Actinokineospora* sp. | 91.51 | RefSeq non-redundant proteins |
| **kinC** | ctg2_2476 | NZ_JANCLX010000004 | *Streptomyces cucumeris* | 40.31 | AntiSMASH v4 |
| | | NZ_WPBY01000002 | *Streptomyces* sp. | 39.99 | AntiSMASH v4 |
| | | NZ_CP114036 | *Streptomyces* sp. | 39.36 | AntiSMASH v4 |
| | | WP_431423573.1 | *Actinokineospora* sp. | 90.73 | RefSeq non-redundant proteins |
| | | WP_187220518.1 | *Actinokineospora xionganensis* | 89.56 | RefSeq non-redundant proteins |
| | | WP_091382965.1 | *Actinokineospora alba* | 60.80 | RefSeq non-redundant proteins |
| **kinA** | ctg2_2479 | NZQHCP01000002 | *Actinokineospora spheciospongiae* CECT 8578 | 54.87 | AntiSMASH v4 |
| | | NZ_CP09251 | *Streptomyces* sp. TYQ1024 | 54.07 | AntiSMASH v4 |
| | | NZ_JIAI01000001 | *Pseudonocardia acaciae* DSM 45401 | 53.64 | AntiSMASH v4 |
| | | WP_166658022.1 | *Actinokineospora alba* | 94.04 | RefSeq non-redundant proteins |
| | | WP_431423569.1 | *Actinokineospora* sp. | 93.01 | RefSeq non-redundant proteins |
| | | WP_187220520.1 | *Actinokineospora xionganensis* | 91.61 | RefSeq non-redundant proteins |

[a] Blin et al., 2024[3], [b] Pruitt et al., 2025[2]

**Supplementary Table S6.** Closest homologues of coding sequences (CDSs) within the *kin* biosynthetic gene cluster of *Actinokineospora* sp. UV203.

| CDS Name | Identifier in BGC | Length [AA] | Closest homologue | Organism of rigin | Identity [%] | Alignment length [AA] | Homologue accession number |
|---|---|---|---|---|---|---|---|
| **kinG** | ctg2_2469 | 511 | amidohydrolase | *Actinokineospora alba* | 92.76 | 516 | WP_228770232.1 |
| **kinX$_2$** | ctg2_2470 | 111 | hypothetical protein | *Actinokineospora xionganensis* | 87.50 | 88 | WP_187220511.1 |
| **kinL** | ctg2_2471 | 465 | salicylate synthetase | *Actinokineospora alba* | 97.57 | 465 | WP_228770233.1 |
| **kinH** | ctg2_2472 | 542 | (2,3-dihydroxybenzoyl)adenylate synthase | *Actinokineospora alba* | 95.94 | 542 | WP_091382958.1 |
| **kinI$_S$** | ctg2_2473 | 342 | ABC transporter substrate-binding protein | *Actinokineospora xionganensis* | 94.15 | 342 | WP_187220515.1 |
| **kinM** | ctg2_2474 | 438 | enterobactin transporter EntS | *Actinokineospora* sp. | 94.08 | 437 | WP_431423575.1 |
| **kinB** | ctg2_2475 | 3627 | non-ribosomal peptide synthetase | *Actinokineospora alba* | 92.67 | 4630 | WP_091382965.1 |
| **kinC** | ctg2_2476 | 2506 | amino acid adenylation domain-containing protein | *Actinokineospora* sp. | 90.73 | 2507 | WP_431423573.1 |
| **kinD** | ctg2_2477 | 69 | MbtH family protein | *Actinokineospora* sp. | 98.55 | 69 | WP_091382967.1 |
| **kinE** | ctg2_2478 | 402 | SagB family peptide dehydrogenase | *Actinokineospora xionganensis* | 95.52 | 402 | WP_187220519.1 |
| **kinA** | ctg2_2479 | 1072 | non-ribosomal peptide synthetase | *Actinokineospora alba* | 94.04 | 1073 | WP_166658022.1 |
| **kinT** | ctg2_2480 | 334 | alpha/beta fold hydrolase | *Actinokineospora alba* | 92.22 | 334 | WP_133794513.1 |
| **kinI$_{Ta}$** | ctg2_2481 | 343 | FecCD family ABC transporter permease | *Actinokineospora xionganensis* | 95.92 | 343 | WP_312880281.1 |
| **kinI$_{Tb}$** | ctg2_2482 | 353 | FecCD family ABC transporter permease | *Actinokineospora alba* | 97.73 | 353 | WP_091382973.1 |
| **kinI$_N$** | ctg2_2483 | 279 | ABC transporter ATP-binding protein | *Actinokineospora* sp. | 96.77 | 279 | WP_431423564.1 |
| **kinF** | ctg2_2484 | 60 | (2Fe-2S)-binding protein | *Actinokineospora* sp. | 91.67 | 60 | WP_431423563.1 |
| **kinO*** | ctg2_1302 | 441 | lysine N(6)-hydroxylase/L-ornithine N(5)-oxygenase family protein | *Actinokineospora alba* | 97.51 | 441 | WP_091382199.1 |

(*) gene located outside of *kin* cluster

**Supplementary Table S7.** Microbial strains used for preliminary antimicrobial susceptibility testing of the kineochelins.

| Type | Classification | Taxonomy | Strain ID | Origin | Medium used | Inhibition |
|---|---|---|---|---|---|---|
| G+ bacteria | Bacillota | *Brevibacillus sp.* | UV260 | environmental, Antarctic | R2A | - |
| | | *Bacillus subtilis* | P12204 | environmental, Antarctic | MHA | - |
| | | *Paenibacillus* sp. | P12302 | environmental, Antarctic | R2A | yes |
| | | *Paenibacillus* sp. | UV231 | environmental, Antarctic | R2A | yes |
| | | *Paenisporosarcina* sp. | P12233 | environmental, Antarctic | MHA | - |
| | | *Staphylococcus aureus* | 708 | clinical | MHA | - |
| | | *Enterococcus faecalis* | 328 | clinical | MHA | - |
| | | *Streptococcus pyogenes* | 556 | clinical | MHA | yes |
| | Actinomycetota | *Nocardioides* sp. | UV208 | environmental, Antarctic | GPHF | - |
| | | *Rhodococcus* sp. | UV292 | environmental, Antarctic | MHA | - |
| | | *Gordonia sp.* | P12182 | environmental, Antarctic | MHA | yes |
| | | *Kytococcus sedentarius* | UV219 | environmental, Antarctic | MHA | yes |
| | | *Cryobacterium* sp. | UV168 | environmental, Antarctic | R2A | yes |
| | | *Micrococcus luteus* | CCM 169$^{T}$ | environmental | MHA | yes |
| | | *Arthrobacter* sp. | P12200 | environmental, Antarctic | MHA | - |
| | | *Pseudoarthrobacter* sp. | UV13 | environmental, Antarctic | MHA | yes |
| G- bacteria | Pseudomonadota (Alphaproteobacteria) | *Sphingomonas* sp. | UVA17_B | environmental, Antarctic | R2A | yes |
| | | *Sphingorhabdus* sp. | UVA30_5 | environmental, Antarctic | R2A | yes |
| | | *Methylobacterium variabile* | UVA24 | environmental, Antarctic | R2A | yes |
| | | *Pararhizobium sp.* | UVA25 | environmental, Antarctic | MHA | yes |
| | Pseudomonadota (Betaproteobacteria) | *Masillia sp.* | UV278 | environmental, Antarctic | R2A | yes |
| | | *Burkholderia cenocepacia* | H111 | clinical | MHA | yes |
| | Pseudomonadota (Gammaproteobacteria) | *Pseudomonas* sp. | UV261 | environmental, Antarctic | MHA | - |
| | | *Escherichia coli* | 328 | clinical | MHA | - |
| | | *Klebsiella pneumoniae* | 927 | clinical | MHA | - |
| | | *Pseudomonas aeruginosa* | 25.211 | clinical | MHA | - |
| | | *Acinetobacter baumanii* | JCH 10.26.3 | clinical | MHA | - |
| Yeasts | Ascomycota | *Candida albicans* | ICA1 | clinical | MHA + 2% glucose + MB | yes |
| | | *Nakaseomyces glabratus* | 6448 | clinical | MHA + 2% glucose + MB | yes |
| | | *Nakaseomyces glabratus* | 8874 | clinical | MHA + 2% glucose + MB | yes |
| | | *Saccharomyces cerevisiae* | 5654 | clinical | MHA + 2% glucose + MB | - |
| | | *Saprochaete clavata* | 5788 | clinical | MHA + 2% glucose + MB | - |

R2A, Reasoner´s 2 agar; MHA, Mueller-Hinton agar; GPHF, glucose-peptone-yeast-beef medium; MB, methylene blue

**Supplementary Table S8**. Minimum inhibitory concentration (MIC) and Minimum fungicidal concentration (MFC) values recorded for tested yeasts.

| Strain | **Disk diffusion assays (mm)** | **MIC (mg/mL)** | **MFC (mg/mL)** |
|---|---|---|---|
| *Micrococcus luteus* CCM 169[T] | 40 [a] | - | - |
| *Saccharomyces cerevisiae* 5654 | 24 [b] | 2.5 | 5.0 |
| *Nakaseomyces glabratus* 6448 | 29 [b] | 0.5 | 2.5 |
| *Nakaseomyces glabratus* 8874 | 30 [b] | 0.5 | 2.5 |

[a]50ul of a 10mg/ml extract; 100ul of a 10mg/ml extract

**Supplementary Table S9**. Results of screening antiproliferative assays with pre-purified kineochelins extract.

| **Type of cells** | **Cell line** | **Cancer type** | **Activity ($IC_{50}$) - E1** | **Activity ($IC_{50}$) - E2** |
|---|---|---|---|---|
| Cancer cells | A549 | human alveolar adenocarcinoma | 544 ± 31 | 714 ± 15 |
| | U-87 MG | human glioblastoma | 773 ± 53 | 742 ± 27 |
| | PaTu 8902 | human pancreatic adenocarcinoma | 783 ± 55 | 1706 ± 96 |
| | Jurkat | leukaemia | 857 ± 144 | 1051 ± 140 |
| | HCT116 | human colorectal carcinoma | 899 ± 68 | 494 ± 48 |
| | A2058 | human metastatic melanoma | 639 ± 32 | 829 ± 57 |
| | MDA-MB-231 | human mammary gland adenocarcinoma | 1611 ± 150 | 1185 ± 125 |
| Non-cancer cells | CCD-18Co | human colon fibroblasts | 8247 ± 874 | 1881 ± 580 |

**Supplementary Table S10.** Information on genome assembly from *Actinokineospora* sp. UV203

| | Genome attribute | UV203 |
|---|---|---|
| Genome characteristics | Genome size (bp) | 6474610 |
| | DNA G+C content (%) | 69.4 |
| | Protein-coding genes | 6,112 |
| | Pseudo genes | 26 |
| | rRNA genes (5S, 16S, 23S | 3, 3, 3 |
| | tRNA genes | 73 |
| | ncRNA genes | 10 |
| | CRISPR | 0 |
| | oriC/oriV | 0 |
| | oriT | 0 |
| Genome quality | Sequencing technology | Oxford Nanopore Technologies |
| | Number of contigs | 2 |
| | Largest contig | 6462755 |
| | Total length | 6474610 |
| | Completeness | 100 |
| | Contamination | 1.78 |
| | Heterogeneity | 0 |
| | N50 | 6462755 |

**Supplementary Table S11.** Clinical yeast strains used in this study and their antifungal susceptibility determined by minimum inhibitory concentrations according to EUCAST breakpoints.

| Yeast strain | Source of isolation | Antifungal susceptibility | | | | | | | |
|---|---|---|---|---|---|---|---|---|---|
| | | **AMB** | **CAS** | **MIC** | **AND** | **VOR** | **POS** | **FLU** | **ITR** |
| *Saccharomyces cerevisiae* 5654 | Invasive infection, blood | S | S | S | S | R | NB | I | I |
| *Saprochaete clavata* 5788 | Invasive infection, blood | S | R | R | R | I | I | R | NB |
| *Nakaseomyces glabrata* 6448 | Invasive infection, blood | S | S | S | S | R | R | R | R |
| *Nakaseomyces glabrata* 8874 | Invasive infection, blood | S | I | S | S | S | R | I | R |
| *Candida albicans* Ica | urinary tract infection, urine | S | S | S | S | S | S | S | S |

S, sensitive; R, resistant; NB, no breakpoint available; I, intermediate; AMB, amphotericin B; CAS, caspofungin; MIC, micafungin; AND, anidulafungin; VOR, voriconazole; POS, posaconazole; FLU, fluconazole; ITR, itraconazole

**Supplementary Table S12.** List of all primers used in this study.

| ID | Sequence 5´→3´ | Origin |
|---|---|---|
| **1492r** | GGTTACCTTGTTACGACTT | Kane et al., 1993[4] |
| **616v** | AGAGTTTGATYMTGGCTC | Juretschko et al., 1998[5] |
| **aprR_Fwd** | GCAACAGTGCCGTTGATCGTGC | This study |
| **aprR_Rev** | TGCCCCTCCAACGTCATCTCGT | This study |
| **LuxR_Fwd** | GCAT**GGATCC**GTAGAATCCGGTTCCAGAGGTG* | This study |
| **LuxR_Rev** | GCAT**GAATTC**ACATCGTGGCGGTGAACGATC | This study |
| **Reg2_Fwd** | GCAT**GCGGCCGC**GTCCGCCAGAACCCGTTG | This study |
| **Reg2_rev** | GCAT**GAATTC**TGATGCTGCGCCTGCGCTAC | This study |
| **UV203_rrnP_Fw** | ACTGTCTA**GAATTC**GGATGTGCGTGTGTTGT | This study |
| **UV203_rrnP_Rev1** | GACA**GGATCCC**CAGCGTTCGTCCTGAGC | This study |
| **UV203_rrnP_Rev2** | GACA**GCGGCCGC**CCAGCGTTCGTCCTGAGC | This study |
| **2475_F_EcoRI** | GACA**GAATTC**CCAGGAACTCCCATTCCAC | This study |
| **2475_R_HindIII** | AGTG**AAGCTT**TAGCTCAGGGATTCGCCC | This study |
| **2475_R_HindIII_B** | ACTG**AAGCTT**GGCGTCGCCTGCATGAC | This study |
| **2479_F_EcoRI** | CTCA**GAATTC**GCGCACCAGCAGGTCA | This study |
| **2479_R_HindIII** | AGTC**AAGCTT**CCAGCCCTTGTCCAGCA | This study |
| **2479_R_HindIII_C** | ACTG**AAGCTT**GCAGTTGACCATGCCCG | This study |
| **QC1_2475_F** | CCTGTCCTACGCCGACTACT | This study |
| **QC2_2475_R** | CGCTGCTGGTGATCAGGTC | This study |
| **QC1_2479_F** | TGGACTACCTGAGCTGGCAG | This study |
| **QC2_2479_R** | CCGACCCGATGCCAAATGTATCG | This study |

(*) Restriction sites are highlighted in bold

**Supplementary Table S13.** List of all strains and vectors used in cloning part of this study

| | ID | Description | Origin |
|---|---|---|---|
| **Bacterial strain** | UV203 | *Actinokineospora* sp. UV203, wild-type | This study |
| | UV203_TC10 | Mutant, harbouring empty vector pSET152 | This study |
| | UV203_TC3_9 | Mutant, harbouring pSET152-ermE*p + LuxR | This study |
| | UV203_TC4_2 | Mutant, harbouring pSET152-ermE*p + Reg2 | This study |
| | UV203_T1A | Mutant, harbouring pSET152- rrn*p + LuxR | This study |
| | UV203_T2B | Mutant, harbouring pSET152- rrn*p + Reg2 | This study |
| | *E.coli* DH5α | general cloning host | New England Biolabs |
| | *E.coli* ET12567 (pUZ8002) | strain for intergenic conjugation; $Km^R$, $Cm^R$ | Flett et al., 1997[6] |
| **Plasmid** | pSOK201 | pSG5 minimal replicon, $Am^R$, *RP4 oriT*, *ColEI* replication origin | Zotchev et al., 2000[7] |
| | pSET152 | integrative ϕC31-based vector, $Am^R$ | Bierman et al., 1992[8] |
| | pSET152 + ermE*p | integrative ϕC31-based vector, $Am^R$, ermE*p | Sioud et al., 2002[9] |
| | pSET152-ActLuxR | pSET152 + ermE*p and LuxR from UV203 | This study |
| | pSET152-ActReg2 | pSET152 + ermE*p and LuxR from UV203 | This study |
| | pSET152-rrnp/LuxR | pSET152 + rrn*p and LuxR from UV203 | This study |
| | pSET152-rrnp/Reg2 | pSET152 + rrn*p and Reg2 from UV203 | This study |
| | pSOK201-Δ2475(600) | pSOK201 with 600bp fragment from core gene ctg2_2475 | This study |
| | pSOK201-Δ2475(1000) | pSOK201 with 1,00bp fragment from core gene ctg2_2475 | This study |
| | pSOK201-Δ2479(600) | pSOK201 with 600bp fragment from core gene ctg2_2475 | This study |
| | pSOK201-Δ2479(1000) | pSOK201 with 1,000bp fragment from core gene ctg2_2475 | This study |

$Am^R$, apramycine resistance; $Km^R$, kanamycin resistance; $Cm^R$, chloramphenicol resistance
